# Multi-Niche Human Bone Marrow On-A-Chip for Studying the Interactions of Adoptive CAR-T Cell Therapies with Multiple Myeloma

**DOI:** 10.1101/2024.04.08.588601

**Authors:** Delta Ghoshal, Ingrid Petersen, Rachel Ringquist, Liana Kramer, Eshant Bhatia, Thomas Hu, Ariane Richard, Reda Park, Jenna Corbin, Savi Agarwal, Abel Thomas, Sebastian Ramirez, Jacob Tharayil, Emma Downey, Frank Ketchum, Abigail Ochal, Neha Sonthi, Sagar Lonial, James N. Kochenderfer, Reginald Tran, Mandy Zhu, Wilbur A. Lam, Ahmet F. Coskun, Krishnendu Roy

**Author notes:** Corresponding author’s.

## Abstract

Multiple myeloma (MM), a cancer of bone marrow plasma cells, is the second-most common hematological malignancy. However, despite immunotherapies like chimeric antigen receptor (CAR)-T cells, relapse is nearly universal. The bone marrow (BM) microenvironment influences how MM cells survive, proliferate, and resist treatment. Yet, it is unclear which BM niches give rise to MM pathophysiology. Here, we present a 3D microvascularized culture system, which models the endosteal and perivascular bone marrow niches, allowing us to study MM-stroma interactions in the BM niche and model responses to therapeutic CAR-T cells. We demonstrated the prolonged survival of cell line-based and patient-derived multiple myeloma cells within our *in vitro* system and successfully flowed in donor-matched CAR-T cells. We then measured T cell survival, differentiation, and cytotoxicity against MM cells using a variety of analysis techniques. Our MM-on-a-chip system could elucidate the role of the BM microenvironment in MM survival and therapeutic evasion and inform the rational design of next-generation therapeutics.

**TEASER:** A multiple myeloma model can study why the disease is still challenging to treat despite options that work well in other cancers.

## INTRODUCTION

Multiple myeloma (MM) is the second-most common type of hematological cancer, affecting antibody-secreting plasma cells. MM cells accumulate in the bone marrow, interfering with normal hematopoiesis and immune functions. MM cells also interact with various components of the bone marrow microenvironment, which provide them with survival signals, growth factors, and protection from chemotherapy [1]. Chimeric antigen receptor (CAR) T cells have shown some promise in treating the disease, first gaining FDA approval in 2021. However, they still have not proven to be a cure, potentially due to their lack of persistence after infusion. This is in contrast to their efficacy in previous indications like leukemia and lymphoma [2–6].

The human bone marrow is a complex and dynamic tissue that is the primary site of ongoing hematopoiesis. The bone marrow consists of various cell types, both immune and stromal. BM-resident immune cells play essential roles in immune surveillance, defense against pathogens, and tolerance induction. The stromal cells in the bone marrow are non-hematopoietic cells that provide structural and functional support to the hematopoietic cells. They include mesenchymal stromal cells (MSCs), endothelial cells, and osteoblasts. These cells secrete ECM components, such as collagen and fibronectin, that form a three-dimensional network that anchors and regulates the behavior of hematopoietic cells. The ECM also contains growth factors, chemokines, and adhesion molecules that interact with BM-resident cells and influence cell proliferation, differentiation, migration, and survival [7].

The bone marrow contains two major niches: endosteal and perivascular. The endosteal niche is located at the bone and bone marrow interface. It is composed of osteoblasts, osteoclasts, and endosteal-like ECM components, such as osteopontin and type I collagen. The perivascular niche is located around the blood vessels in the bone marrow and comprises endothelial cells and supportive pericytes. The endosteal and perivascular niches are not mutually exclusive but cooperate and communicate to maintain dynamic equilibrium [8].

MM cells can exploit both niches to gain their survival and growth advantages. MM cells can adhere to osteoblasts and osteoclasts in the endosteal niche through integrins and ECM receptors, such as CD44. This interaction activates signaling pathways in MM cells, such as NF-κB and Wnt, that promote their proliferation, drug resistance, and osteolytic activity [9]. MM cells can also migrate to the perivascular niche and associate with endothelial cells and perivascular MSCs. This interaction stimulates angiogenesis and vasculogenesis in the bone marrow, enhancing nutrient and oxygen delivery to MM cells [10].

Moreover, MM cells can induce changes in the bone marrow microenvironment that favor their growth and evade immune surveillance. MM cells can also modulate the immune system by suppressing T cells and natural killer cells and enhancing regulatory T cells and myeloid-derived suppressor cells. Additionally, MM cells can alter the ECM composition and function in the bone marrow by reducing the levels of ECM proteins, such as fibronectin, collagen IV, and osteopontin, and by increasing the expression of proteases, such as matrix metalloproteinases (MMPs) and cathepsins [11, 12]. These changes affect the adhesion and migration of MM cells and other hematopoietic cells.

Many groups have attempted to model the bone marrow microenvironment, focusing on recapitulating one *in vivo* aspect or another. These models have used ECM and protein scaffolds [13–15] or bulk gels [16–21] to study the bone marrow and how its interactions with MM cells grant some degree of drug resistance. While these approaches involve multiple niches, they do not incorporate microvascular networks or the ability to study the BM microenvironment in standard imaging and plate reading equipment. Other modeling approaches utilized animal implantation before *in vitro* culture [22–25], which is costly and limited by the difficulties associated with correlating *in vivo* mouse findings to human physiology.

Microfluidic devices and tissue-on-chip models are another tool proposed to bridge the gap between 2D cell culture and animal models due to their ability to simulate complex microenvironments on a smaller scale [26–29]. These models have been used previously to simulate the bone marrow microenvironment with stromal cells and endothelial cells [30], including by our group [31]. These systems are desirable for studying external stimuli, such as therapeutic drugs or cells, within a complex system and for studying the 3D organization of the tumor microenvironment [29, 32–38].

A functional model of human MM enables detailed studies of its pathological effects in a small-scale, tunable culture system. Limitations of animal models, especially concerning precisely controlled manipulations and dynamic evaluation inside the marrow, have limited our ability to reach a consensus regarding the roles and character of the niche environments, their cross-talk, specific cell types, and signaling pathways.

Understanding the pathophysiology of MM in a physiologically relevant human model that utilizes primary patient samples to ensure diversity within the BM microenvironment may also potentially lead to a better understanding of CAR-T cell toxicity, especially how stromal and immune cells in the MM microenvironment interact with and respond to the infused cells. This could aid us in enhancing cell migration and retention within the MM niche, increasing memory T cell development and proliferation, and enhancing response durability through increased persistence and decreased T cell attrition. To that end, we created a 3D microvascularized culture system to model the spatially-distinct endosteal and perivascular bone marrow niches, allowing us to study MM-stroma interactions and model responses to therapeutic CAR-T cells that were flowed into the perfusable microvasculature.

## RESULTS

Please note that, for the entirety of this article, the term *plate* will be used to delineate a single 96wp that contains a 4×2 array of hBM-on-a-chip/hMM-on-a-chip devices (**Figure 1A**), whereas a *device* refers to a single unit within an 8-device plate (**Figure 1B**).

**Figure 1:**
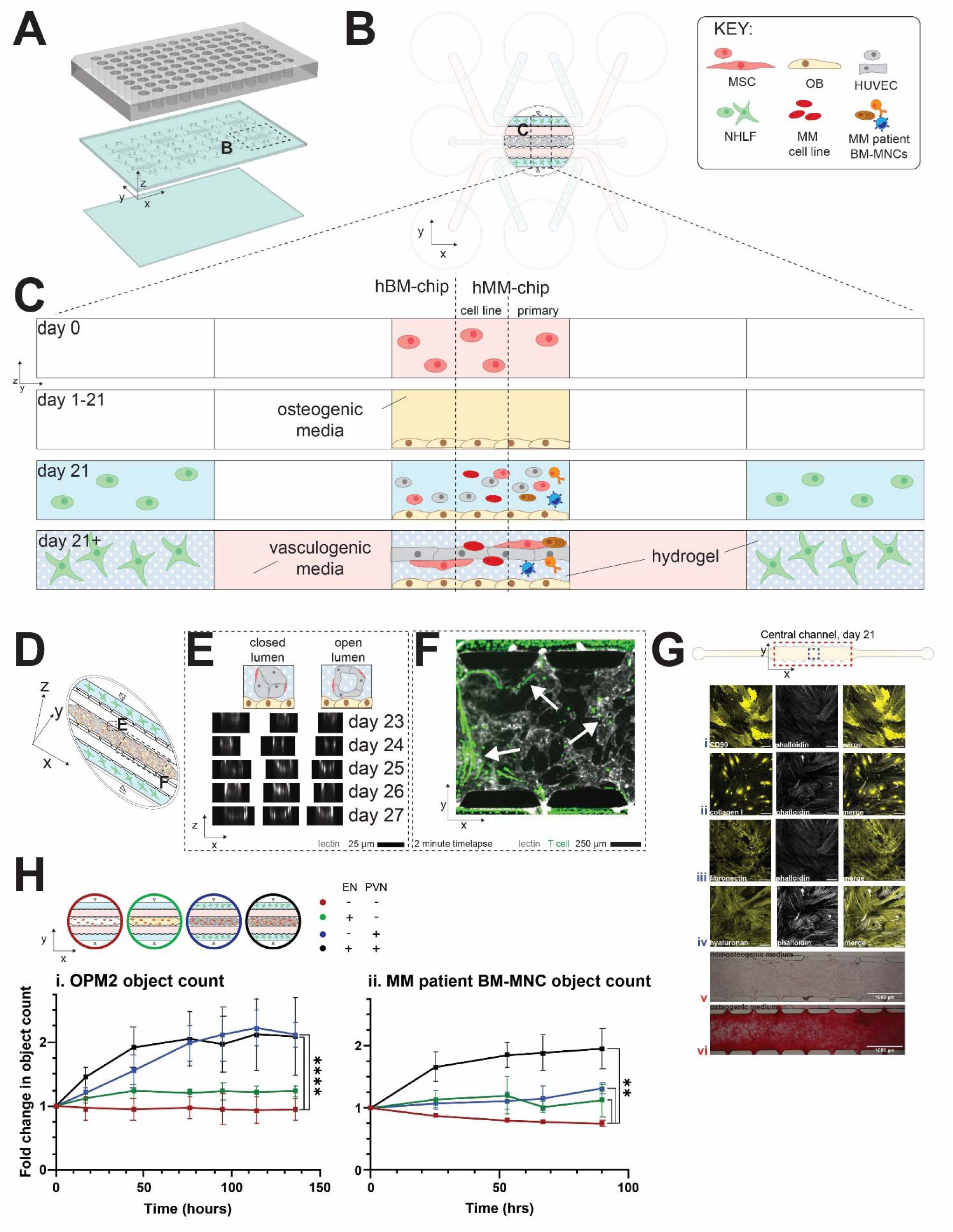
Overview of hBM-on-a-chip and hMM-on-a-chip. (A) macroscopic exploded view of a single plate that contains 8 devices. Dashed box is for (B) inset of a single device’s top view (XY). Dashed box is for (C) inset of 5-channel side view (YZ) over the culture period. Note that the central channel can contain either the BM stroma alone (left) or BM stroma plus MM cells, either in the form of an MM cell line (center) or primary BM mononuclear cells from an MM patient (right). (D) Rotated view of the 5 channels of a single device, with dashed boxes for insets in (E) and (F). (E) Cartoon depiction of immature, closed lumens compared to matured, donut-shaped lumens (top). Over the culture period, the lumens of the vessel network open, as shown by the XZ view of lectin-labeled endothelium (bottom). (F) This enabled the perfusion of cells that were introduced via the media channels, as shown by the superimposed images from a 2-minute timelapse. As seen by the white arrows, perfused cells followed along the paths of lectin-labeled vessels. (G) By the end of the 21-day osteogenesis induction phase, the central channel expressed hallmark endosteal and MSC markers, such as (i) CD90, (ii) Type I collagen, (iii) fibronectin, and (iv) hyaluronan. Extended focus confocal of representative devices. Scale bars = 100 µm. The MSC-derived osteoblasts also mineralized the channel in response to the osteogenic factors supplemented within the medium, as evidenced by alizarin red staining of devices fed with (v) non-osteogenic medium and (vi) osteogenic medium.

### hMM-on-a-chip design enabled a spatially distinct culture of endosteal and perivascular niches

The human MM-on-a-chip devices were comprised of a 5-channel microfluidic channel set, with specific dimensions described previously [31], with 8 individually addressable devices per 96-well plate (**Figure 1A**). Inlet and outlet ports were punched into each channel of each device to enable liquid and hydrogel loading. There were 2 outer channels for holding pro-vasculogenic factor-secreting fibroblasts. These two channels were separated from the central channel by two media channels (**Figure 1B**). A graphical summary of device culture may be found in **Figure 1C**. Briefly, human bone-marrow-derived primary MSCs were loaded into the central channel on day 0. These were induced to differentiate into OB over the next 21 days by adding pro-osteogenic factors to the MSC culture medium within the central channel to form the endosteal niche (EN). After this phase of culture was complete, hydrogel containing the perivascular niche (PVN) was loaded atop this MSC monolayer containing HUVECs, pericytes (more MSCs), and any additional cell types of interest, like MM cell lines or bone marrow mononuclear cells from MM patients (BM-MNCs). In parallel, the outer channels were loaded with normal human lung fibroblasts (NHLFs). In this way, the cells of the endosteal niche could be cultured on a different z-plane than the cells in the hydrogel-encapsulated perivascular niche.

### hMM-on-a-chip perivascular niche contained perfusable vascular networks

Over the following days of culture, the outer channel NHLFs secreted pro-vasculogenic factors into the media of the media channels, which is also supplemented with the pro-vasculogenic cytokines VEGF and Ang-1. This resulted in a network of endothelial cells, supported by pericytes, forming in each device. This network was visualized in 3D using confocal and epifluorescence microscopy (**Figure 1C**).

By staining live devices with fluorescent *Lycopersicon esculentum* lectin, we could then visualize the lumens opening in response to pro-angiogenic signaling (the donut shapes in **Figure 1E**, bottom, as compared to the schematic in **Figure 1E**, top). These lumens faced outward through the spaces between pillars. They allowed the introduction of external cells to the system through the media channel, such as T cells, for downstream cytotoxicity studies (**Figure 1F**).

### Modeled endosteal niche expressed hallmark extracellular matrix and mineral components

As seen in **Figure 1G**, different hallmark ECM proteins and MSC surface markers were readily visualized through confocal microscopy. We were able to detect robust expression of CD90, type I collagen, fibronectin, and hyaluronan (**Figure 1G(i-iv)**), as well as osteopontin, osteocalcin, and stem cell factor (**Supplementary Figure 1B(i-iii)**), all as compared to secondary and unstained controls (**Supplementary Figure 1D**). As seen in **Figure 1G(v)**, there was very little red signal within the central channel after 21 days of culture within non-osteogenic medium. In contrast, after 21 days of culture with osteogenic medium, large amounts of mineralization were readily apparent, evenly staining the entire central channel of the device (**Figure 1G(vi)**). Multiple biological replicates were quantified, showing significant increases in red color brightfield signal between the two media conditions, as quantified by Welch’s nonparametric t-test (**Supplementary Figure 1C**).

### Modular hMM-on-a-chip design demonstrated need for inclusion of multiple BM niches

To convert the human bone marrow on-a-chip (hBM-on-a-chip) design[31] into the human multiple myeloma on-a-chip (hMM-on-a-chip), we cocultured either MM cell lines or primary bone marrow mononuclear cells from MM patients (BM-MNCs) within the devices. One of the benefits of the hBM-on-a-chip system was its modularity; any number of cell types could be added or omitted from the device’s culture. For example, as portrayed in the **Figure 1H**, top, devices could contain only hydrogel-encapsulated MM cells (red), hydrogel-encapsulated MM cells atop the MSC-derived OB monolayer (green), hydrogel-encapsulated MM alongside HUVECs and MSCs without the OB monolayer below (blue), or finally, MM cells cultured within the full hBM-on-a-chip system (black).

We decided first to investigate the individual inputs of each niche on MM cell proliferation by systematically omitting them from devices. After daily image monitoring of devices to track the proliferation of labeled target cells, the proliferation of labeled MM objects significantly increased for devices that contained at least the PVN components by 2-way repeated measures ANOVA and Dunnett’s posthoc tests between the control group (MM only, red trace) and the other groups (**Figure 1H**, bottom). Stars show the statistical significance of the final time point.

### Fluorescent MM cell lines proliferated within the hMM-on-a-chip environment

After successfully creating a microvascularized model of human bone marrow that recapitulated key markers of both the endosteal and perivascular niches, we next cultured several fluorescently labeled MM cell lines.

All transduced cell lines retained their wild-type surface expression profiles of BCMA+CD138+CD45-CD19- (BCMA MFI data shown in **Supplementary Figure 2**) while expressing RFP clearly. Furthermore, these fluorescent cell lines could readily be tracked visually using fluorescent microscopy within the hMM-on-a-chip (**Figure 2B**, **Supplementary Figure 3D**). Indeed, over the timeline of culture (**Figure 2A**), the number of objects increased over time for multiple fluorescent tested cell lines and stayed stable for the others (**Figure 2B**). Meanwhile, the average RFP MFI (**Supplementary Figure 3A**) for all groups (except OPM2) did not decrease, confirming the stable expression of RFP within daughter cells. Although we mainly reported fluorescent object count throughout this article, we also measured object size (**Supplementary Figure 3B**) and total area (**Supplementary Figure 3C**) in order to account for clumped, dividing cells. We have observed a stable culture of MM for as long as 12.5 days, but chose endpoint devices after about 7 days for most experiments to retain stable vessel networks throughout (**Supplementary Figure 4**).

**Figure 2:**
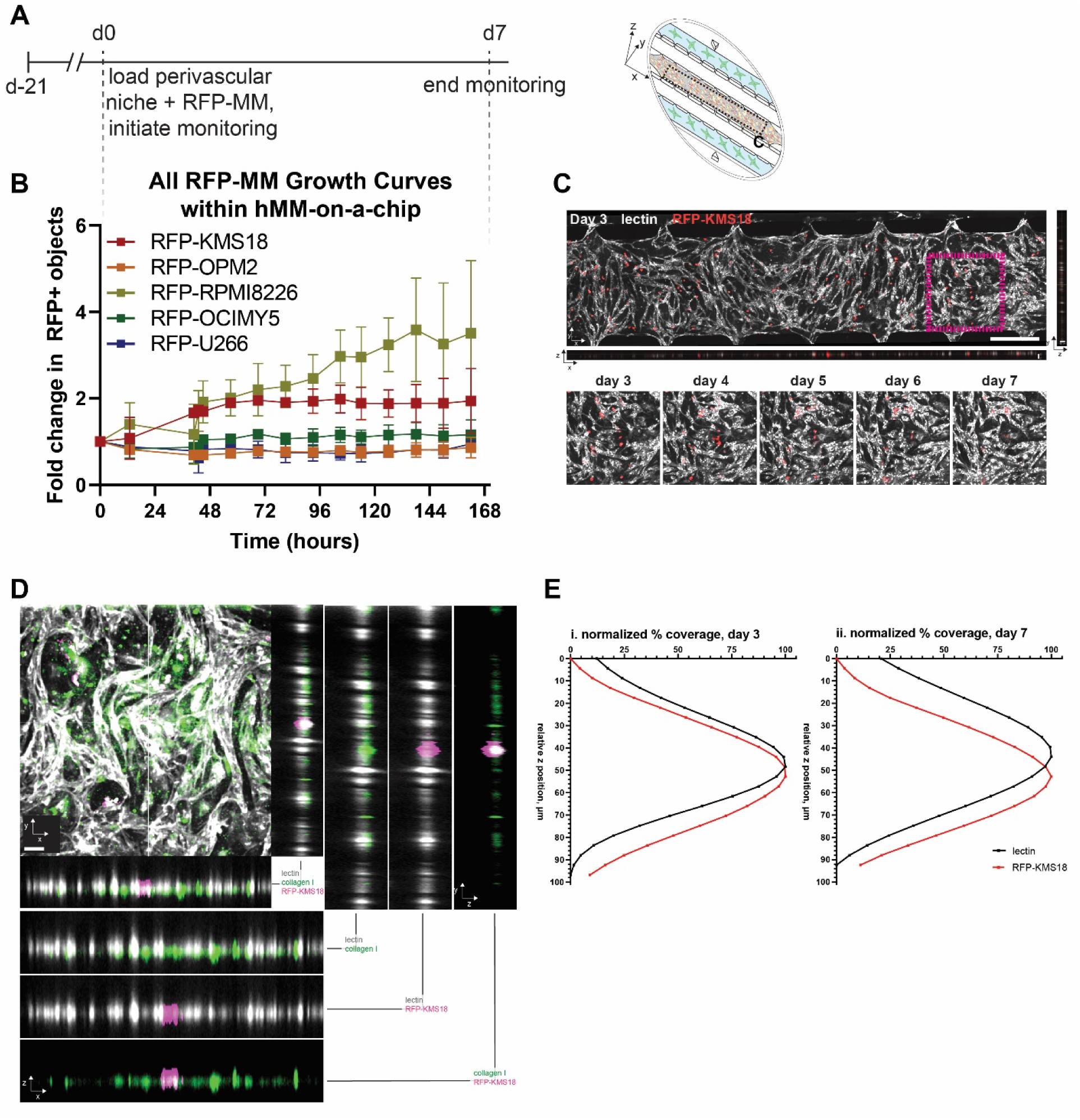
Localization of fluorescent MM cell lines within in vitro BM niche. (A) Timeline of device culture, alongside a 3D device schematic, to orient the reader. Dashed line gives viewing plane of panel (C). (B) Fluorescent timecourse proliferation of 5 different cell lines. n = 8-10 individual devices across 3 experimental sets for each cell line. (C) 3D views of a single confocal z stack for RFP-KMS18 on day 3, showing qualitatively the localization of the red-fluorescent MM cell line (shown in red) to the LEL-labeled vascular network (shown in white). The XZ and YZ views confirm that both signals are in the same z-planes, and inset views of the magenta box show the same ROI over the 7-day culture period. (D) The positive association of RFP-KMS18 to the perivascular niche is shown by counterstaining devices at the culture endpoint for the endosteal-associated collagen I. The RFP-KMS18 (magenta) localized in the same z-planes as lectin (white), which were different z-planes than collagen I (green). Different combinations of the 3 fluorophores are shown in XZ and YZ for ease of interpretation. Scale bar = 50 µm. (E) This perivascular localization was also quantified by % coverage of lectin and RFP signal in every relative z-position in (C) for (i) Day 3 and (ii) Day 7 for RFP-KMS18, respectively.

### MM cell lines localized to the perivascular niche of the hMM-on-a-chip

We already observed in **Figure 1H(i)** that the OPM2 cell line seemed to proliferate best in the presence of the perivascular niche of the hMM-on-a-chip, and we had confirmed this using cyclic immunofluorescence to interrogate multiple niche markers from the endosteal and perivascular niches and their coexpression with OPM2 (**Supplementary Figure 6**). We next investigated whether that meant that the MM cells also strongly localized to the PVN. Using the same daily confocal imaging used to study the open lumens in **Figure 1E**, we studied the relative abundance of the RFP-KMS18 signal in z, as compared to the abundance of the LEL-tagged vasculature and qualitatively observed that the MM cell lines seemed to localize strongly to the lectin-labeled vasculature (**Figure 2C**, **Supplementary Figure 3D**), as evidenced by all 3 3D planes of view (please reference **Figure 2A** for an orientation to the axes). We also demonstrated that MM cell lines localized to a z-plane above the endosteal niche by counterstaining for Type I collagen (**Figure 2D**, **Supplementary Figure 5**). Finally, we quantified the percent coverage of RFP and LEL signals from the 3D confocal images for the 3^rd^ and 7^th^ days of culture (the first day after LEL addition and the last day of culture), and confirmed that the red signal seemed to peak at or above the peak of the LEL, further supporting that the two signals were colocalized in z (**Figure 2E**), within the theoretical 11 µm z-resolution of the microscope setup.

### Primary plasma cells survived within hMM-on-a-chip over the time period of culture

To that end, we cultured patient BM-MNCs within the system and digested devices at periodic time points. By using spectral flow cytometry, we found that, as seen in **Supplementary Figure 7**, the starting population of patient BM-MNCs was diverse, comprising of RBCs and T cells, as well as plasma cells and other less well-defined immune and stromal populations, as characterized by their expression of markers such as CD3, CD38, and CD45. When cultured within the device, however, the main cell types that survived long-term were the plasma cells and a very small subset of T cells (**Figure 3A(i-iii)**). In fact, the main cell types detectable in the UMAP analysis were CD31+ HUVECs and CD90+ MSCs, which are the only cell types found within the central channel of the human BM-on-a-chip model (**Figure 3A(iv-vi)**). However, it is certainly possible that bone marrow-resident EPC and other stromal cells also clustered within these populations for the hMM-on-a-chip digestions. After 4 days of culture, most of the less well-defined populations disappeared, leaving behind the main stromal populations and small plasma and T cell populations.

**Figure 3:**
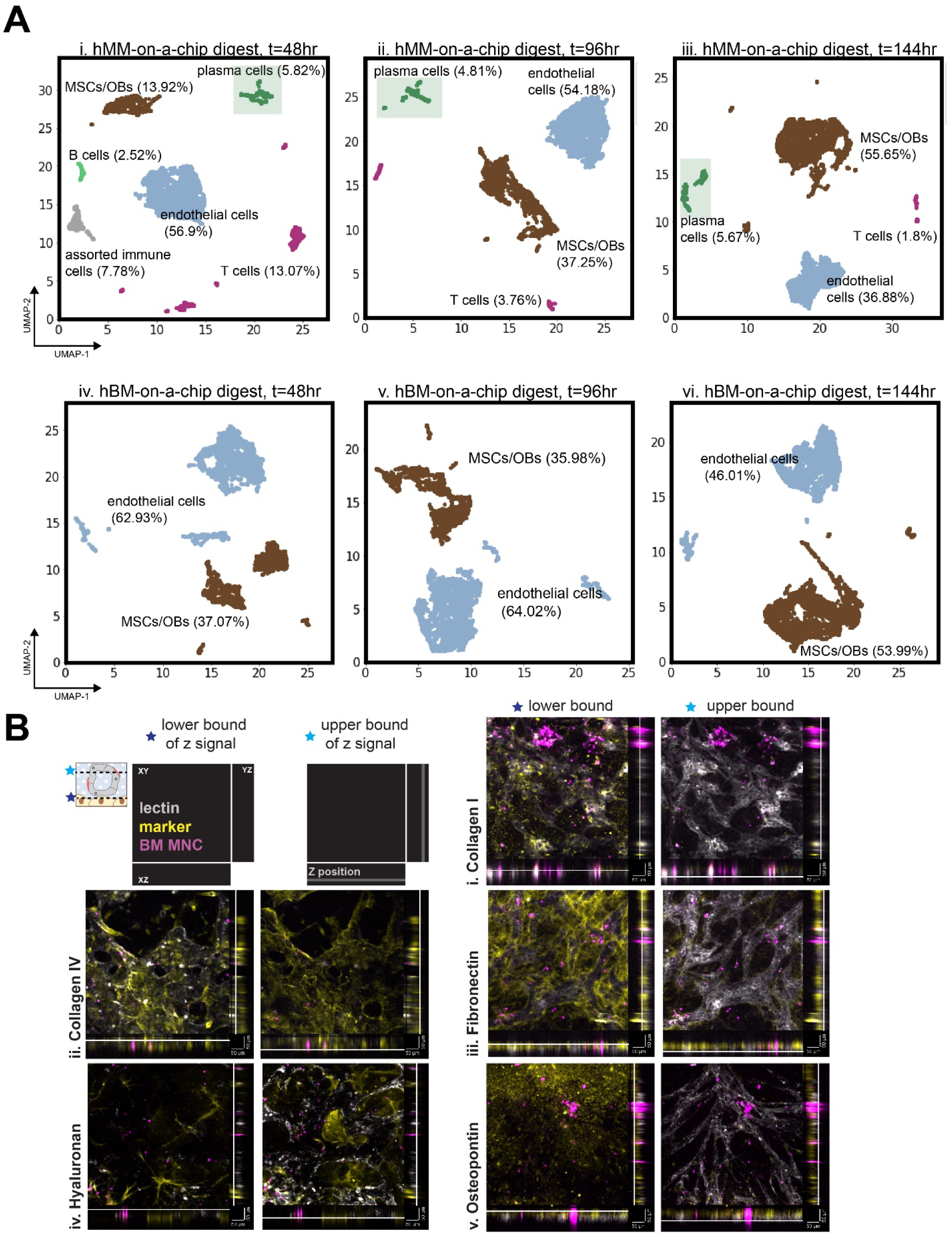
Diversity of primary cell populations and ECM spatial expression within hMM-on-a-chip. (A) The human MM-on-a-chip (i-iii) and BM-on-a-chip (iv-vi) could be digested to harvest its cells for flow cytometric analysis. UMAP could be performed at several timepoints in order to cluster the cells into different subpopulations based on their surface marker expression (labeled on each plot alongside their population frequency with the plasma cell cluster highlighted). This could be tracked over different timepoints, like (i, iv) 2 days, (ii, v) 4 days, or (iii, vi) 6 days following perivascular niche hydrogel loading. (B) Representative Z slices at the (left) bottom and (right) top of each z-stack to measure ECM spatial localization. In the top left, there is a schematic at the top left showing how the XY, XZ, and YZ views are displayed at multiple z-positions, with a line pointing to the active Z-slice. For the endosteal markers on the right column of (i) collagen I, (iii) fibronectin, and (v) osteopontin, the signal (yellow) was found strongly in the bottom half of the z-stack, but not at the top where the lectin-labeled vessels could be seen. In contrast, in the left column, (iv) hyaluronan could be found throughout, expressed in an MSC-hallmark spindle shape, and (ii) collagen IV was found at the same spatial locations as the endothelial network throughout. Scalebars = 50 µm.

### Extracellular matrix expression within the hMM-on-a-chip

We hypothesized that the observed prolonged survival of primary plasma cells could be due to the primary cells interacting with the extracellular matrix components of the device, as well as stroma. Therefore, we decided to perform 3D confocal microscopy to interrogate the expression of a variety of ECM markers within the device. **Supplementary Figure 8A** contains extended-focus displays of the results and unstained controls.

We looked at the signal at the bottom (right) and top (left) bounds of each z-stack’s discernible signal to confirm, for example, that type I collagen and osteopontin were primarily found at lower z-positions (**Figure 3B (i, v)**), type IV collagen was found wherever the lectin was (**Figure 3B (ii)**), and hyaluronan had its characteristic spindle morphology in both niches (**Figure 3B (iv)**). The fibronectin was mainly found in lower z-positions, but it was also visible within pericytes closely-associated with the labeled vessels (**Figure 3B (iii)**). These images also demonstrated the abundance of the labeled patient BM-MNCs throughout the different niches. Unlike the cell lines, which seemingly only localized around the PVN, primary cells could be found both within vessels, in the central marrow space, and near the endosteal niche, and seemingly proliferated in multiple spots. This was very interesting and further supports the view that these diverse cells were localizing everywhere and interacting with multiple stromal and ECM-derived factors within the hMM-on-a-chip. **Supplementary Figure 8B** contains a summarized quantification of signal coverage for 3 devices per staining condition, with error bars indicating the standard deviation in percent coverage at each acquired z-slice.

### Target cells within hMM-on-a-chip reduced in number when cocultured with CAR-T cells

We evaluated the fluorescence intensity and count for the red-fluorescent target cells within the hMM-on-a-chip after effector T cell addition as a proxy for T cell-mediated cytotoxicity. The same T cells were also assayed against target cells in 2D (**Supplementary Figure 9**). As shown in **Figure 4B(i)**, the number of red-fluorescent objects decreased over time after addition of CAR-T cells, compared to the untransduced (UTD) T cell-treated devices. This difference was statistically significant, particularly at later time points (2-way repeated measures ANOVA with Sidak’s post-hoc test). Furthermore, the images that the BioSpa acquired for quantification could also be studied to qualitatively observe that the devices with CAR-T cells had fewer target cells by the endpoint of the study, and the only remaining RFP-KMS18 within the images was far away from an effector T cell. This was in direct contrast to devices with UTD-T cells; MM cells seemed to proliferate there and even target cells that were nearby a labeled T cell still appeared visible by the study’s endpoint (**Figure 4C**). Further, the fluorescence intensity of CTV+ CAR-T cells decreased significantly over time, compared to that of the UTD-T cells (**Figure 4B(ii)**, **Supplementary Figure 12B**). This suggested division of the CAR-T cells rather than photobleaching over prolonged imaging exposure.

**Figure 4:**
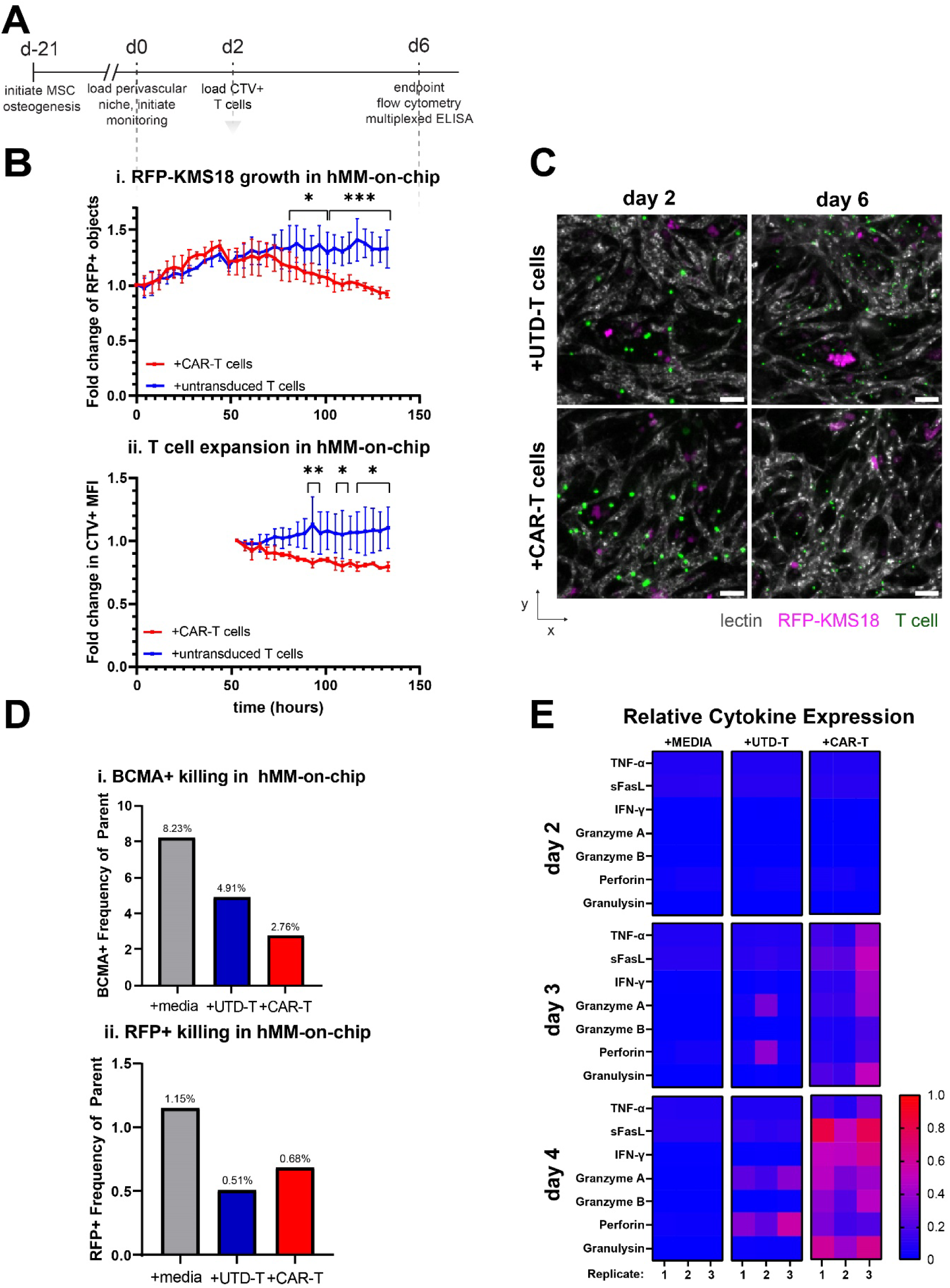
T cell cytotoxicity against target cells within hMM-on-a-chip. (A) Culture timeline for experiment. (B). (i) The number of RFP-KMS18 objects within hMM-chip devices decreased significantly with the addition of CAR-T cells compared to untransduced. (ii) In addition to RFP signal increasing over time, the MFI of the CellTrace Violet (CTV) signal from the labeled T cells decreased in the CAR-T cell group. This was not due to photobleaching over extended imaging because the untransduced cells did not reduce their signal, suggestive of T cell proliferation in response to target recognition. Statistics: 2-way repeated measures ANOVA and Sidak’s post-hoc tests for the CAR-T vs UTD-T groups. * = p < 0.05, *** = p < 0.001 for differences within any given timepoint. (C) Representative images of CellTrace Violet-labeled T cells (green) interacting with RFP-expressing KMS18 (magenta) and lectin-labeled vasculature (gray). In the case of CAR-T cells, the day 6 images demonstrated a reduction of red-fluorescent signal by the endpoint of culture. Scale bar =100 μm. (D) Digestion of hMM-on-chip at the endpoint of culture showed a decrease in BCMA+ cells in the final population, showing killing. Alternatively, the same trend could be seen with the RFP+ signal but with fewer cells in the population leading to some uncertainty. As devices were pooled together for these analyses, the data represents a summary of multiple biological replicates, so no statistics were run. (E) Heatmaps of key T cell-associated analytes at 3 different timepoints within the hMM-on-chip + media, untransduced T cells, or CAR-T cells, showing T cell activation within the CAR-T cell condition and some low-level background secretion by untransduced T cells. Each of the 9 heatmaps is an individual timepoint and effector treatment group. Each major row is a timepoint, with individual cytokines plotted. Each major column is a type of effector, and each subcolumn of cells is the cytokine release data over time for a single biological replicate (device) within the group.

As the most direct evidence for target cell killing, we digested the devices after culture with effectors, then measured the differences in expression of the CAR target BCMA (**Figure 4D(i)**) and the differences in RFP+ reporters (**Figure 4D(ii)**). As the devices were pooled together in order to yield enough cells for a reliable flow cytometry sample, statistical analyses could not be performed. The untreated group (gray) naturally had the highest percentage of remaining RFP+ cells. The UTD-T cell group (blue) and the CAR-T cell group (red) had the most reduction in RFP+ signals. In order to confirm the target-specificity of killing, we used the measurement of BCMA-expressing cells within the endpoint samples as a more direct metric of T cell effector function. Compared to untreated and UTD-T cells, CAR-T cells resulted in the largest decrease in BCMA expression in the overall sample. This suggested that CAR-T cells within the device recognized and activated against BCMA+ target cells. The reduction in UTD-treated target cells, as measured by both BCMA and RFP, made us suspect some form of graft-versus-host effect within the system.

### CAR-T cells released activation-associated cytokines within hMM-on-a-chip

The media was surveyed pre-T cell addition as well as 24 and 48 hours later. In **Figure 4E**, as with other metrics of observation, the CAR-T cell-treated devices had increases in T cell activation cytokines, displayed as normalized to their maximal expression level. The raw concentrations are reported in **Supplementary Figure 10**. Not every cytokine listed in **Supplementary Figure 9A** was detectable or upregulated here, and the raw concentrations of each cytokine were 55-16000-fold lower than those found in the 2D assay. This was not surprising because of the much lower effective E:T ratios found within the devices that had sparse target cell expression (**Figure 4D(i)**), notwithstanding the need for the T cells to travel to the target cells before exerting their functions. This, coupled with the BioSpa data from the same devices, supported the evidence for CAR-T cell-mediated target cell death within the hMM-on-a-chip.

### T cell activation cytokines were released from target- and graft-associated release pathways

We also observed some low-level release of T cell activation cytokines against the hBM-on-a-chip alone (**Supplementary Figure 11**), as well as some reduction in target cell burden within UTD-treated devices (**Figure 4B**), making it imperative to understand which cytokines were released as a result of graft-versus-host (GVH) effects *in vitro*. To that end, we added CAR- and UTD-T cells from either the same donor (“autologous”) or from a different patient (“allogeneic”) and sampled media periodically for analysis of cytokine release (**Figure 5A**). Not every cytokine listed in **Supplementary Figure 9** was detectable or upregulated here, and the raw concentrations (**Supplementary Figure 13**) of each cytokine were 55-300-fold lower than those found in the 2D assay as previously observed.

**Figure 5:**
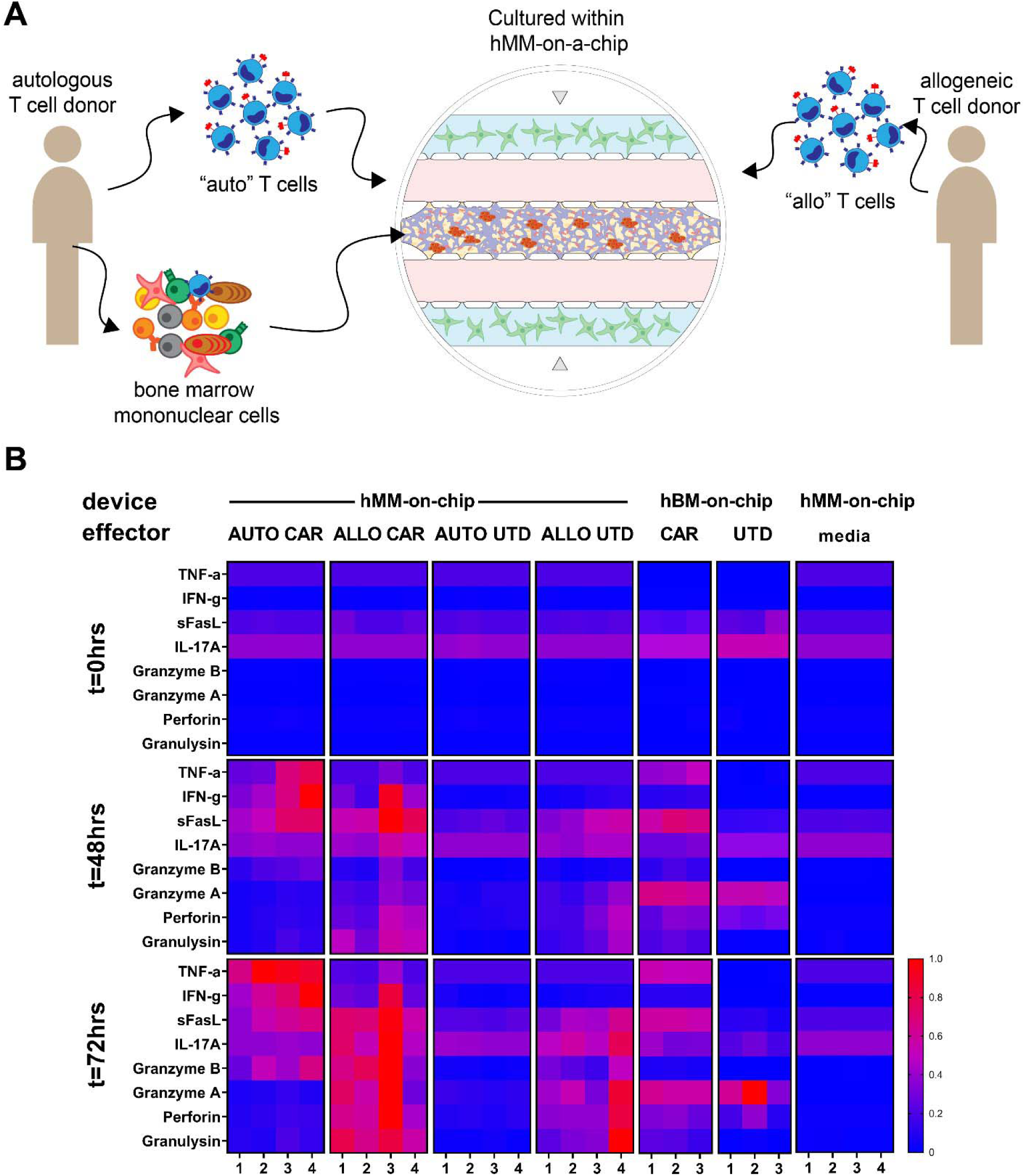
Graft-versus-host effects of CAR-T cells within the hMM-on-a-chip microenvironment. (A) Experimental schematic. (B) Heatmaps of the same cytokine release within treated hMM-on-a-chip devices. Each major row is a timepoint, with individual cytokines plotted. Each major column is a type of effector, and each subcolumn of cells is the cytokine release data over time for a single device within the group.

As shown in **Figure 5B**, allogeneic T cells did secrete activation-associated cytokines, regardless of whether they were CAR-T cells or untransduced. However, only specific cytokines were released in devices treated with autologous CAR-T cells, namely TNF-α and IFN-γ. Other cytokines like Granzyme A, perforin, and granulysin were upregulated in allogeneic-treated devices.

### Primary target cells shown to be susceptible to CAR-T cell cytotoxicity after distance filtering

Next, we wished to measure the proliferation of the PKH26-labeled primary BM-MNCs after the addition of CAR-T cells or UTD-T cells from the same donor, using the experimental setup described in **Figure 6A**. Since the plasma cells comprised only about 8% of the labeled population, and because dead cells did not lose their membrane label, we only quantified BM-MNCs that came within 60 μm of a T cell at some point during data acquisition (**Figure 6B**, unfiltered data in **Supplementary Figure 14A**). After application of this filter, it was apparent in **Figure 6C(i)** that primary BM-MNCs within 60 μm of a CAR-T cell did not proliferate as much as the cells within 60 μm of a UTD-T cell, suggesting that the CAR-T cells were able to control the proliferation somehow, potentially by BCMA-specific activation. **Supplementary Figure 14B, C** give the distribution of distances that gives justification for our choice to use a 60 µm cutoff. We also attempted to use real-time monitoring of caspase activity as a metric for cell death, but could not detect statistically-significant differences between groups, even with distance filtering (**Supplementary Figure 15**).

**Figure 6:**
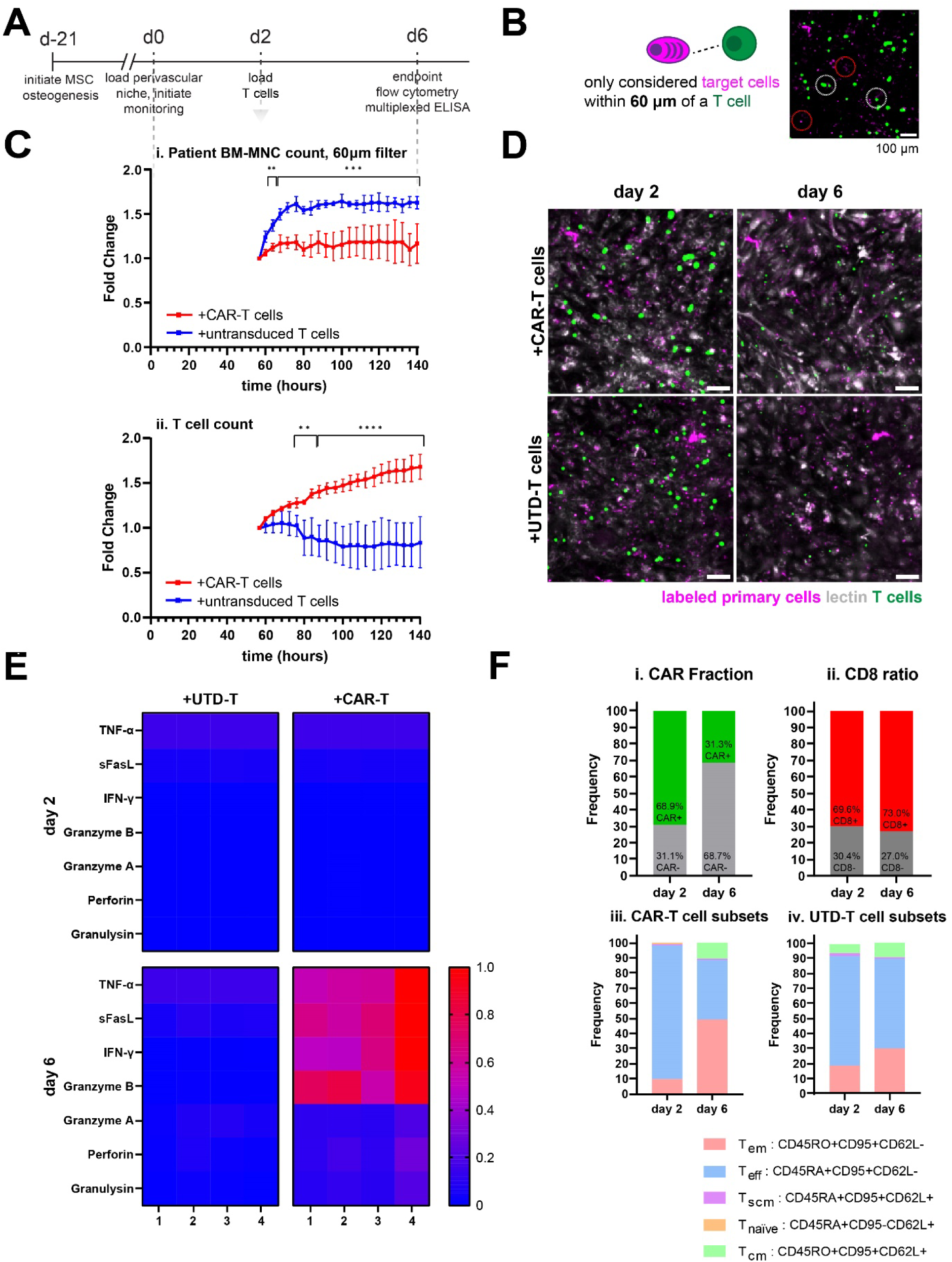
CAR-T mediated killing of primary BM-MNCs within hMM-on-a-chip. (A) Timeline of device culture, to orient the reader. Dashed line gives timescale of panel (C). (B) Schematic of Euclidean distance calculation for a single representative labeled target cell. (C) After distance filtration, the (i) resultant number of patient-derived BM-MNCs or (ii) CellTrace Violet-labeled T cells after introducing labeled CAR-T cells (red) or UTD-T cells (blue). n=4 devices per condition. 2-way repeated-measures ANOVA, with Sidak’s post-hoc test. ** = p < 0.01, *** = p < 0.001, **** = p < 0.0001 for differences within any given timepoint. (D) Representative images of T cell interaction with PKH26-labeled primary cells within lectin-labeled vessels, taken at the start and end of culture for the same ROI. Scale bar =100 μm. (E) Heatmaps of cytokine release within treated hMM-on-a-chip devices after 0 and 96 hours of treatment with effectors. Each of the heatmaps is an individual timepoint and effector treatment group. Each major row is a timepoint, with individual cytokines plotted. Each major column is a type of effector, and each subcolumn of cells is the cytokine release data over time for a single device within the group. (F) Flow cytometric characterization of patient T cells before (day 2) and after (day 6) hMM-on-a-chip culture, in terms of CAR fraction, CD8+ ratio, and memory phenotypes.

### CAR-T cells divided in response to primary target cells within hMM-on-a-chip

CAR-T cells increased in number within the hMM-on-a-chip system compared to UTD (**Figure 6C(ii)**); this effect was observed in both raw and distance-filtered data. We also evaluated the confocal images of each condition, to validate that T cells were indeed traveling through the lectin-labeled vessels. As seen in **Figure 6D**, the T cells were indeed colocalized to the LEL signal. When 20X confocal images were taken, T cells were confirmed to be localized near labeled primary cells. Furthermore, we observed visual evidence of the CAR-T cell proliferation in the quantification (**Supplementary Figure 16A**).

### CAR-T cells released activation-associated cytokines within hMM-on-a-chip, even with lower primary target cell burden

The media was surveyed pre-T cell addition as well as 96 hours later. As shown in **Figure 6E**, the CAR-T cell-treated devices had increases in T cell activation cytokines. The raw concentrations of each cytokine (**Supplementary Figure 17**) were 55-1000-fold lower than those found in the 2D assay (**Supplementary Figure 9**). This, coupled with the BioSpa data from the same devices, supported the evidence for CAR-T cell-mediated target cell death within the hMM-on-a-chip.

### CAR-T cell fraction, CD8 ratio, and memory populations were preserved within hMM-on-a-chip

We surveyed the CD3+ fraction of digested devices to see what types of T cell subsets survived within the devices. In order to avoid the loss of memory T cell subsets following thaw that we observed in the prior healthy-donor T cell experiments (**Supplementary Figure 12A**), we loaded freshly-expanded patient T cells without cryopreserving them. We also analyzed the subsets present within the devices after 7 days of culture. As seen in **Figure 6F**, the CAR fraction was retained over extended culture within the device, as was the ratio of CD8+/- T cells. Additionally, the memory phenotypes of the T cells within the sample reflected detectable fractions of central memory T cells (T_CM_s) and stem cell memory T cells (T_SCM_s) post-expansion. By the endpoint of device culture, the T_CM_ population seemed to be expanded within the devices, more so in CAR-T conditions, and the T_SCM_ fraction seemed to remain somewhat stable.

## DISCUSSION

This work aimed to develop a 3D model of human multiple myeloma compatible with a wide array of data acquisition techniques to better understand MM’s response to anti-cancer therapeutics, with a particular focus on therapeutic chimeric antigen receptor (CAR)-T cells.

First, we extended our human bone marrow on-a-chip (hBM-on-a-chip) model [31] into a model of human multiple myeloma. This well-plate-based design reduced device-to-device variability and facilitated the ease of automated data analysis using a variety of data acquisition modalities, such as timelapse imaging, flow cytometric analysis, and immunohistochemical staining (**Figure 1A-D**). Even following the introduction of primary or cell-line-based MM samples, the system retained hallmark characteristics of the bone marrow’s endosteal and perivascular niches, namely open lumens to enable transport of therapeutic agents like cells (**Figure 1E-F**) and expression of endosteal ECM, surface markers, and mineralization (**Figure 1G**) localized to the phalloidin of the MSCs as expected [17, 18, 39–45].

Next, we evaluated whether both niches were necessary to study MM proliferation. We confirmed that both cell line-based and primary MM samples only significantly proliferated at the final time point when cultured with niche components (**Figure 1H**), and plasma cells from primary patient-derived BM-MNC samples were preserved within the hMM-on-a-chip by flow cytometric analysis (**Figure 3A**). The perivascular niche seemed to have the most influence, which was unsurprising when considering the later investigations into MM’s preferred niche (**Figure 2C-E**, **Supplementary Figure 6**). Another potential explanation for this was the likely secretion of hydrogel-remodeling matrix metalloproteinases (MMPs) like MMP-2 and MMP9 by MM-associated ECs; these factors help endothelial cells degrade the hydrogel as they neovascularize, but may also have played a role in allowing MM cells to spread and proliferate [46, 47]. However, MM lines and diverse BM-MNCs from patients constitutively secrete MMPs on their own, so it is unlikely that hydrogel remodeling by endothelium was the only reason for increased growth [12].

We then studied cell-line-based multiple myeloma in the system and its localization to the perivascular niche (PVN) using genetically modified reporter versions of various multiple myeloma cell lines. We were able to monitor the PVN in real-time through the use of fluorescent-tagged *Lycopersicon esculentum* lectin (LEL). While others have used lectins to label vessel networks *in vivo* and *in vitro* in the past as an endpoint assay [48, 49], to our knowledge, this is the first time that LEL has been used to label live vascular networks *in situ* within a microfluidic device to study vessel development, maturation, and interaction with tumor or effector cells. We found that, while cell line MM localized strongly to the PVN (**Figure 2C-E**), primary MM cells were localized throughout the bone marrow niches in z, as were various ECM markers.

It was essential to study these markers throughout the z-planes of the system, as the endosteal and perivascular niches secreted different factors in different regions and were spatially separated in z. Furthermore, merely considering the extended-focus view of the devices gave a sometimes misleading view of the expression of ECM within the device. For example, as expected, type IV collagen was unambiguously expressed in and around the endothelial basement membrane, as shown by its extended-focus-apparent localization to the lectin signal in **Figure 3B(ii)**. Similarly, HA’s spindly morphology could be traced to pericytes wrapped around the lectin-labeled vessels (**Figure 3B(iv)**). However, markers like type I collagen and osteopontin showed up as diffusely-expressed and often-punctate signals throughout the extended focus images (**Figure 3B(i, v)**. Fibronectin, while having negative expression in the “grooves” where the LEL was expressed (**Figure 3B(iii)**), was still ambiguous about its perivascular expression. Therefore, we employed a similar 3D analysis of each z-plane as was used in **Figure 2E** to evaluate the relative expression of labeled patient BM-MNCs, labeled endothelium, and each ECM marker of interest (**Supplementary Figure 8B**).

From the summarization of each signal over z, given that each z-slice was acquired 4.4μm apart, type I collagen, fibronectin, and osteopontin were confirmed to be primarily endosteal in nature due to their signals peaking 4 z-slices (or 17.6 μm) apart from each other (**Figure 3B**, **Supplementary Figure 8B (i, iii, v)**). This was in direct contrast to collagen IV and HA’s peak differentials being 1-2 (4.4-8.8 μm apart) in **Figure 3B**, **Supplementary Figure 8B (ii, iv)**. As the z-resolution of the microscope we used was 11 μm due to the limits of its numerical aperture, the differences in peak between HA’s and collagen IV’s signals and lectin could not be resolved to different z-planes. In contrast, the differences in collagen I, fibronectin, and osteopontin could be, as 17.6 μm was within the capabilities of the microscope to resolve in z. However, that was not to say that these markers were not also found within the hydrogel compartment as well, which explained their curves’ shallow right shoulders as the signal dropped off from its peak. For example, osteopontin is canonically expressed by other cell types other than osteoblasts, such as endothelial cells, despite its name [43]. Similarly, fibronectin was a canonically endosteal-associated marker, supported by its peak shift of 4 z-slices as well (**Supplementary Figure 8B (iii)** and its strong expression in the EN-only devices of **Figure 1G(iii)**. However, it is known to also be secreted by pericytes as well, which could be seen slightly in the extended-focus view (**Supplementary Figure 8B(ii)**) [50, 51]. HA was similarly observed in both niches, particularly in a spindly morphology that suggested that it was strongly expressed by the MSCs [9].

The error bars in **Supplementary Figure 8B** are considerable, especially for the lectin signal of multiple groups. This was due to the relative nature of the confocal acquisition; the center of each device was located using the beacons discussed in previous sections. Then, the z-position with maximal lectin expression was selected, and then a 110 μm z-stack to capture the entire 3D volume was programmed around this focus point. This point was set qualitatively and the focus point of each device differed slightly. Additionally, gravity and differences in vascular network patterning resulted in the devices having different planes of focus for the hydrogel in space as well. As a result, the summarization of the signal could result in large variations due to the variation of the peak signal expression in z over different XY points in different devices. Therefore, qualitative evaluation of the images was always performed as well to serve as a “sanity check” for the observations from the quantitation.

After characterizing the cell and ECM components found within the hMM-on-a-chip using various techniques, we utilized the hMM-on-a-chip system to study the effect of therapeutic T cells. We initially performed our studies using T cells from healthy donors and fluorescent MM reporters, demonstrating these cells’ ability to travel to, proliferate against, and kill MM cells (**Figure 4**). However, some low-level release of T cell activation cytokines against the hBM-on-a-chip alone (**Supplementary Figure 11**) made it imperative to investigate the device’s ability to recapitulate graft-versus-host (GVH) effects *in vitro*. Graft-versus-host disease (GVHD) is less of a concern in MM and CAR-T cell therapies, as the stem cell transplantation tends to be autologous and the CAR-T cells are always autologous due to current lack of availability for allogeneic options. However, allogeneic stem cell transplantation is nevertheless prescribed on occasion [52]. Instead, our interest in GVH on-chip had more to do with ensuring that the observations we made were specific to the tumor cells rather than any off-target stroma, since that would limit the utility of our device technology.

However, the data demonstrated interesting differences between on- and off-target cells. First, CAR-T cells against the hBM-on-a-chip alone had mildly upregulated perforin and granzyme A expression. While there were universal slight increases, even at t=0hours, of TNF-α, IL-17a, and sFasL, they were also detectable in the media-only group, suggesting that the native hMM-on-a-chip was secreting these coordinating cytokines, which are not cytotoxic in and of themselves (**Figure 5**). Despite the granule proteases being detectable, we did not observe microscopic differences in vessel quality within CAR-T cell treated hBM-on-a-chip devices from previous data, so this upregulation was not highly concerning. Meanwhile, UTD-T cells against the hBM-on-a-chip alone similarly upregulated Granzyme A and perforin. Still, a similar lack of vascular network degradation caused us not to be concerned.

The more interesting observations came for the hMM-on-a-chip devices. First, as expected, the on-target, donor-matched UTD-T cells did not upregulate anything significantly from the media-only conditions, not even the perforin or granzyme A. We think this may have been due to the fact that the high number of primary BM-MNCs loaded within the PVN of the hMM-on-a-chip may have overcrowded the other stroma such that the T cells mainly “saw” their own matched-donor cells and so, therefore did not react; this was very promising. However, the off-target UTD-T secreted granule-associated perforin, granulysin, and granzymes, suggesting that they reacted strongly to the allogeneic BM-MNCs and the reaction was not BCMA-specific due to the lack of CAR expression. From the cytokine landscape within UTD-T cell conditions, we also saw that IFN-γ and TNF-α seemed to be antigen-recognition-specific cytokines. This made sense because the IFN-γ pathway was already identified as being particularly crucial for CAR-T cell cytotoxicity within solid tumors, suggesting that IFN-γ-IFNGR signaling was a key hallmark of CAR recognition [53] and because the 4-1BB transmembrane domain of the CAR molecule is known to upregulate the IFN-γ secretion pathway [54–56]. Furthermore, 4-1BB is a member of the TNF superfamily, and ECs were known to increase its expression after stimulation withm TNF-α. This suggested an autocrine loop for TNF-α secretion following 4-1BB stimulation downstream of CAR:BCMA crosslinking [54, 56]. Also, there was no significant difference in secretion level of IFN-γ between on- and off-target CAR-T cells (nor in sFasL), plus only on-target CAR-T cells secreted TNF-α. These suggested purely antigen-recognizing CAR-T cell mediated secretion. Therefore, the increased expression of TNF-α, IFN-γ, and sFasL were found to be potent predictors of antigen recognition within our system. The granule-secreted molecules (granulysin, perforin, and the granzymes) were less so, along with IL-17a. While there were significant differences between on- and off-target secretion for all five, the off-target versions of both cell types actually secreted more. That suggested that T cells secreted cytolytic granules against mainly non-BCMA+ cells, likely the other patient BM-MNCs within the system due to their abundance compared to the BCMA+ targets. Then, upon revisiting previous LEGENDplex data, we could strengthen our conclusions about healthy CAR-T cells inducing a BCMA-specific antitumor response within hMM-on-a-chip devices containing RFP-KMS18 (**Figure 4**).

Notably, the detected off-target cytokine secretion did not deleteriously affect the vascular network within the devices, as evidenced by the BioSpa tracking of healthy CAR-T and UTD-T cell-treated devices (**Figure 4C**). However, they may have been the reason why UTD-treated devices still had a plateau in RFP-KMS18 growth. Regardless, these data lent heavier weight to the measured levels of the CAR-specific cytokines TNF-α, IFN-γ, and sFasL. Furthermore, cytokine release data correlated well with image-based and flow cytometry-based metrics of target cell killing.

We next studied primary target cell killing within the hMM-on-a-chip system using matched-donor T cells from anonymous MM patients. Their prior lines of therapy were unknown, so we could not speak to the relative fitness of their starting cell populations [57]. Regardless, it was encouraging to see successful transduction of patient T cells despite previous MM treatment. Only a small percentage of the target cell population was a BCMA-expressing plasma cell (**Supplementary Figure 7**), but they were all labeled uniformly with the membrane dye and would not lose their fluorescence after death. As a result, primary cell target killing was more difficult to quantify using live-cell real-time microscopy. In order to mitigate this issue, we chose to apply a distance filter to the data, only considering the labeled target cell objects that ever came within 60 µm of a labeled T cell; the rationale for this algorithm was to only consider objects that could potentially have been influenced by the external effector cells added. Essentially, because T cells were allowed to flow randomly into the devices and could travel anywhere, depending on the vessel network architecture, it was possible that CAR-T cells were traveling to regions with very little target cell presence. Especially given the already-low plasma cell percentage within these devices, this could unfairly mask differences in signal. Therefore, we calculated the Euclidean distance of each T cell to each labeled BM-MNC within the devices. Then, we could filter out the target cell objects that never traveled within 60 μm of a T cell.

The 60 µm threshold was chosen qualitatively based on the hypothesized speed of T cell movement within the system. Lenient filtration would not solve the issue of signal masking. Still, conservative filtration could choke out T cell objects that potentially traveled near a target cell between sampling time points and then traveled slightly too far away to be considered. The 60 μm choice resulted in a 10-30% reduction in measured T cell objects, which we subjectively deemed to be appropriately restrictive (**Supplementary Figure 14**). However, this filter choice was the reason we chose to report any single-cell histogram data as normalized so that the area under the curve (AUC) was equal to 1, to prevent confusion from histograms with different numbers of reported objects. The distance filter did not change the reported results for the healthy-donor T cell experiment (data not shown), probably due to the lack of ambiguity in target cells. By definition, any red-fluorescent object in that experiment was RFP-KMS18 and therefore had BCMA expression and was sensitive to CAR-T cell-mediated cytotoxicity. In the patient-specific experiment, however, the filter helped to reduce noise from T cells that already were too far away from any potential targets.

We also attempted to use real-time caspase tracking to quantify the apoptosis of target cells, as a positive metric for cell killing. There were no significant differences in caspase signal between CAR and UTD, even with distance filtration. This was likely due to the nonspecific universal labeling of all apoptotic, labeled target cells and stromal cells within the hMM-on-a-chip. Since the caspase dye universally labeled all cells, which may have had any number of reasons for undergoing programmed cell death, it was not possible to discern CAR-mediated apoptosis within the system significantly. However, we qualitatively observed increases in caspase activity within the microscopy images (**Supplementary Figure 15**).

While there were definitely large numbers of more terminally-differentiated effector memory T cells (T_EM_s) and T_eff_s found within the devices after extended culture, these data were promising suggestions that more immature T cell memory phenotypes could be more durable within the hMM-on-a-chip’s TME (**Figure 6**). This supported the commonly accepted dogma about desirable T cell memory subsets for CAR-T cell therapy. Naïve T cells (T_N_s), T_SCMs_, and T_CMs_ are widely associated with longer-lasting antitumor effects in adoptive cell transfer treatments. This may be due to more immature phenotypes being able to express genes associated with lymphoid homing and release cytokines that contribute to the milieu that recruits a more complete immune response, as well as the risk of more terminally differentiated cells entering into pro-apoptotic and/or senescent states [58, 59]. In fact, it has been shown that more terminally differentiated effector and effector memory T cells can interact with “younger” T cells to promote their differentiation and impair their efficacy [60]. While T_eff_s can efficiently kill target cells, they are terminally differentiated. Therefore, there is interest in creating CAR-T cells that belong to a less differentiated subset. These cells are longer-lived (which was recapitulated in our system) and can readily differentiate into more toxic effector cells when they encounter the target antigen, enabling potentially longer progression-free survival. Importantly, CD8+ T_SCM_ cell numbers and CD4:CD8 ratios were predictive of BCMA CAR-T cell efficacy in recent MM clinical trials [5, 61].

We found that both healthy and patient T cells could cause detectable target cell cytotoxicity within the hMM-on-a-chip, using a variety of analysis methods to complement one another. Furthermore, we could see the preservation of several key memory T cell subpopulations after prolonged culture within the device, which is promising for using this technology to study effector cell durability *in vivo* to potentially develop a cure for multiple myeloma.

## MATERIALS AND METHODS

### Experimental Design

We fabricated human multiple myeloma on-a-chip (hMM-on-a-chip) devices by assembling the consumable components and then culturing the cell types of interest within. These cell types were often labeled beforehand for visualization in real-time. We then collected and analyzed data using immunofluorescence microscopy, flow cytometry, and cytokine analysis.

### Device Fabrication

Details of device fabrication reagents are in Supplementary Materials (**Table 1**).

**Table 1:**
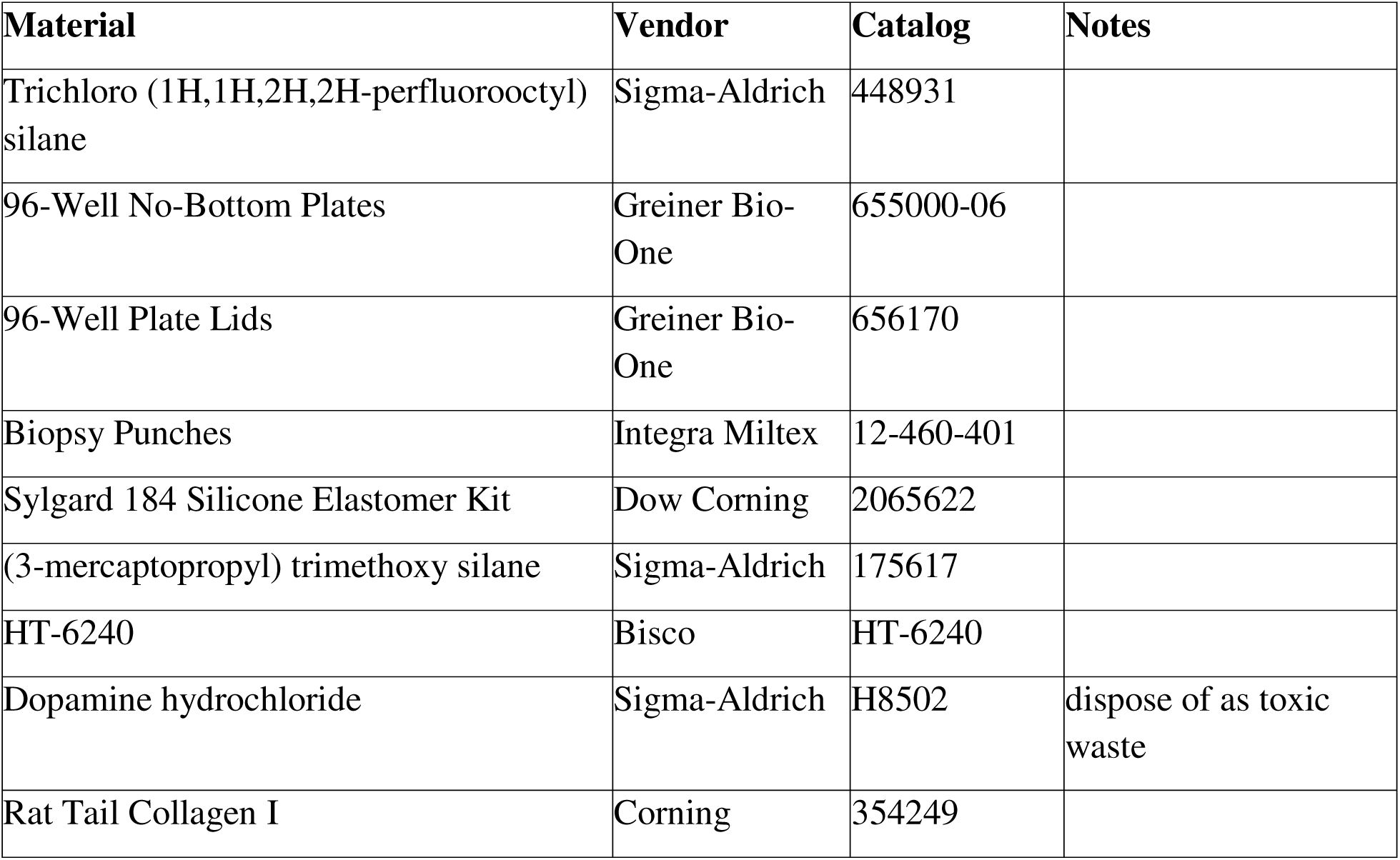
Device fabrication materials list.

#### PDMS casting

Wherever PDMS (Dow Corning) was used, it was mixed thoroughly 10:1 (elastomer base: curing agent) by weight, cast onto the mold, and allowed to cure at 65LJ for at least 4 hours.

#### Channel fabrication

Channel dimensions were as previously described [31]. The silicon wafer used for device making was commissioned from MuWells, Inc. PDMS was used to create a master cast of the patterned wafer. Then, EpoxAcast 670 HT (Smooth-On) was used to create a replica of the channels that could be used to cast subsequent devices. Multiple molds were created from the same master and reused to increase device production throughput.

To create the channels for each plate of devices, PDMS was cast on the molds to a thickness of approximately 2mm. Loading ports were bored into cured PDMS using a 1 mm biopsy punch (Integra Miltex).

### Plate bonding

After hole-punching, the feature layer of PDMS was bonded to a bottomless 96-well plate (Greiner Bio-One) using a chemical gluing method [62]. Briefly, the 96-well plate was immersed in 2% (v/v) 3-mercaptopropyl trimethoxysilane (Sigma-Aldrich) in methanol for 2 minutes to functionalize the polystyrene with silane functional groups. The excess was rinsed with deionized water (DI H_2_O), and the plates were thoroughly dried. Within 30 minutes, these treated plates were plasma bonded to the PDMS features using a plasma cleaner, followed by a 65LJ incubation overnight to finish the bonding process.

#### Plate washing and port correction

After the channels were bonded to the plate, they were washed by immersion in 70% EtOH. As this ethanol drained, it emptied from wells that contain punched ports. Any wells with ports that retained liquid likely had improperly-punched ports. A 1-mm wire or biopsy punch was used at this stage to correct any ports and facilitate proper drainage. After immersion and correction, excess ethanol was discarded, and plates were allowed to fully dry at 65LJ for at least 30 minutes prior to film bonding.

#### Film bonding

In order to seal the PDMS feature layer, the bottom surface was covalently bonded to commercially-purchased silicone film, HT-6240 (Bisco). The plastic backing was removed from one side, and the exposed silicone was bonded to the channel surface of the devices using the same plasma cleaning protocol that was used in prior steps.

#### Ethanol sterilization

Devices were sterilized prior to culturing cells within. All 5 channels of each device (2 outer, 2 media, and 1 central) were flooded thoroughly with 70% ethanol. This ethanol was allowed to fully evaporate at 65LJ overnight, after which the devices were only opened in a biosafety cabinet.

#### Dopamine-Collagen I treatment and UV sterilization

After the devices were sterilized, dopamine was used to render the central channel PDMS hydrophilic to facilitate collagen I adhesion and downstream cell adhesion, as described previously [63]. Briefly, sterile 0.1 mg/mL dopamine HCl (Sigma) was pipetted into the 1mm port of the central channel of each device. After 1 hour of room-temperature incubation, dopamine was washed out with an excess of sterile 1X PBS. Then, 20 μg/mL sterile collagen I solution in PBS was loaded similarly and incubated for 1 hour at room temperature, then washed out with excess 1X PBS as previously. Devices were allowed to dry thoroughly at 65LJ prior to central channel loading and used within 1 week of treatment. Before cell culture, devices underwent secondary 365nm UV sterilization for at least 90 minutes.

### Cell culture

All cell culture reagent details are in Supplementary Materials in **Table 2**.

**Table 2:**
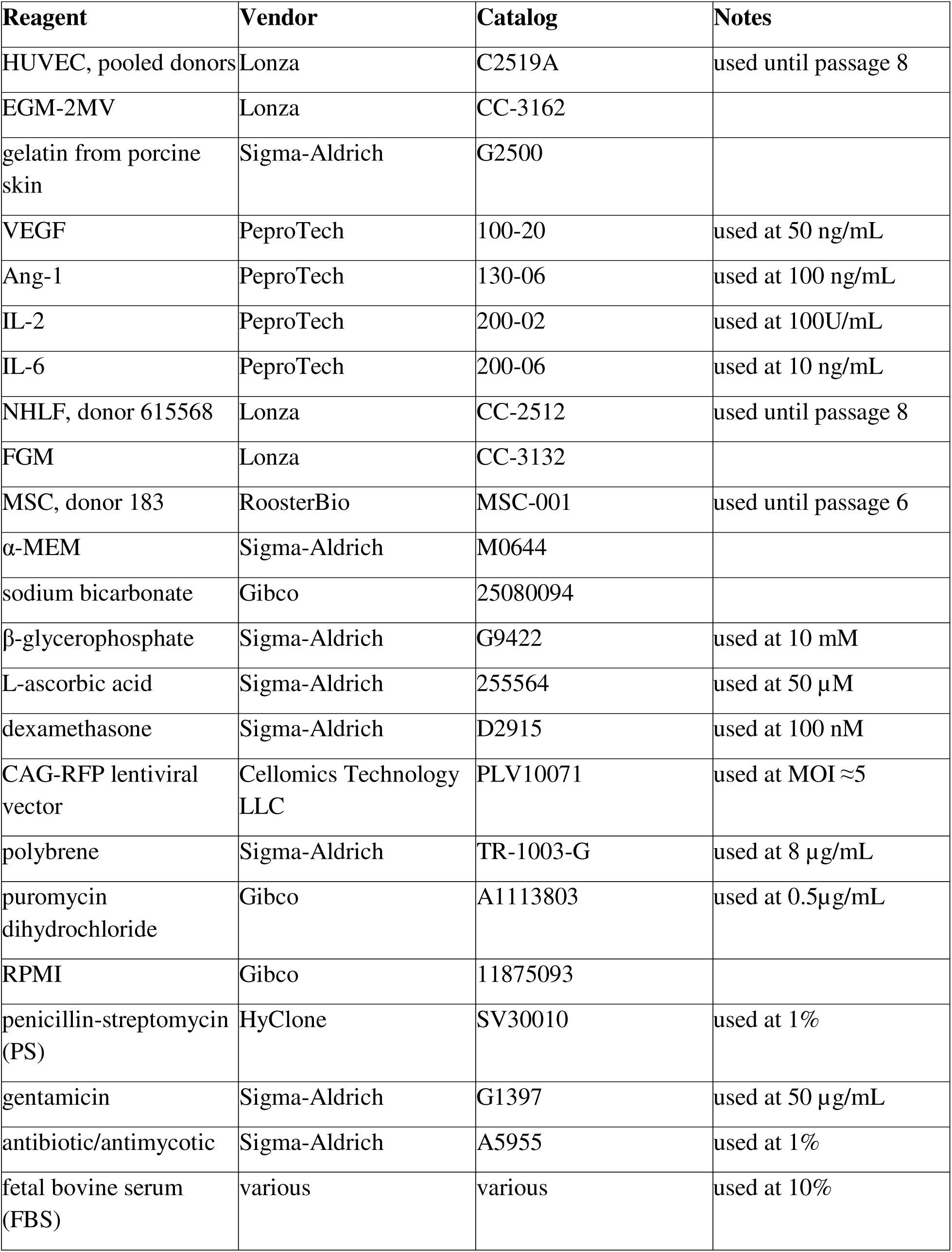
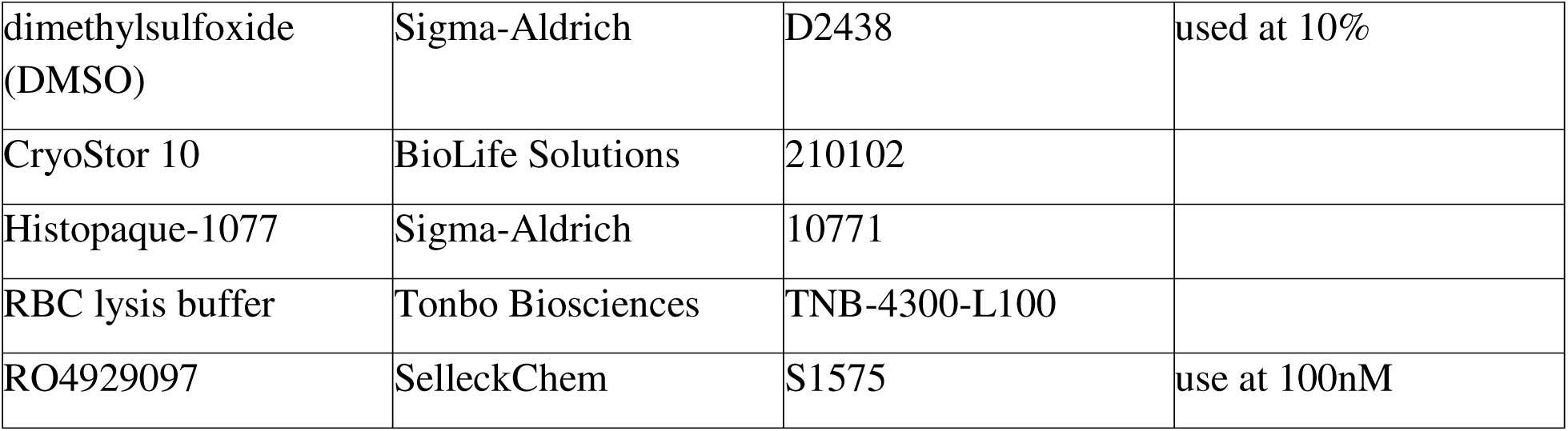
Cell culture reagent list.

#### Mesenchymal Stromal Cells (MSCs)

Human BM-derived MSCs (RoosterBio, donor 183) were verified by flow cytometry to be CD34-, CD45-, CD73+, CD90+, and CD105+. MSCs were expanded and cultured on tissue culture polystyrene (TCPS) in αMEM (Sigma Aldrich) supplemented with 10% fetal bovine serum (FBS) (Hyclone) and 1% penicillin-streptomycin (PS) (Hyclone). MSCs were cultured at 37LJ and 5% CO_2_. MSCs were used up to passage 6 and harvested by incubating with TrypLE (Gibco) to gently detach the cells from the culture surface.

#### Human Umbilical Vein Endothelial Cells (HUVECs)

HUVECs from pooled donors (Lonza) were cultured in microvascular endothelial growth medium (EGM-2MV) (Lonza) on 0.1% bovine gelatin (Sigma-Aldrich) coated TCPS at 37LJ and 5% CO_2_. HUVECswere used up to passage 8 and harvested akin to MSCs.

#### Normal Human Lung Fibroblasts (NHLFs)

NHLFs (Lonza) were cultured in fibroblast growth medium-2 (FGM) (Lonza) on TCPS at 37LJ and 5% CO_2_. They were used up to passage 8 and harvested akin to MSCs.

#### Primary bone marrow mononuclear cells (BM-MNCs)

Primary bone marrow mononuclear cells (BM-MNCs) from MM patients were obtained pre-frozen from Emory’s Winship Cancer Center. They were pre-characterized with their plasma cell percentage (defined as a percentage of cells within the population that were CD138+ CD38+ CD45-), and cryovials were identified only by this percentage and their clinical trial ID number. They were thawed and rested overnight in complete RPMI before labeling (if applicable) and culture. These primary cells were never subcultured.

#### MM cell lines

KMS18, OCIMY5, OPM2, RPMI8226, and U266 were cultured in complete RPMI and subcultured by half roughly every 2-3 days, or when the cell concentration reached approximately 5e5 viable cells/mL. The culture vessels were TCPS in 37LJ and 5% CO_2_. As they were all suspension cell lines, they could, therefore, be passaged without trypsinization. However, as RPMI8226 and U266 were semi-adherent, those flasks were scraped before cell harvest.

### Creation of red fluorescent protein (RFP)-expressing cell lines

In order to monitor MM cell proliferation through a reporter gene without relying on membrane labeling dyes, lentiviral transduction was used to create RFP-expressing MM cell lines. Briefly, a lentiviral vector encoding for CAG-promoter TurboRFP (Cellomics Tech) was used to transduce wild-type MM cells according to manufacturer’s protocol and purified using puromycin resistance.

### Device Culture

#### Osteogenesis in hBM-on-a-chip and hMM-on-a-chip

For the formation of the endosteal niche as previously described [31], MSCs were seeded within the central gel channels of devices at a density of 3·10^4^ cells/device. Cells were cultured within the devices for 21 days in αMEM osteogenic media (10% FBS, 1% PS, 50 μg/mL gentamicin (Sigma-Aldrich), 10 mM β-glycerophosphate (Sigma-Aldrich), 50 μM ascorbic acid (Sigma-Aldrich), and 100 nM dexamethasone (Sigma-Aldrich)).

#### Vasculogenesis in hBM-on-a-chip and hMM-on-a-chip

Vasculogenesis in the central gel channel was accomplished using previously reported approaches [31, 64, 65]. Briefly, after washing the central channel with PBS, HUVECs (4.5·10^6^ cells/mL) and MSCs (6·10^5^ cells/mL) were suspended in EGM-2MV supplemented with thrombin (4 U/mL) (Sigma-Aldrich). A solution of fibrinogen (8 mg/mL) (Sigma-Aldrich) and collagen I (2 mg/mL) (Corning) in PBS was mixed thoroughly with the thrombin-supplemented human umbilical vein endothelial cell (HUVEC)/MSC cell suspension, and 5 μL was immediately loaded into the central gel port. Devices were then incubated for 15 minutes at 37LJ, 5% CO_2_ to allow the fibrin gel to form. Vasculogenic media, EGM-2MV supplemented with 50 ng/mL VEGF, was added to both media channels and pulled through with 10 μL of negative pressure. Cells were cultured with daily media exchange for 5 days to allow for vasculogenesis. Media was supplemented with VEGF (50 ng/mL) on all days and Angiopoietin 1 (Ang-1) (100 ng/mL) (Peprotech) from day 3 onward.

#### Culture of MM cells within hMM-on-a-chip

Additional cell types of interest, such as primary or cell line-based MM, were optionally labeled with PKH26 (red), PKH67 (green), CellVue Claret (far red), or CellTrace Violet (blue) dye according to manufacturer instructions (Sigma-Aldrich), or left unlabeled. They were then loaded in the thrombin cell suspension prior to and loaded during the previously-described vascular hydrogel loading step.

#### Live-cell imaging in hMM-on-a-chip

In order to monitor the behavior of cells cultured within the hMM-on-a-chip, nondestructive microscopy was the default choice. For most cases, this could be achieved using standard epifluorescence or confocal microscopy, as both instrument types were equipped with temperature and gas control capabilities. Thus, the green (i.e., GFP or PKH67), red (i.e. RFP or PKH26), far-red (ie DyLight 649 or CellVue Claret), and blue (ie CellTracker Blue or CellTrace Violet) fluorescent channels of each device could be imaged. For long-term automated imaging, the BioTek Cytation 3 epifluorescence microscope was used, as it was paired with a BioTek BioSpa automated incubator. This incubator could culture up to 8 96-well plate devices at 37LJ and 5% CO_2_ and be programmed to periodically image them over extended culture periods using a robotic arm.

ImageJ was used to compile single-channel, single-timepoint images into a single multidimensional (X, Y, time, and channel) image for each device. Then, ImageJ was used to gather relevant metadata and single-cell measurements for each image, channel, and timepoint. Finally, Python was used to compile, filter, and unblind each measurement to categorize the data into the different experimental groups for downstream plotting and statistical analyses.

#### Vessel tagging in live devices

In order to label vessel networks within devices during ongoing culture, the culture medium was supplemented with 0.25 µg/mL fluorescent *Lycopersicon esculentum* lectin (LEL) (Vector Labs) and allowed to incubate for at least 4 hours before acquisition of the first images, which could be acquired using either epifluorescence or confocal microscopy.

#### Effector cell coculture within hMM-on-a-chip

During experiments requiring the introduction of effector T cells, T cells were either used fresh following a 10-day expansion or rested overnight in 100U/mL IL-2. Before culture, depending on the experiment, they were dyed with 10 μM CellTrace Violet (CTV) according to the manufacturer’s instructions or left unlabeled. Then, the cells could be flowed into the media channels of the devices about 48 hours after initiation of vasculogenesis. Cell behavior was studied longitudinally using imaging and media sampling or flow cytometric analysis at the endpoint.

### Preparation and analysis of devices using flow cytometry

Flow cytometry could be performed on digested devices in order to yield high-dimension surface marker expression data at the cost of spatial resolution of such signals. Generally, to yield enough cells in each sample to obtain meaningful results, digested devices were pooled together such that cells from 2-3 devices were present in each test sample. All flow cytometry reagent details may be found in Supplementary Materials, in **Table 4**.

#### Cell Harvest

0.25% trypsin (Gibco) was added to each device and incubated for about 30 minutes at 37LJ. Gels were then harvested and quenched with complete RPMI. Devices were imaged pre- and post-harvest to confirm the balled-up morphology of trypsinized cells and then to confirm that all of the cells were harvested from the now-empty devices. Then, the cells were pelleted, washed, stained, and analyzed. This trypsin formulation degraded the fibrin-collagen hydrogel of the central channel but left the pure fibrin hydrogel of the outer channels mostly intact, preventing the need to account for fibroblast presence in the cell samples.

#### Staining and fixation

Cell samples were stained for viability using amine-reactive dyes in various colors (BioLegend). Then, they were either fixed with 4% formaldehyde-containing buffer (BD) for thirty minutes or immediately stained for surface markers and then fixed. Depending on whether the sample type contained FC receptors, an FC-blocking step was performed before antibody incubation. Samples were run on the Cytek Aurora spectral flow cytometer and then analyzed in FlowJo.

#### 2D CAR-T cell killing assay

Chimeric antigen receptor (CAR) and UTD-T cells were thawed and rested overnight in 100U/mL IL-2. CAR-T cells were previously transduced using a chimeric antigen receptor construct encoded to recognize the B cell maturation antigen with a 4-1BB costimulatory domain [66]. These cells were then dyed with CTV according to the manufacturer’s instructions. Then, target cells, RFP-expressing KMS18 (RFP-KMS18), were plated in EGM-2MV supplemented with VEGF and Ang-1 to simulate the media conditions found within the hMM-on-a-chip. The effector cells were added at a range of effector cell to target cell (E:T) ratios and the plates were cultured within the BioSpa imaging platform to image the wells periodically. The E:T ratios tested were slightly different due to each cell type’s availability; the UTD-T cells’ yield forced the maximum tested E:T to be lower. At the end of the culture period, media was sampled from the wells for multiplexed cytokine analysis, and the plate was stained for viability and run on the CytoFLEX S flow cytometer to measure the number of red-fluorescent viable target cells at the end. This value was normalized to the number of viable cells in the target cell-only wells to yield the % cytolysis. The nonlinear fit of each E:T curve for both CAR and UTD conditions and their half-maximal inhibitory concentration (IC50) were calculated using GraphPad Prism.

#### Multiplexed cytokine detection

Media was collected from devices at designated time points by tilting the plates to enable the pooling of media in downstream media wells. Then, 50 μL could be collected or all media was collected at the endpoint. Device media was stored at −20LJ prior to analysis. Samples were randomized on the assay plates and analyzed using the LEGENDplex human CD8/NK panel (BioLegend). Samples were read on a CytoFLEX S flow cytometer (Beckman Coulter). Sample unblinding and downstream analyses were performed in Python and GraphPad Prism.

### Preparation and analysis of devices using immunohistochemistry (IHC)

All IHC reagent details are in Supplementary Materials (**Table 3**).

**Table 3:**
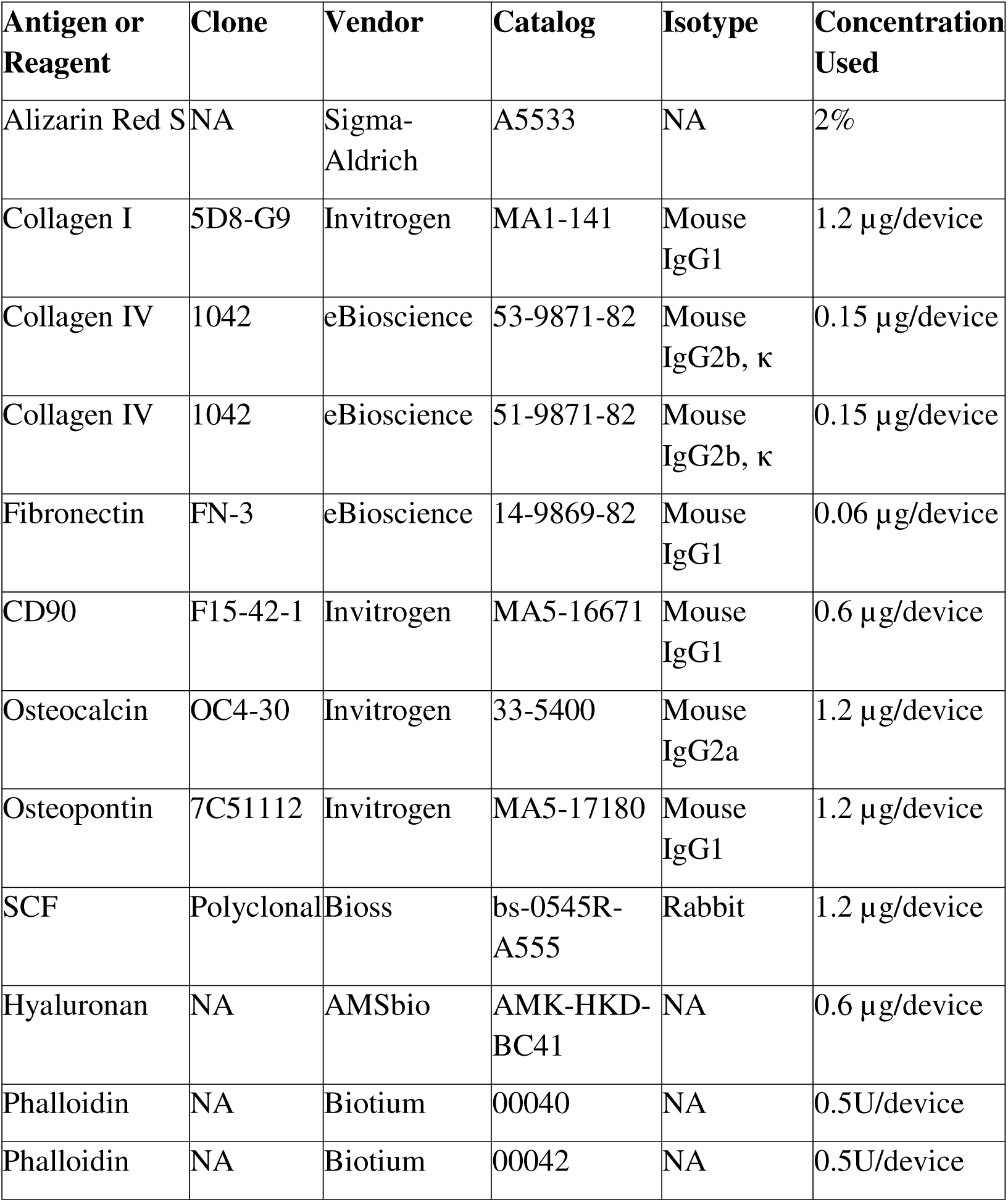
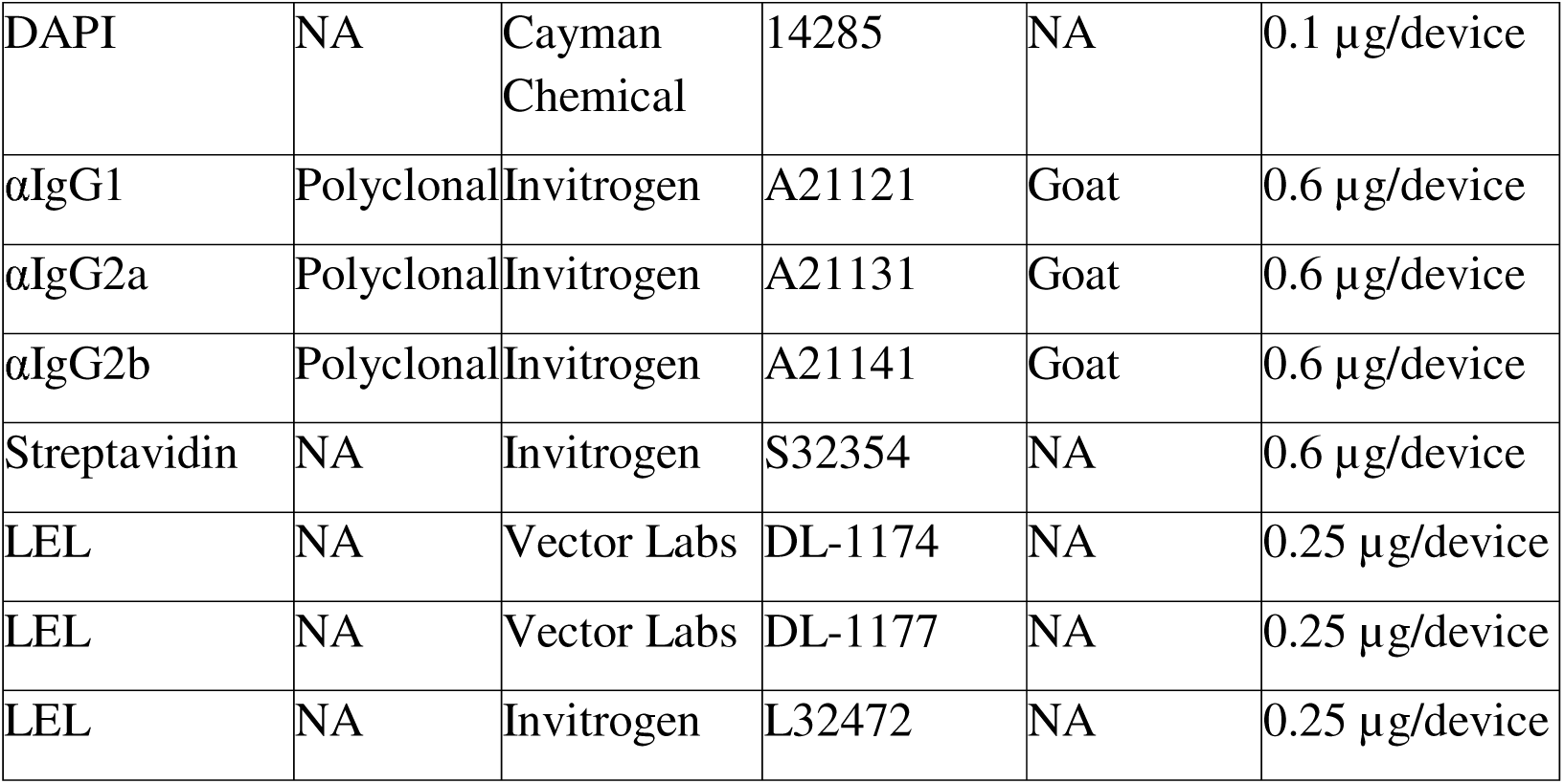
IHC antibody list.

**Table 4:**
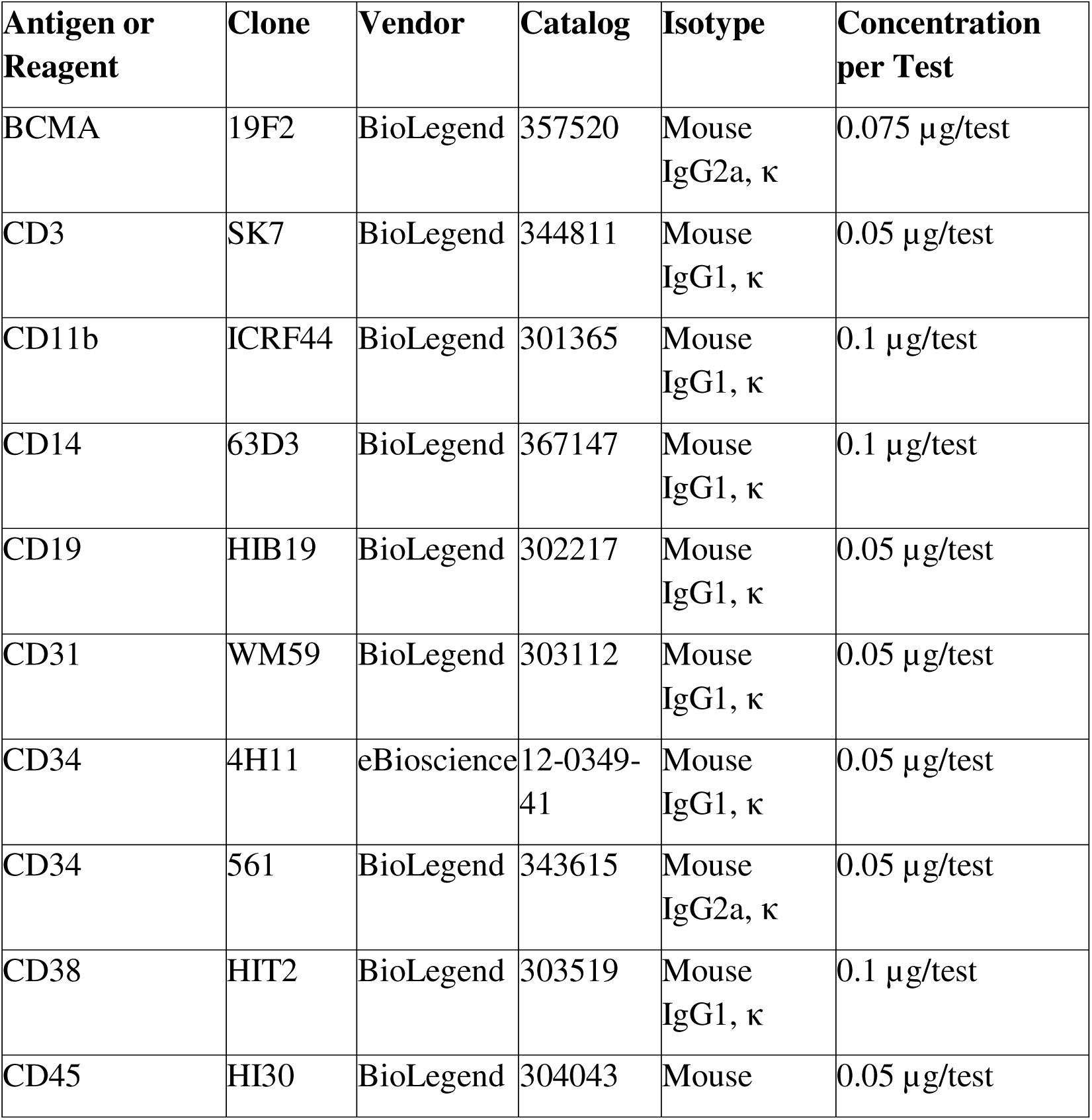
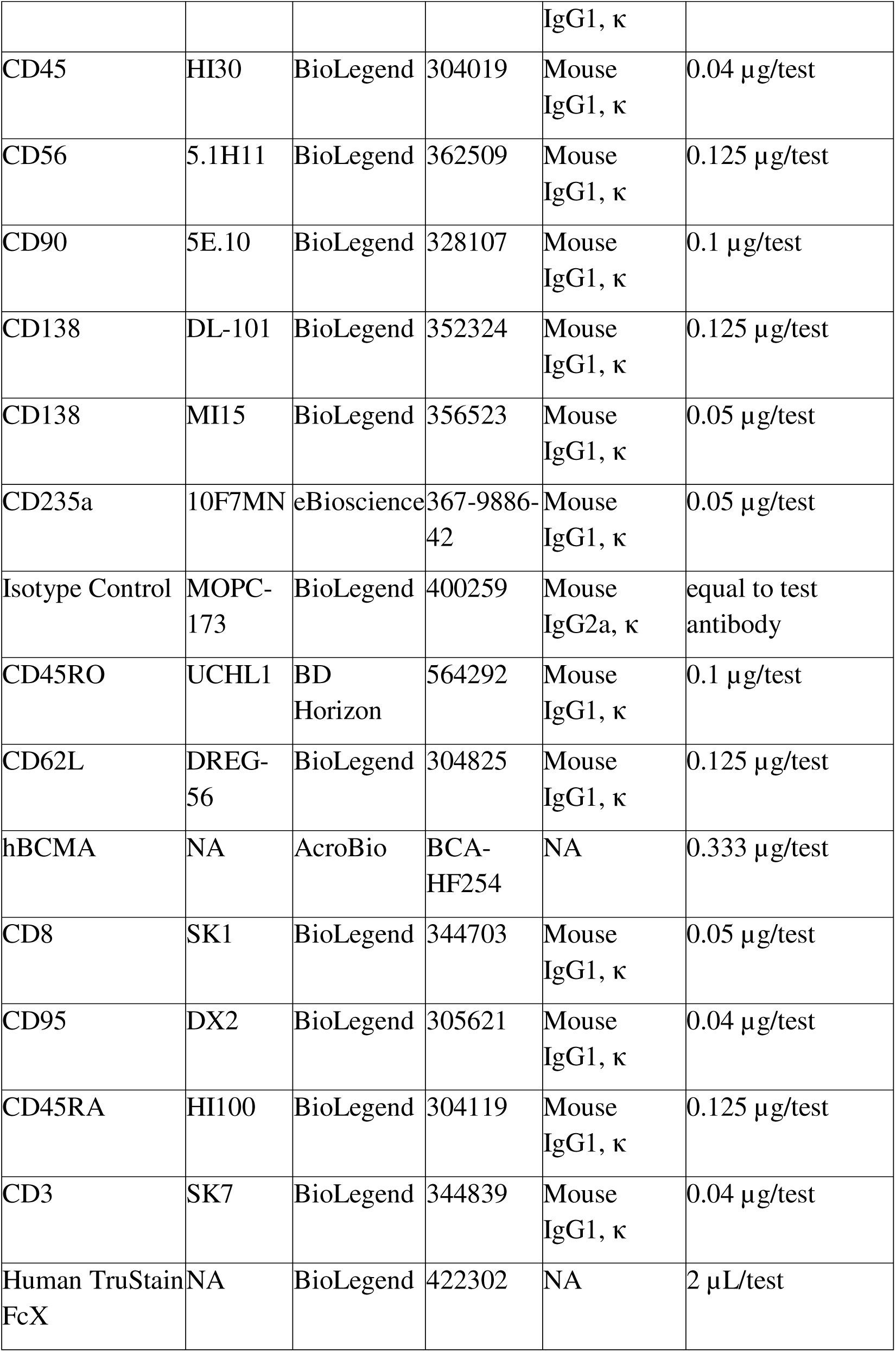
Flow cytometry antibody list.

#### Fixation and Permeabilization

Prior to staining and downstream analysis, devices were fixed with formaldehyde to halt cellular processes. Each device was washed with PBS, then 4% formaldehyde in PBS was added for 15 minutes. Devices were washed twice more with PBS and could then be stored long-term, protected from light, at 4LJ.

After fixation, if it was desired to analyze intracellular markers like actin, devices were permeabilized with 0.1% Triton-X 100 for 15 minutes. Devices were washed twice more with PBS and could then be stored long-term, protected from light, at 4LJ.

#### Alizarin red quantification

For alizarin red staining, devices were fixed and then washed twice with DI H_2_O. They could then be stained for 5 minutes with 2% alizarin red (Sigma-Aldrich) in DI H_2_O [pH 4.1-4.3]. Alizarin red stain was removed by several washes with DI H_2_O, until liquid was clear. Stained devices were imaged using a Lionheart FX (BioTek Instruments). Color brightfield images were analyzed using ImageJ’s open-source software [67, 68].

#### IHC staining and microscopy

Prior to staining, if using unconjugated primary antibodies, devices were blocked with 5% bovine serum albumin (BSA), and 3% goat serum in PBS for ≥ 2 hours at room temperature. Primary antibodies were diluted (typically to 1-10 μg/device) in a blocking buffer, and devices were stained overnight at 4LJ. Devices were then washed with 0.1% BSA in PBS and, if necessary, stained with secondary antibodies or streptavidin (typically 10 μg/mL) diluted in wash buffer for 2 hours at room temperature. Devices were washed with wash buffer and then imaged using either a Lionheart FX (BioTek Instruments) epifluorescence microscope or a spinning disk confocal microscope (PerkinElmer) at 4X, 10X, or 20X magnification; the bottom film was too thick for higher magnification lenses, which tend to have shorter working distances and oil-objective lenses. For cyclic immunofluorescence (CycIF) studies, stained devices were incubated with bleaching solution (4.5% hydrogen peroxide, 25 mM NaOH in PBS) solution for 1 hour at room temperature, exposed to white benchtop light, and then washed with PBS before re-imaging with new conjugated antibodies.

### Statistical Analysis

All statistical analyses were performed in GraphPad Prism, and the relevant statistical tests, sample sizes, and P-values can be found in each figure caption.

## CONCLUSION

In summary, this work presented a novel model of human multiple myeloma that could be used to recapitulate effector cell cytotoxicity against tumor cells within a spatially organized tumor microenvironment.

The human multiple myeloma on-a-chip (hMM-on-a-chip) design was compatible with laboratory- and industry-standard equipment and techniques. This lent itself beautifully to seamless automation of both data acquisition and analysis. To further support its robustness, the hMM-on-a-chip was fabricated from only 3 components, 2 of which were commercially available for better reproducibility.

Next, the perfusable microvascularized network within the device could be used to model highly-vascularized human bone marrow, particularly the perivascular niche. This would enable modeling more physiologically-relevant phenomena, such as T cell trafficking through vessels to encounter and activate against target cells.

Finally, there is a significant gap in knowledge about the phenotypic changes undergone by adoptively transferred T cells, which is difficult to study in human subjects. Our model was based on human cells and enabled the systematic inclusion or removal of any cell type of interest. Therefore, we were able to harvest and study T cells pre- and post-“implantation” within the system in order to better understand the immunomodulation that occurs in the immunosuppressive microenvironment of MM.

## ACKNOWLEDGMENTS

OPM2 was a gift from Madhav Dhodapkar. KMS18, U266, RPMI8226, and OCIMY5 were gifts from Larry Boise. We would like to acknowledge the Georgia Tech Parker H. Petit Institute Core Facilities for their services and shared resources that enabled us to produce this publication. Parts of this work were performed at the Georgia Tech Institute for Electronics and Nanotechnology, a member of the National Nanotechnology Coordinated Infrastructure (NNCI), which is supported by the National Science Foundation (ECCS-2025462).

## Funding

Georgia Tech Foundation (KR)

Georgia Tech Research Alliance (KR)

Marcus foundation (KR)

NSF Engineering Research Center for Cell Manufacturing Technologies (CMaT) (NSF Grant EEC 1648035 (KR)

NSF Graduate Research Fellowship under Grant No. DGE-1148903 (DG)

Career Award at the Scientific Interface from Burroughs Wellcome Fund (AFC)

Bernie Marcus Early-Career Professorship (AFC)

National Institutes of Health Grant (R35GM151028) (AFC)

Paula and Rodger Riney Foundation (SL)

NIH R35HL145000 (WAL)

## Author contributions

Conceptualization: DG, KR

Methodology: DG, IP, RR, AFC

Investigation: DG, IP, RR, LK, EB, TH, AR, RP, JC, SA, AT, SR, JT, ED, FK, AO, NS

Visualization: DG

Key experimental reagents: SL, JNK, RT, and MZ

Supervision: AFC, KR

Writing – original draft: DG

Writing – review & editing: IP, KR

## Data and materials availability

All data, code, and materials used in the analyses and depicted in the main text or supplementary materials are available upon request by contacting the corresponding author, Dr. Krishnendu Roy. The BCMA CAR constructs were provided under a materials transfer agreement (MTA) between the Georgia Institute of Technology and the National Institutes of Health.

## SUPPLEMENTARY

**Supplementary Figure 1:**
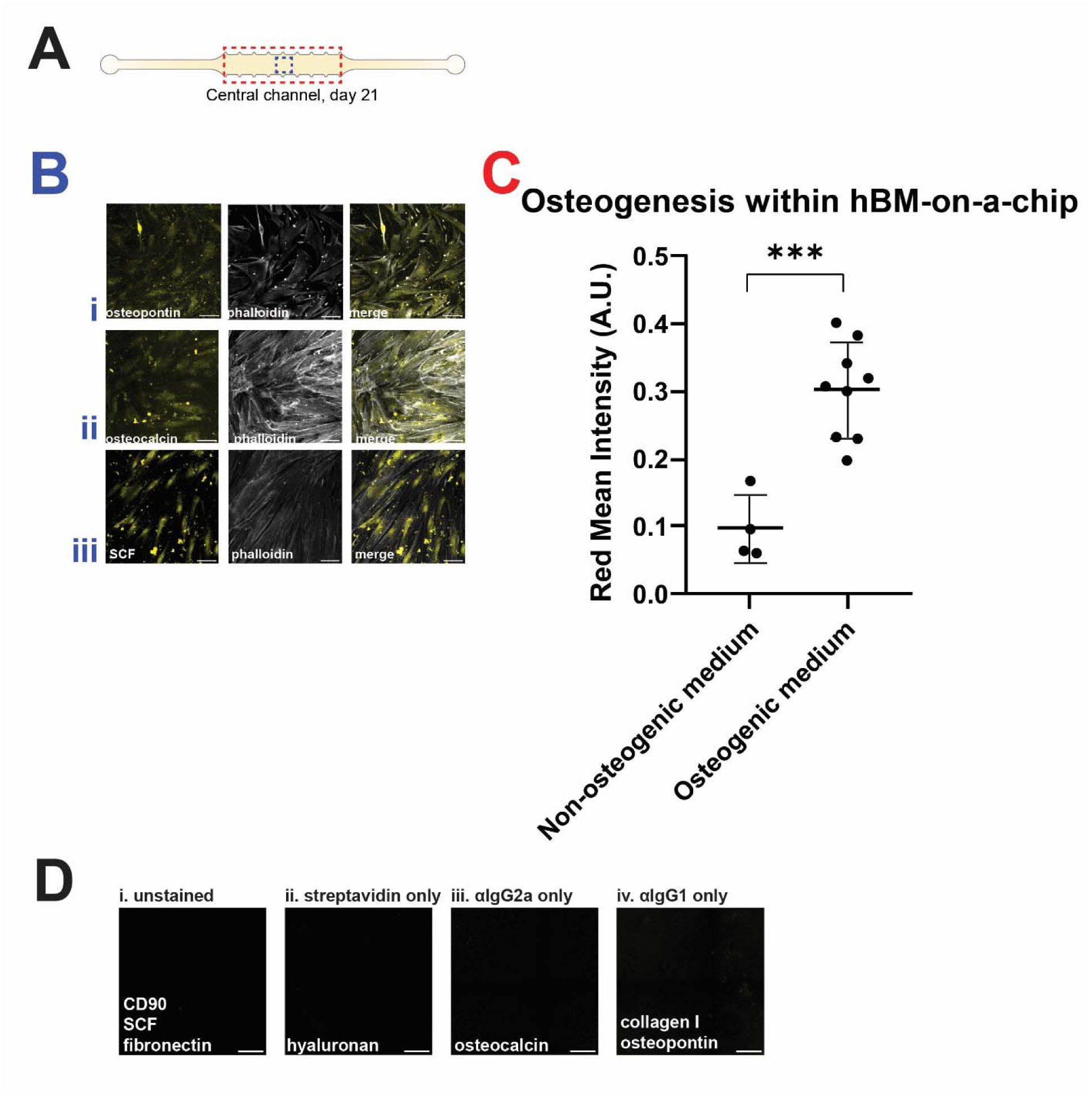
Overview of hBM-on-a-chip and hMM-on-a-chip. (A) cartoon depiction of the central channel, where analyses were conducted. The blue dashed box signifies approximately where (B) extended-focus confocal images of (i) osteopontin, (ii) osteocalcin, and (iii) stem cell factor, counterstained for actin, were collected (representative images). The red dashed box signifies where (C) alizarin red was used to label calcium deposits left in devices by the MSC-derived osteoblasts (n=9) or undifferentiated MSCs (n=4). These deposits were quantified for their red intensity in AU. *** = p ≤ 0.001, Welch’s non-parametric t-test. (D) IHC control images for (i) directly-conjugated antibodies (CD90, SCF, and fibronectin), (ii) biotinylated HA binding protein, and unconjugated antibodies against (iii) osteocalcin and (iv) collagen I and osteopontin.

**Supplementary Figure 2:**
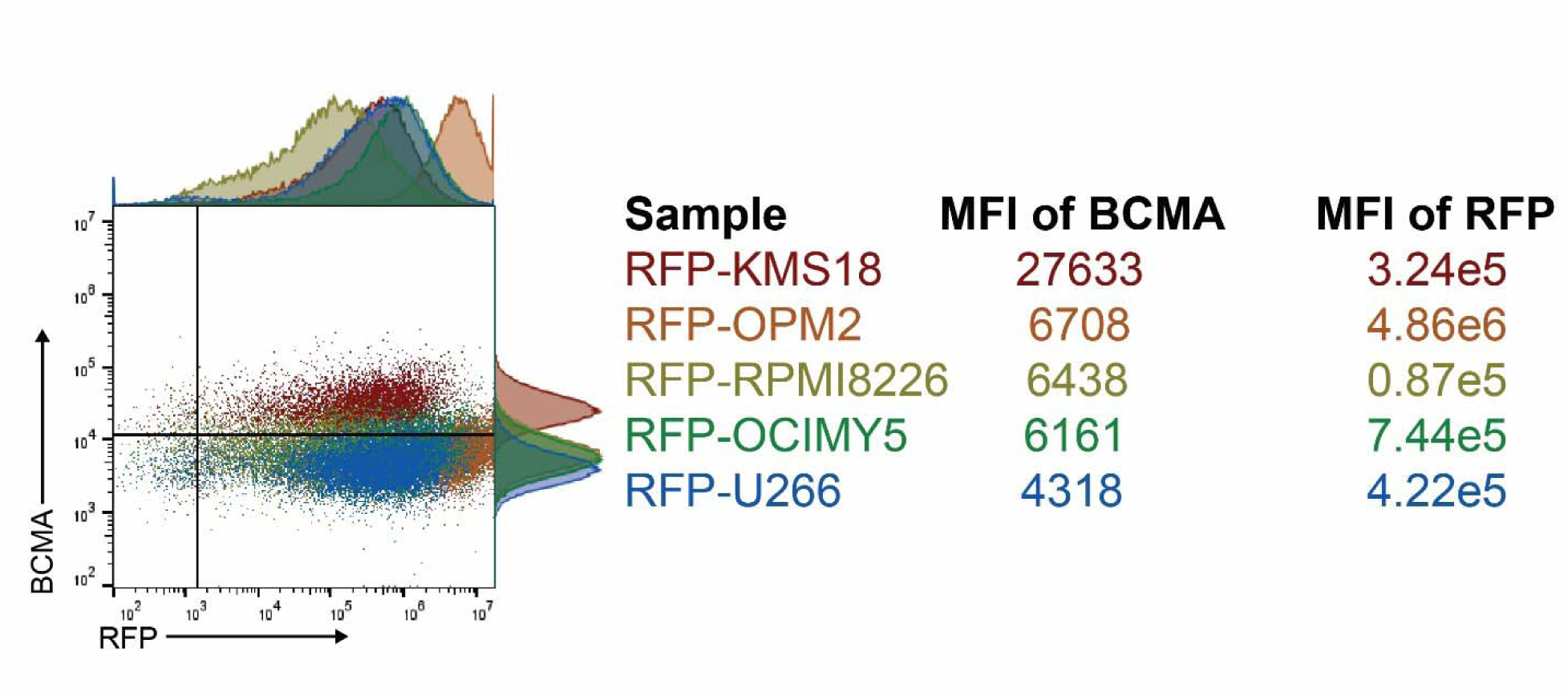
Characterization of reporter MM cell lines. (A) Flow cytometric analysis of RFP expression plotted versus BCMA expression showed varying, albeit detectable, degrees of expression.

**Supplementary Figure 3:**
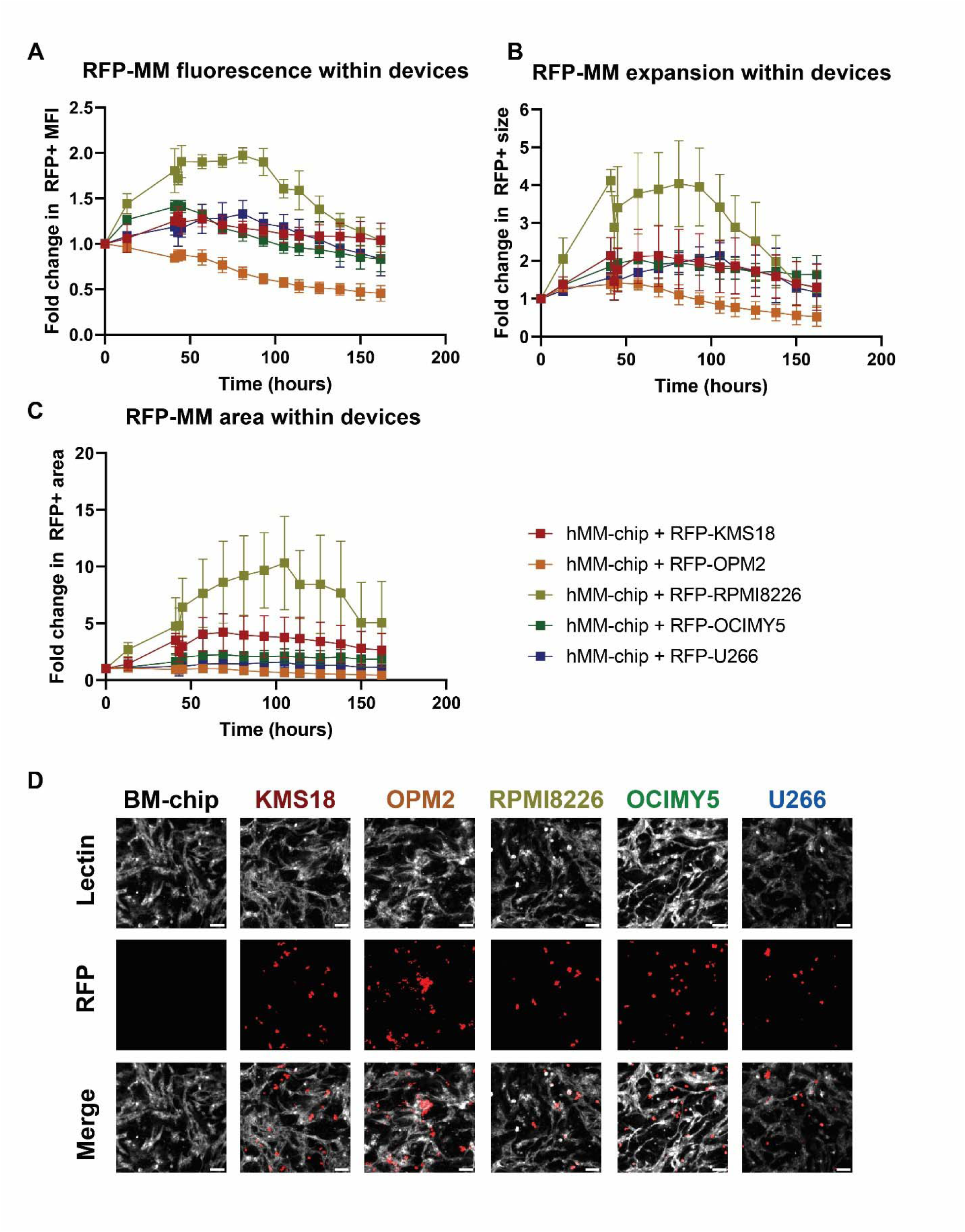
Further quantification of MM reporter lines within hMM-on-chip. All figure axes were chosen to match those in the main figure. (A) The relative MFI of MM objects for 4 of the cell lines did not decrease; OPM2’s loss of MFI could indicate the cells were not surviving well within the system, as supported by the proliferation data. (B) Average object size and (C) Number of objects for the 5 considered cell lines. Because object size increased in several cases while number of objects often remained stable, this suggested potential clonal expansion of the cell lines. This is why area was chosen to quantify MM cell growth within the system. (D) Clonal expansion and perivascular localization for each cell line. Scalebars = 100 µm.

**Supplementary Figure 4:**
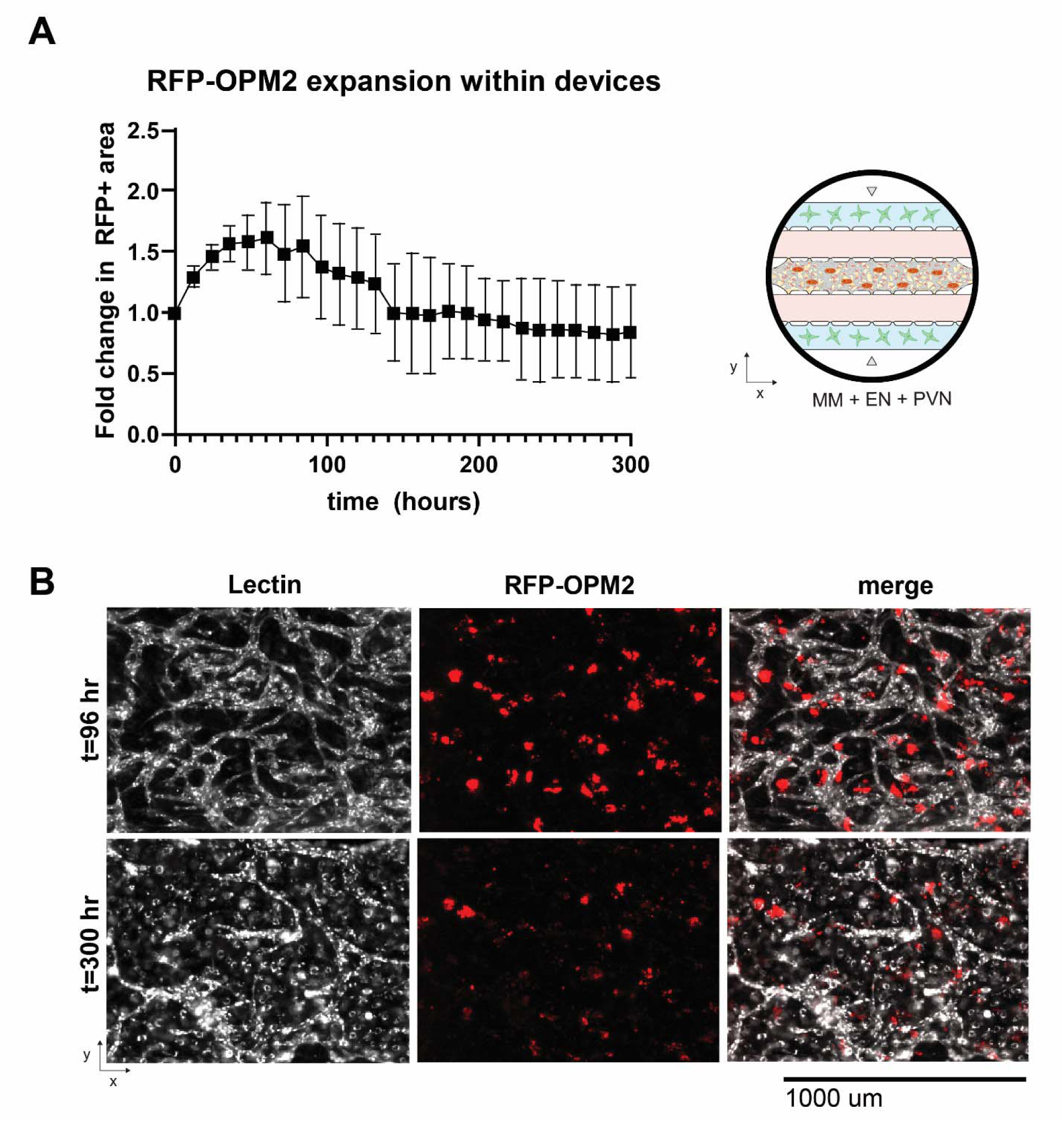
Longevity of reporter MM cell culture within hMM-on-a-chip. (A) When RFP-OPM2 was cultured within the hMM-on-a-chip, we were able to track its proliferation for about 12.5 days (schematic on right). (B) Culture could have continued theoretically indefinitely because the MM cells seemed to reach a steady state of survival, but the devices were endpointed because the lectin-labeled vessels seemed to have overgrown into non-patent monolayers from their original network.

**Supplementary Figure 5:**
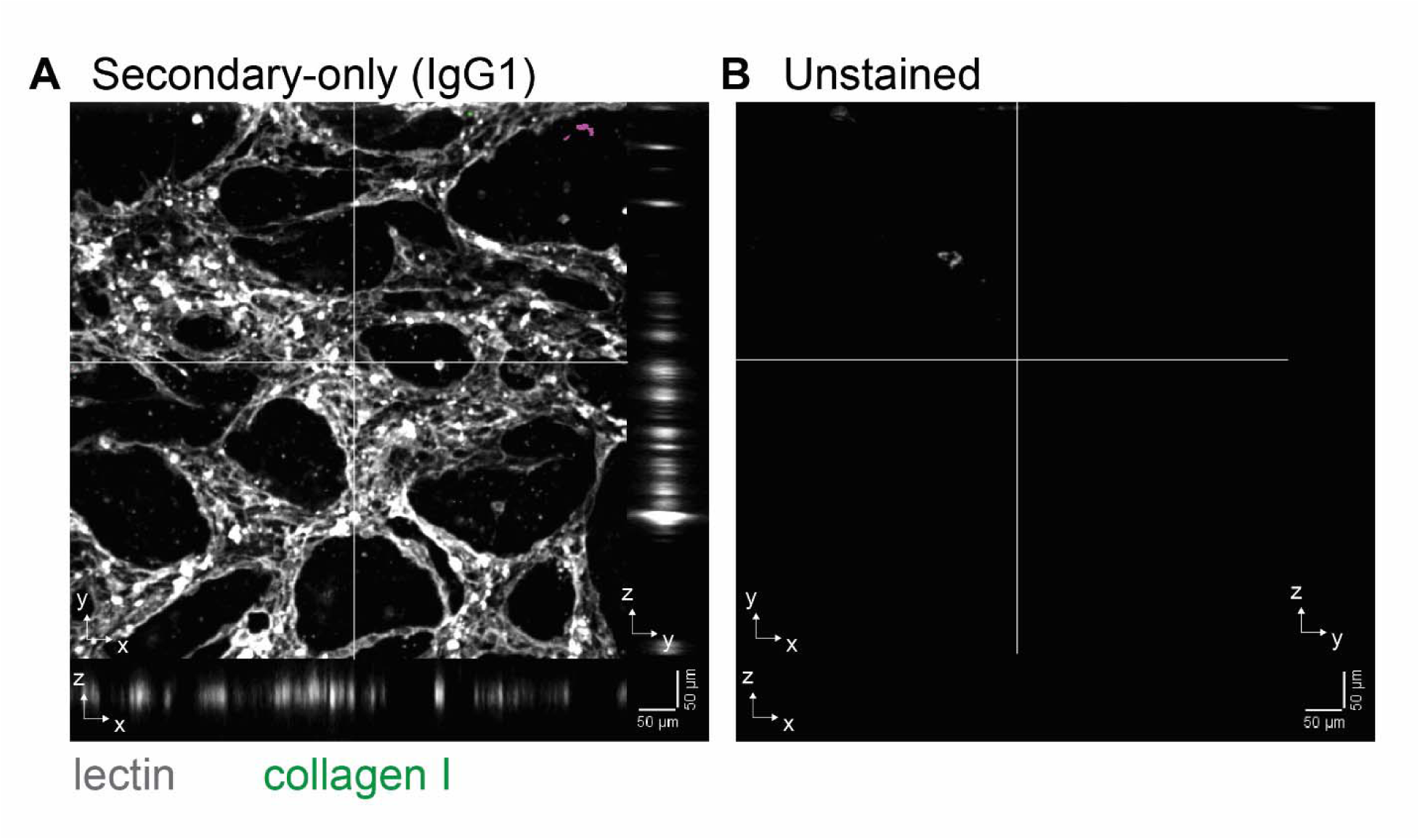
Staining controls for MM localization. (A) Secondary-only staining control and (B) unstained control to demonstrate fidelity of collagen I staining in main figure. Scalebars = 50 µm.

**Supplementary Figure 6:**
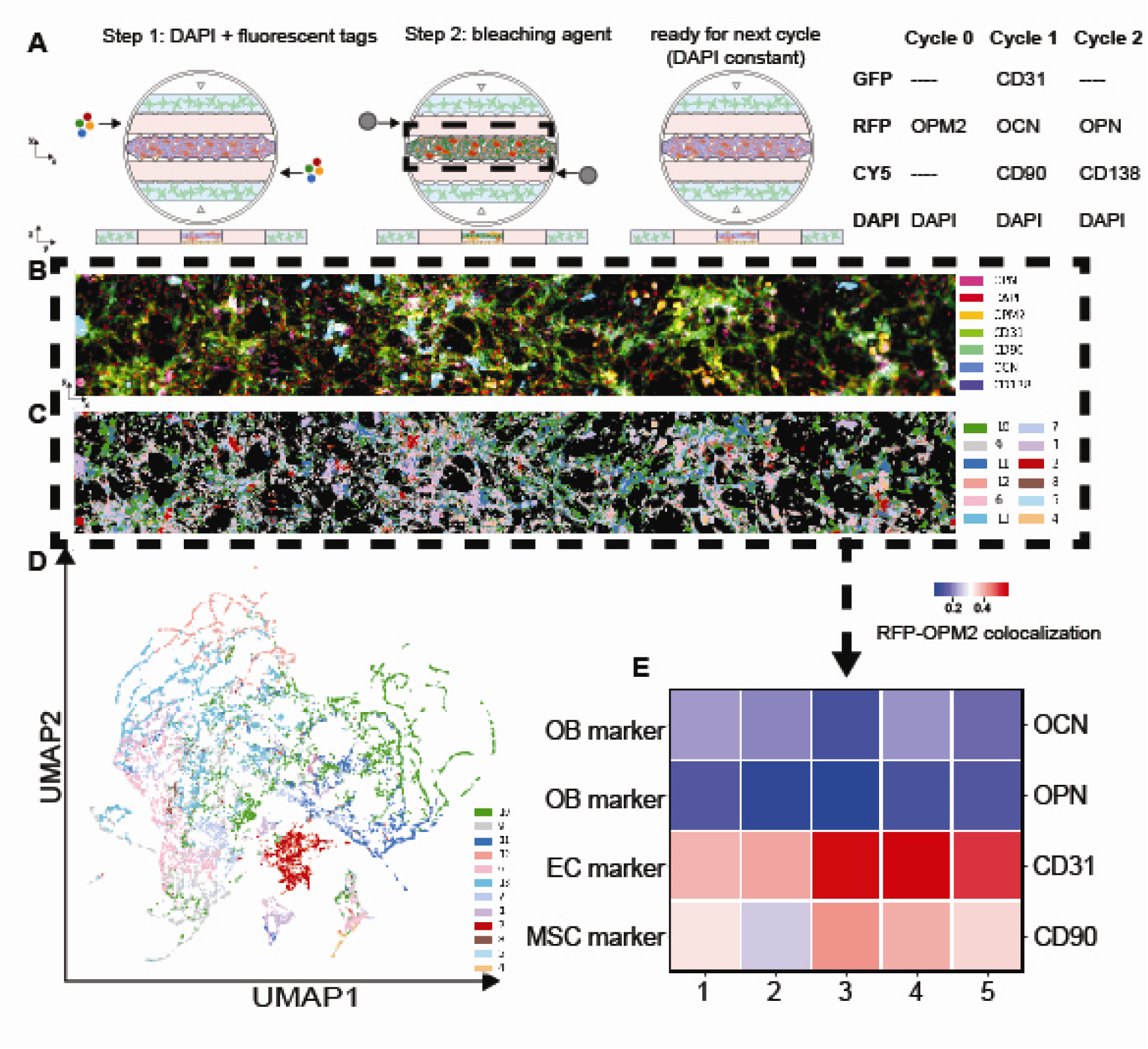
Cyclic immunofluorescence (CycIF) to confirm perivascular colocalization of MM cell lines. (A) Schematic of CycIF workflow, as well as markers tested. (B) Superimposed image of 7 different bone marrow and multiple myeloma markers stained within a representative device. When UMAP analysis was performed on the images, the clustering data was used to color-code the pixels of the image (C). (D) The overall UMAP plot of clustered pixels. (E) The heatmap of representative devices showing relative co-expression of RFP-OPM2 signal with 4 different bone marrow niche markers (rows) within 5 representative devices.(columns) showed significant colocalization of OPM2 MM cells within the perivascular niche of the hMM-on-a-chip.

**Supplementary Figure 7:**
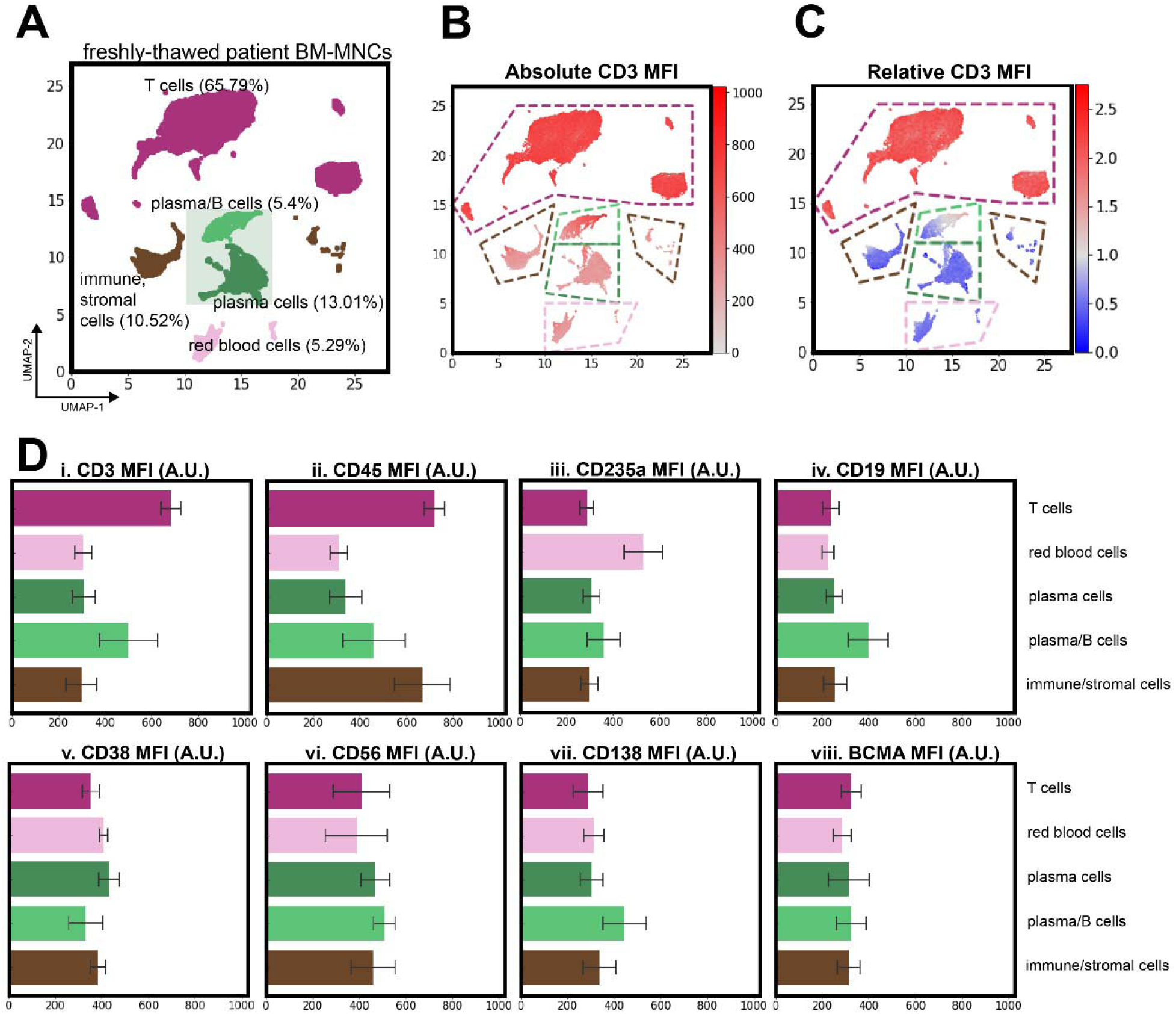
Quantification of patient BM-MNCs prior to introduction to hMM-on-a-chip. (A) the original patient bone marrow mononuclear cells (BM-MNCs) that were cocultured within the hMM-on-a-chip. In order to identify the cell populations within each cluster, the (B) absolute MFI of each flow cytometric marker could be studied, with CD3 shown as an example. We could also (C) normalize the MFI within each cluster in order to measure the differential expression within populations. These observations could be tabulated for multiple antigens as in (D) to justify the cell population labels assigned to each cluster (MFI shown as mean +/- SD within each cluster, with colors corresponding to Panel A). For example, the T cell cluster (purple) was CD3+ CD45+ and the red blood cell cluster (pale pink) was CD235a+. Other cell types were less straightforward to classify; for example, a combination of CD38+ CD56+ CD138+ BCMA+ CD19lo CD45- was needed to classify plasma cells.

**Supplementary Figure 8:**
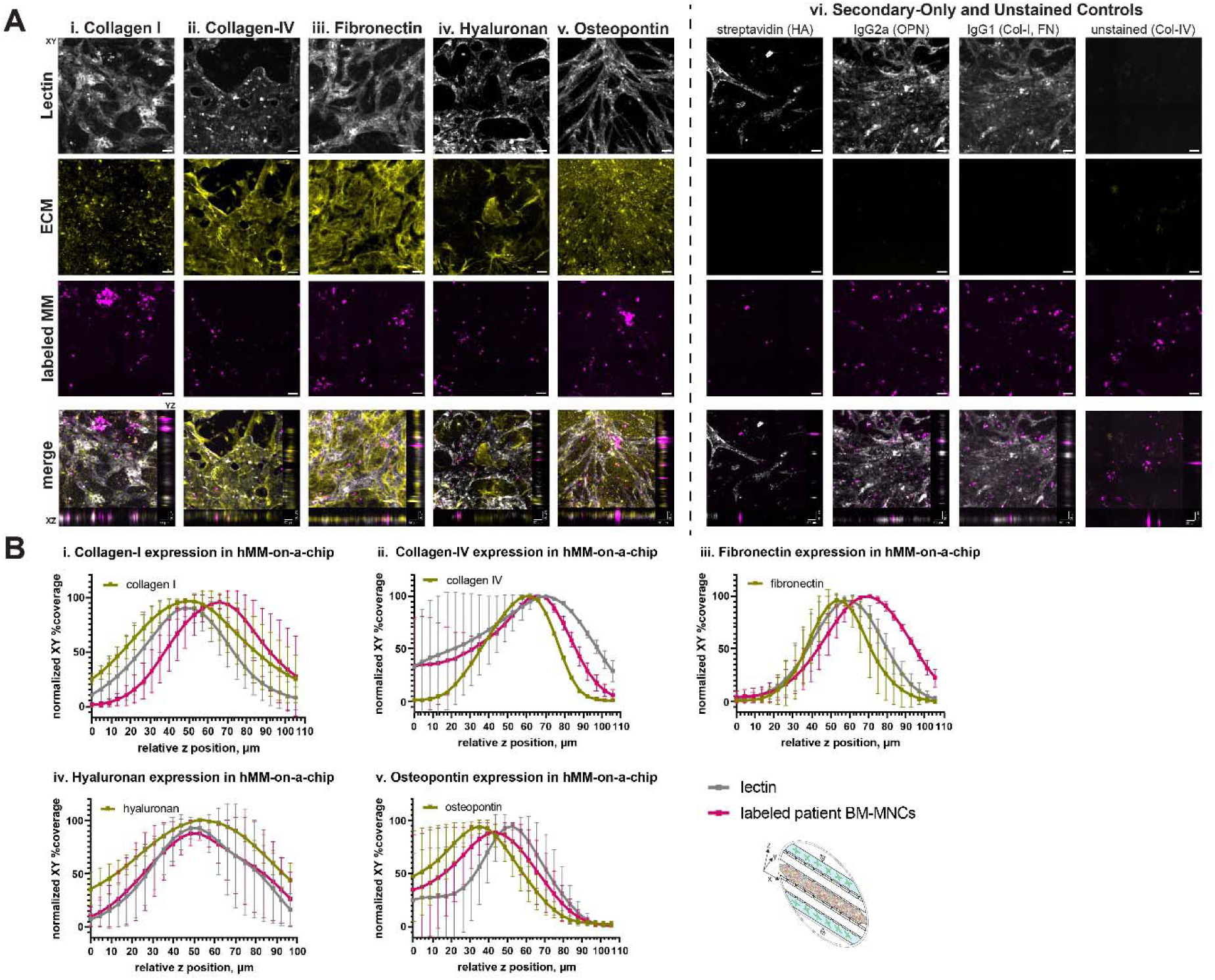
Further quantification of ECM localization within hMM-on-a-chip. (A) Extended-focus Z-stacks of the MM-chip could be used to study the expression of different ECM niche markers (i-v) and their localization with the perivascular niche, as represented by the lectin signal tagging the endothelial cell networks. Each panel is the XY plane of the device in a single channel and the merged image has the XZ and YZ planes focused on a particular ROI. (vi) unstained and secondary-only images that were brightness- and contrast-adjusted the same way as the fully-stained images. Scalebars = 50 µm. (B) the relative percent coverage at each Z slice of n=3 devices could then be plotted for each channel of the signal (lectin, labeled patient BM-MNCs, and the antigen of interest) to further quantify whether the signal was endosteal or perivascular in its location. Endosteal markers, as expected, were (i) collagen I, (iii) fibronectin, and (v) osteopontin. (iv) Hyaluronan could be found throughout, as MSCs were loaded in both niches of the hMM-on-a-chip, and (ii) collagen IV was localized to the perivascular niche. Schematic of device as viewed in 3D provided to orient the reader.

**Supplementary Figure 9:**
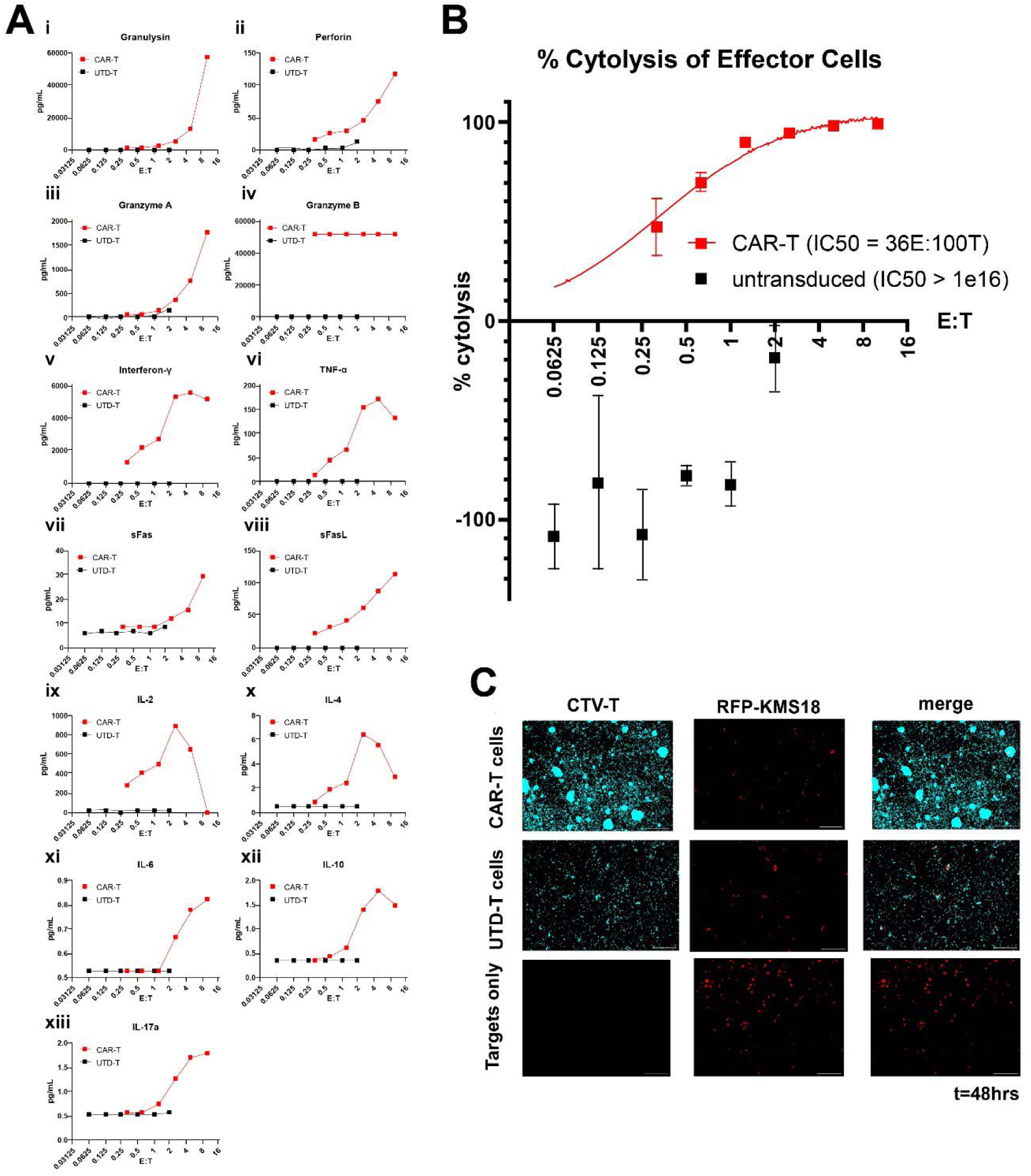
2D cytotoxicity results for CAR-T cells from a healthy donor. (Ai-xiii) 13-plex LEGENDplex detection of T cell activation cytokines from the coculture of CAR-T cells (red traces) or UTD-T cells (black traces) after 48 hours of coculture with RFP-KMS18 target cells at a variety of E:T ratios, with the estimated IC50 listed for each effector type. (B) Flow cytometric analysis of the percentage of these target cells remaining after this 48-hour coculture. (C) representative images of effector and target cells after 48 hours; scale bar =100 μm.

**Supplementary Figure 10:**
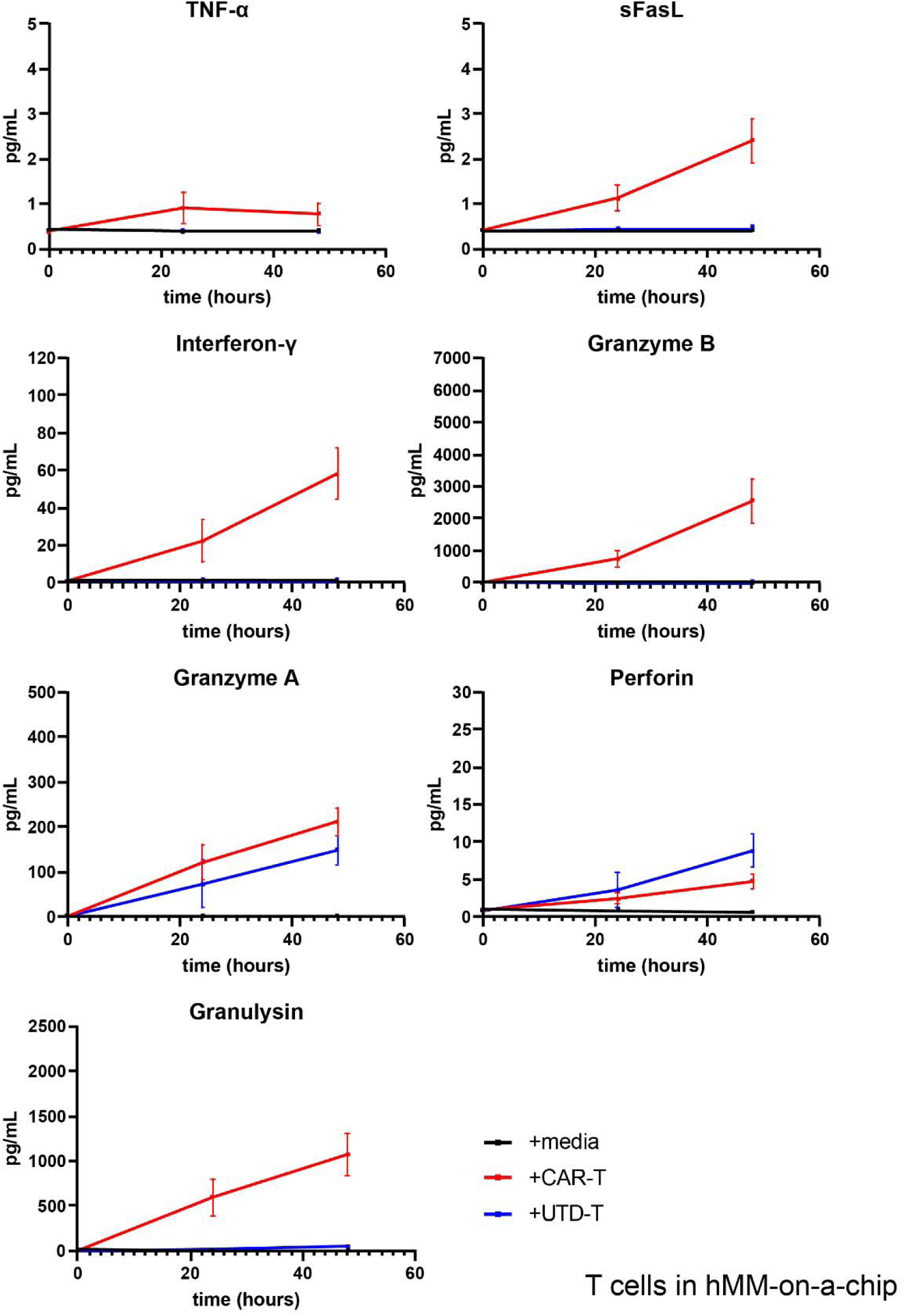
Cytokine release data within hMM-on-a-chip. Timecourses of cytokine release within treated hMM-on-a-chip devices (n=3-4) after 0, 24, and 48 hours of treatment with effectors.

**Supplementary Figure 11:**
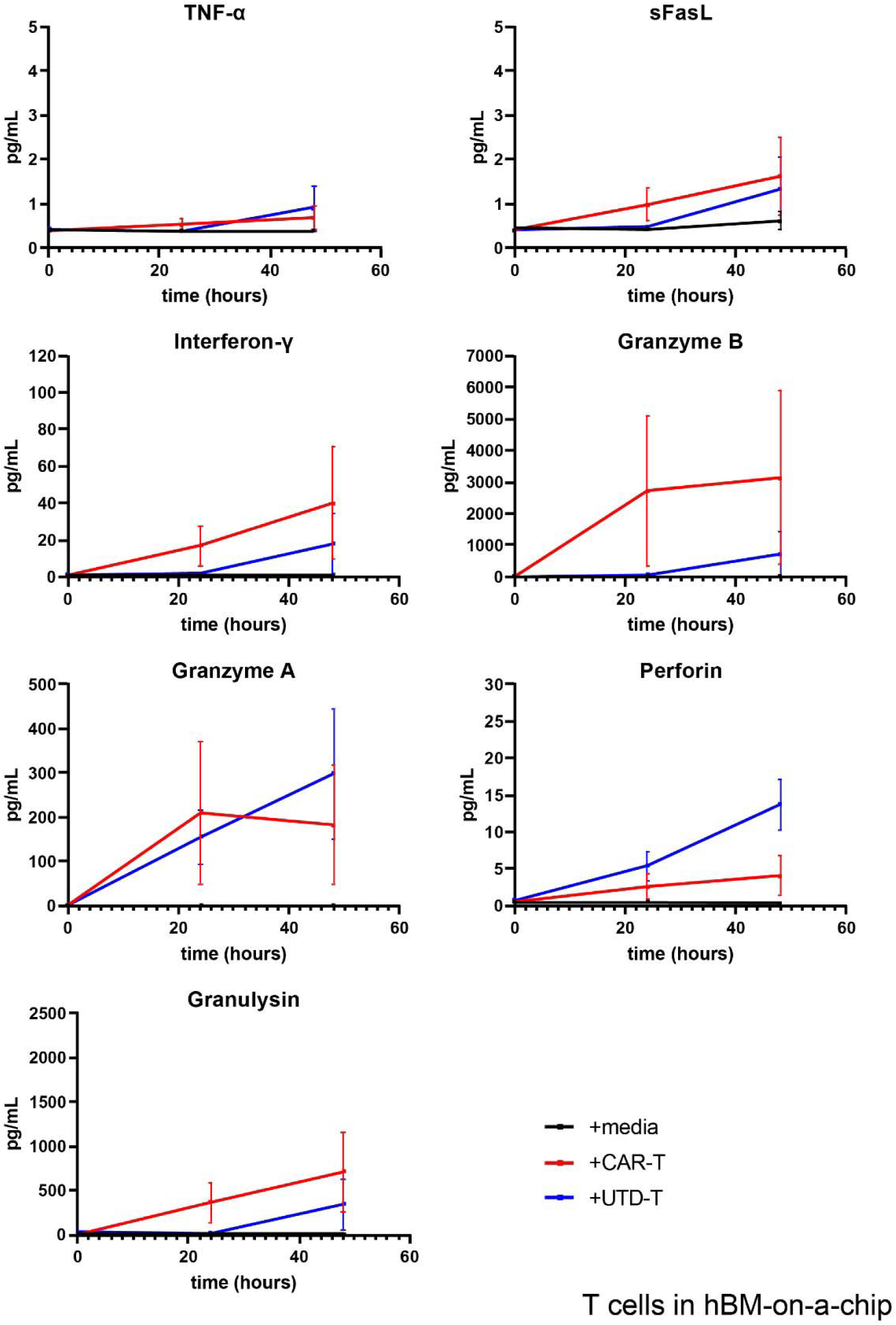
Cytokine release data within hBM-on-a-chip. Timecourses of cytokine release within treated hBM-on-a-chip devices that do not contain any target cells (n=3-4) after 0, 24, and 48 hours of treatment with effectors.

**Supplementary Figure 12:**
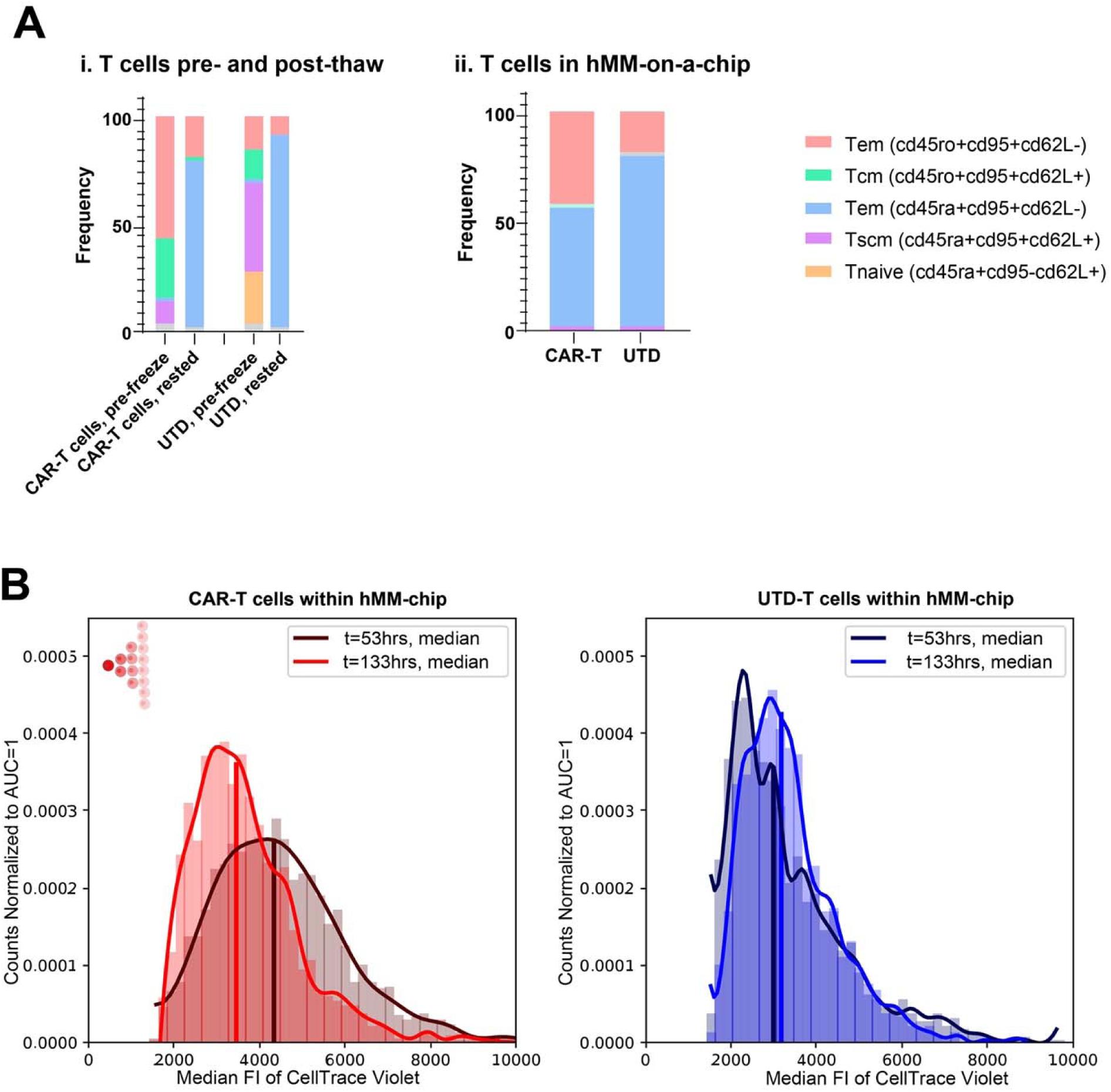
T cell characterization within hMM-on-a-chip treated with healthy-donor T cells. (A) Subsets of T cells (i) before and after cryopreservation, as well as (ii) relative abundances after endpoint within devices. (B) Comparison of MFI for individual labeled T cells at the initiation and termination of culture, to investigate whether the cells divided and lost their fluorescent signal (left inset schematic). n=3-4 devices per condition.

**Supplementary Figure 13:**
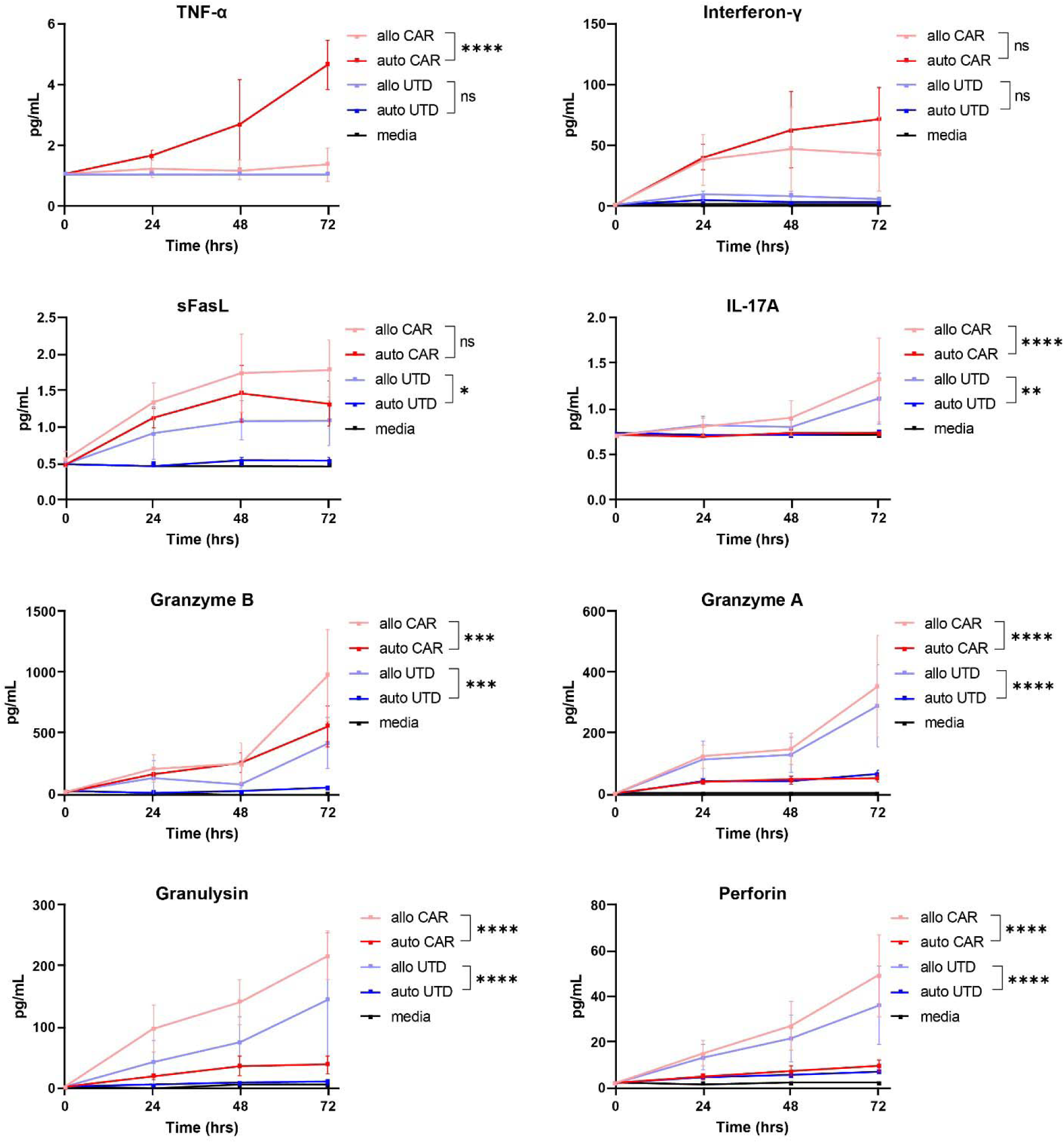
Graft-versus-host cytokine release within hMM-on-a-chip. Raw concentrations of cytokines tracked over time between allogeneic and autologously-introduced CAR- and UTD-treated devices. 2-way repeated measures ANOVA with Tukey post-hoc test, with stars indicating significance of final timepoint between on- and off-target cells. * = p ≤ 0.05, ** = p ≤ 0.01, *** = p ≤ 0.001, **** = p ≤ 0.0001.

**Supplementary Figure 14:**
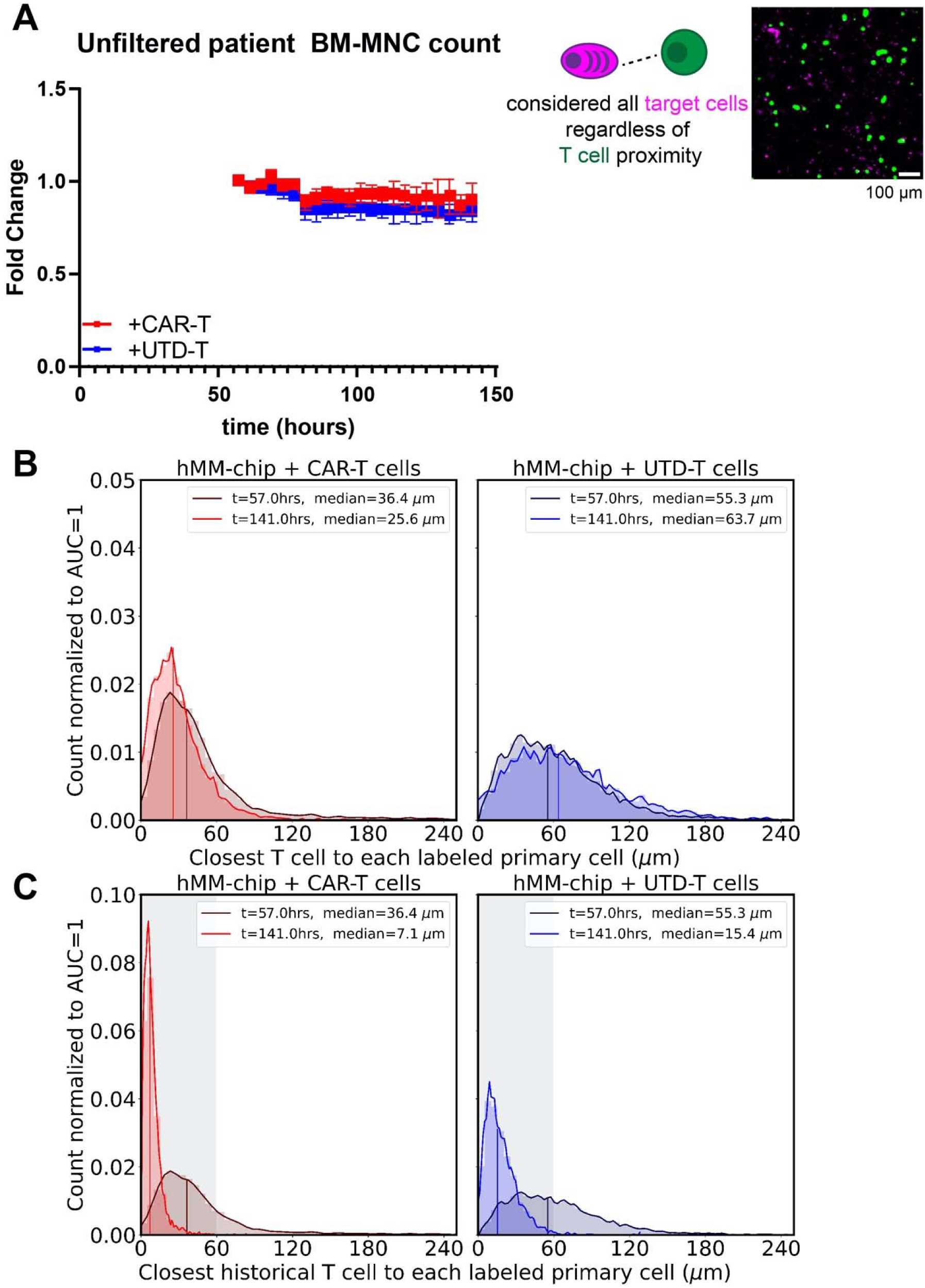
Justification for distance filtering. (A) After CellTrace Violet-labeled CAR-T cells (red trace) or UTD-T cells (blue trace) were introduced into the hMM-on-a-chip, the number of red-fluorescent cell objects did not significantly differ between effector conditions without distance filtration of the data. 2-way repeated-measures ANOVA, n=3-4 devices per condition. (B) Normalized histogram of the nearest T cell distances for each labeled primary cell for each timepoint. (C) Normalized histogram of the nearest historical T cell distances for each labeled primary cell for all cumulative timepoints. Shaded box indicates the 60 µm threshold applied to data depicted in the main figure.

**Supplementary Figure 15:**
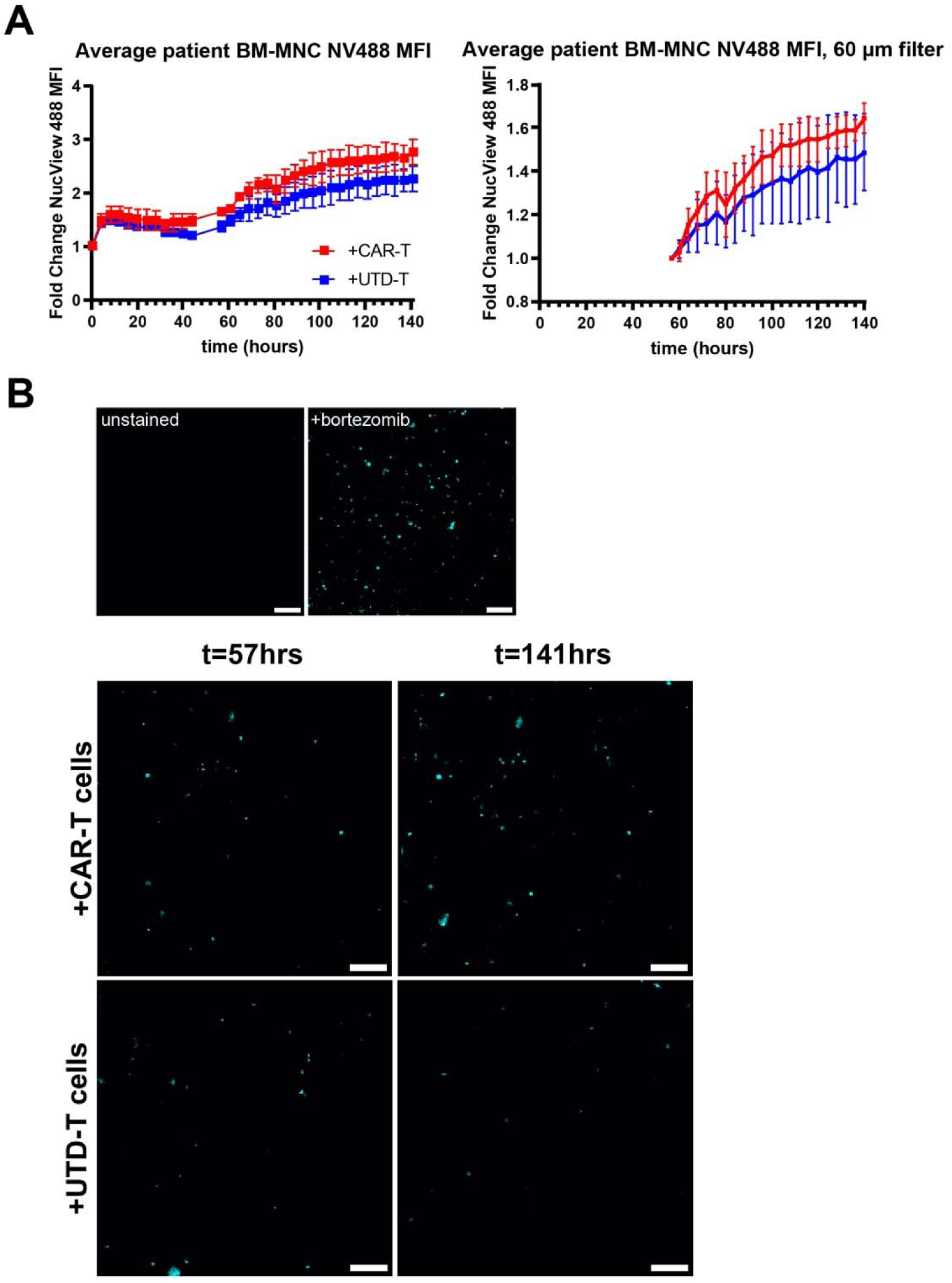
Results from using real-time caspase-sensitive dye to track target cell death. (A) After CellTrace Violet-labeled CAR-T cells (red trace) or UTD-T cells (blue trace) were introduced into the hMM-on-a-chip about 50 hours after initiation of culture within the BioSpa, the MFI of NucView 488 within labeled target cell objects did not significantly differ between effector conditions for either (i) raw data or (ii) distance-filtered data. n=3-4 devices per condition. (B) Representative images of NucView 488 data that was quantified in part (A). Top: signal from unstained device and device treated with bortezomib, an apoptosis inducer. Bottom: NucView 488 signal at beginning of T cell culture (t=57 hours) and at culture endpoint (t=141hrs)

**Supplementary Figure 16:**
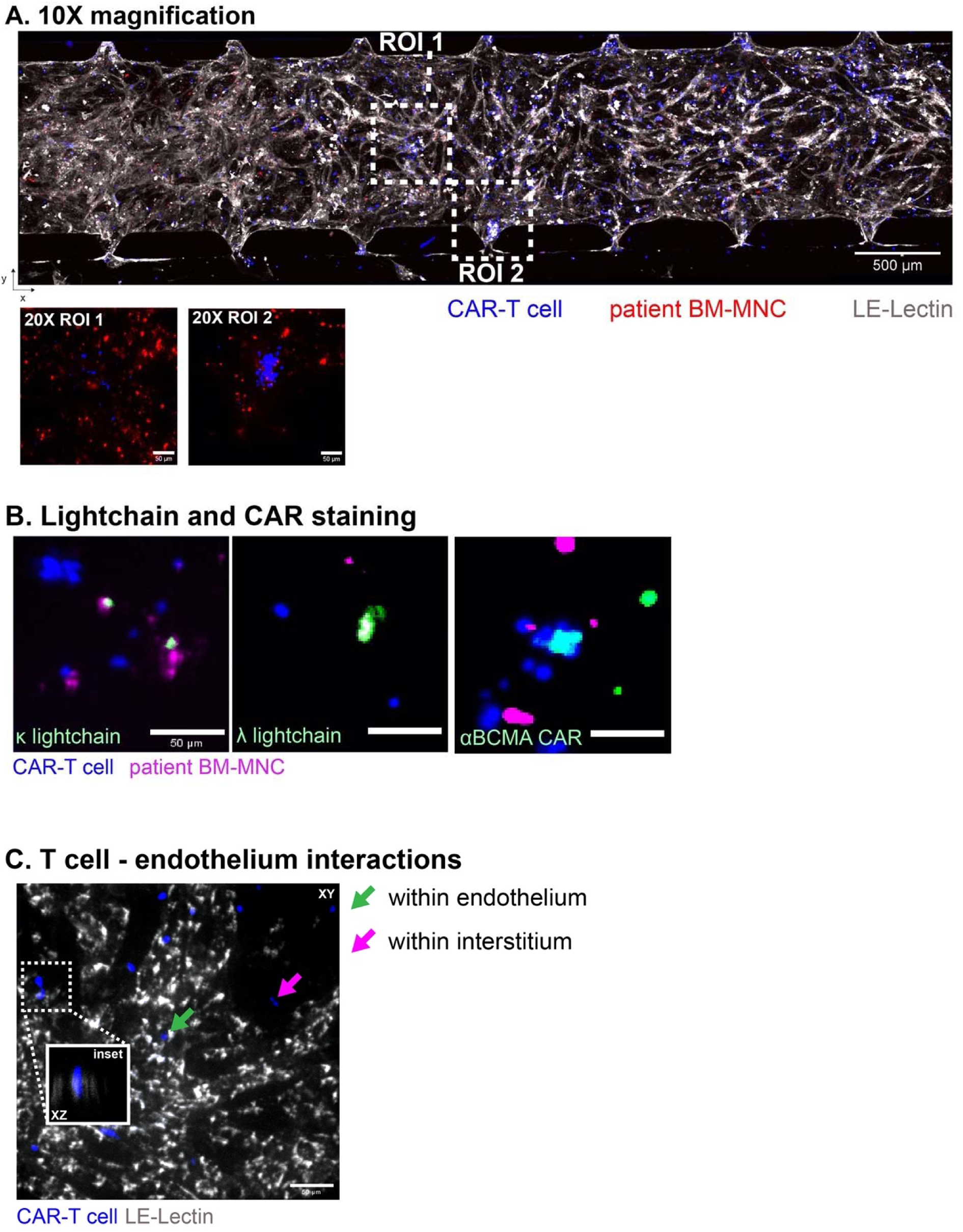
Microscopy of T cell and MM cell markers within hMM-on-a-chip. (A) CAR-T cells could be seen in 10X entering through each pillar opening and mostly remaining within the boundaries of lectin-labeled endothelium. When 20X images were captured (insets, lectin signal omitted for ease of viewing). (B) Labeled CAR-T cells (blue) interacting with labeled BM-MNCs (magenta) stained for κ and λ light-chain+ plasma cells (left, middle) or the anti-BCMA CAR molecule on the T cell (right). (C) T cells flow into vasculature and home to various regions within the vessels or in the interstitial space. They can travel from within the vessels to the interstitium by way of extravasation through the endothelium wall (20X inset).

**Supplementary Figure 17:**
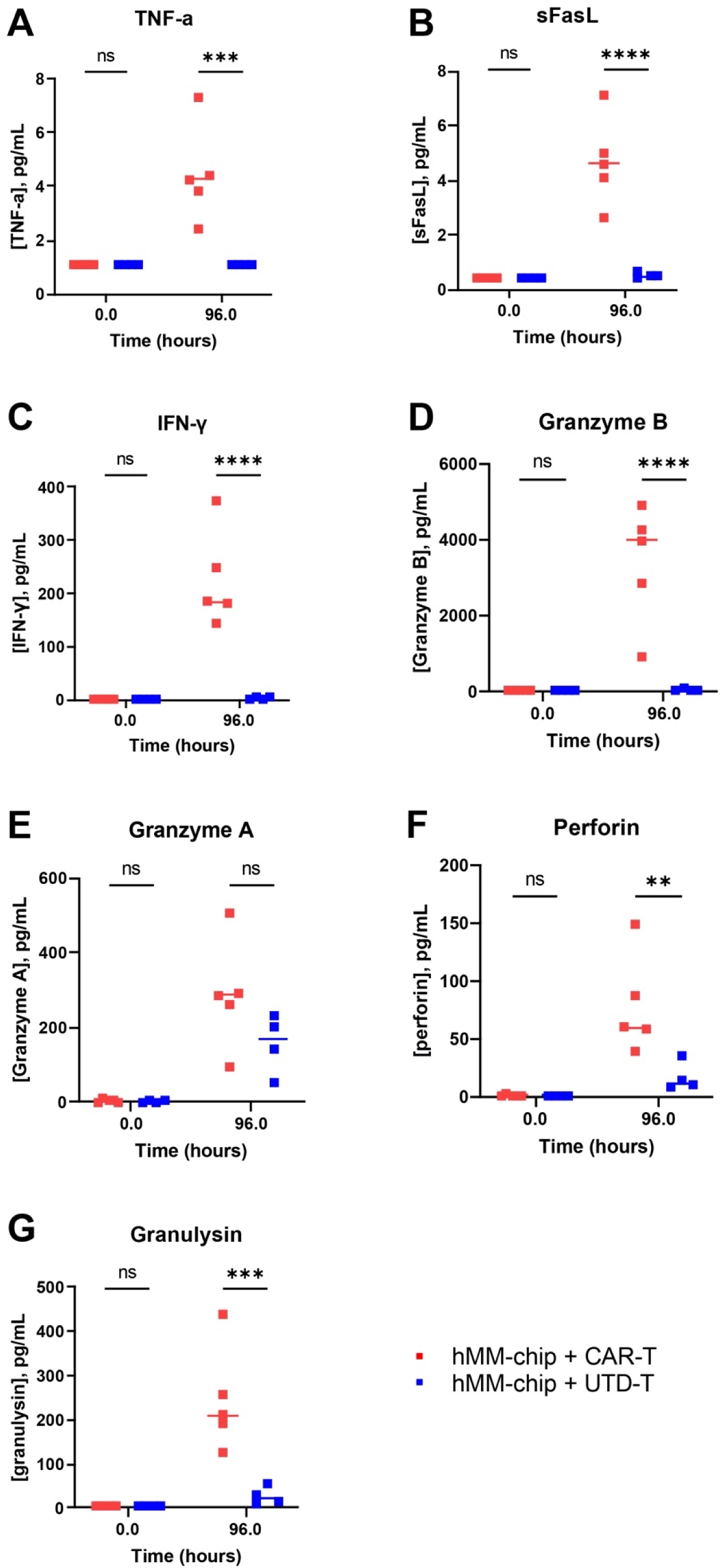
Raw cytokine release values for hMM-on-a-chip with primary BM-MNC targets. Welch’s non-parametric t-test with Dunnett T3 post-hoc tests, ** = p ≤ 0.01, *** = p ≤ 0.001, **** = p ≤ 0.0001. The CAR-T-specific cytokines, as before, were (A) TNF-α, (B) sFasL, (C) IFN-γ, and (D) Granzyme B. Non-specific cytokines released by both cell types were (E) Granzyme A, (F) Perforin, and (G) Granzyme A.

