## Supplementary Materials for "Multi-Niche Human Bone Marrow On-A-Chip for Studying the Interactions of Adoptive CAR-T Cell Therapies with Multiple Myeloma"

***Science* *Advances* Supplementary Materials Template Instructions**

This is the *Science Advances* template for presenting and formatting your supplementary materials. To organize your supplementary materials section, please follow the instructions below. Once formatted, you should delete this first page of instructions.

**Overview:**

Supplementary Materials present additional information in support of the conclusions of your paper, such as a description of the materials and methods, controls, or tabulated data presented in Tables or Figures. It will consist of one PDF file with embedded figures and tables (if needed). Audio or movie files or large data Tables can be presented as separate files. See the [*Science Advances* website](https://advances.sciencemag.org/content/information-authors) for detailed instructions.

Supplementary Materials should not be used for additional discussion, analysis, or interpretations. It is not to be used as a forum to critique other publications.

References can be cited in the Supplementary Text section. These should be cited in order following the references in the main text as per *Science Advances* style (i.e., italicized number in parentheses). Include Supplementary Materials references in the full reference list at the end of the main paper.

**Using the Template**

Paste the title, first author full name (see examples for format), and corresponding author email address(es) from the main text file onto the cover page. On the cover page, complete the relevant table of contents of the SM and delete text that does not apply.

Copy and paste relevant text into each appropriate section of the template. For consistency use Times, 12 pt. Left-align all paragraphs, separating each paragraph by a line-break.

Each figure or table should be on a separate page and can be placed above each caption. To add additional captions, simply copy and paste (repeatedly) the last caption template. Large tables that extend beyond the width of the page should be provided as separate files in an appropriate spreadsheet format (.xlsx or similar).

Large amounts of text can be grouped by subheads. To repeat subheads, simply copy the subhead and repeat/rename.

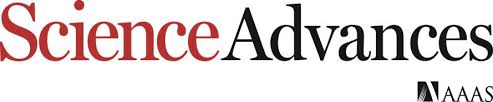

Supplementary Materials for

**Multi-Niche Human Bone Marrow On-A-Chip for Studying the Interactions of Adoptive CAR-T Cell Therapies with Multiple Myeloma**

Delta Ghoshal *et al.*

**This PDF file includes:**

Figs. S1 to S17

Tables S1 to S4

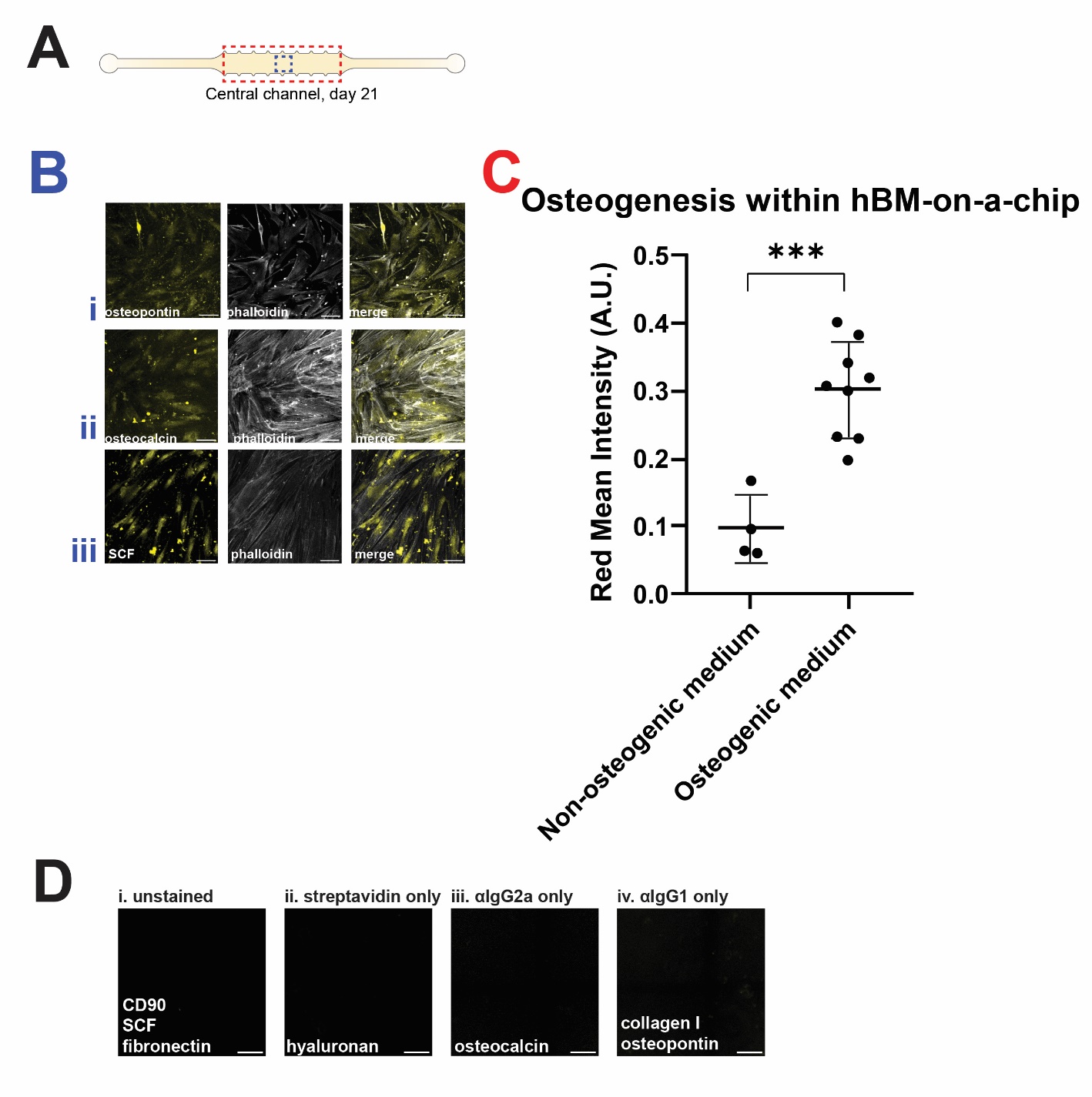

Fig. S1.

Overview of hBM-on-a-chip and hMM-on-a-chip. (A) cartoon depiction of the central channel, where analyses were conducted. The blue dashed box signifies approximately where (B) extended-focus confocal images of (i) osteopontin, (ii) osteocalcin, and (iii) stem cell factor, counterstained for actin, were collected (representative images). The red dashed box signifies where (C) alizarin red was used to label calcium deposits left in devices by the MSC-derived osteoblasts (n=9) or undifferentiated MSCs (n=4). These deposits were quantified for their red intensity in AU. *** = p ≤ 0.001, Welch's non-parametric t-test. (D) IHC control images for (i) directly-conjugated antibodies (CD90, SCF, and fibronectin), (ii) biotinylated HA binding protein, and unconjugated antibodies against (iii) osteocalcin and (iv) collagen I and osteopontin.

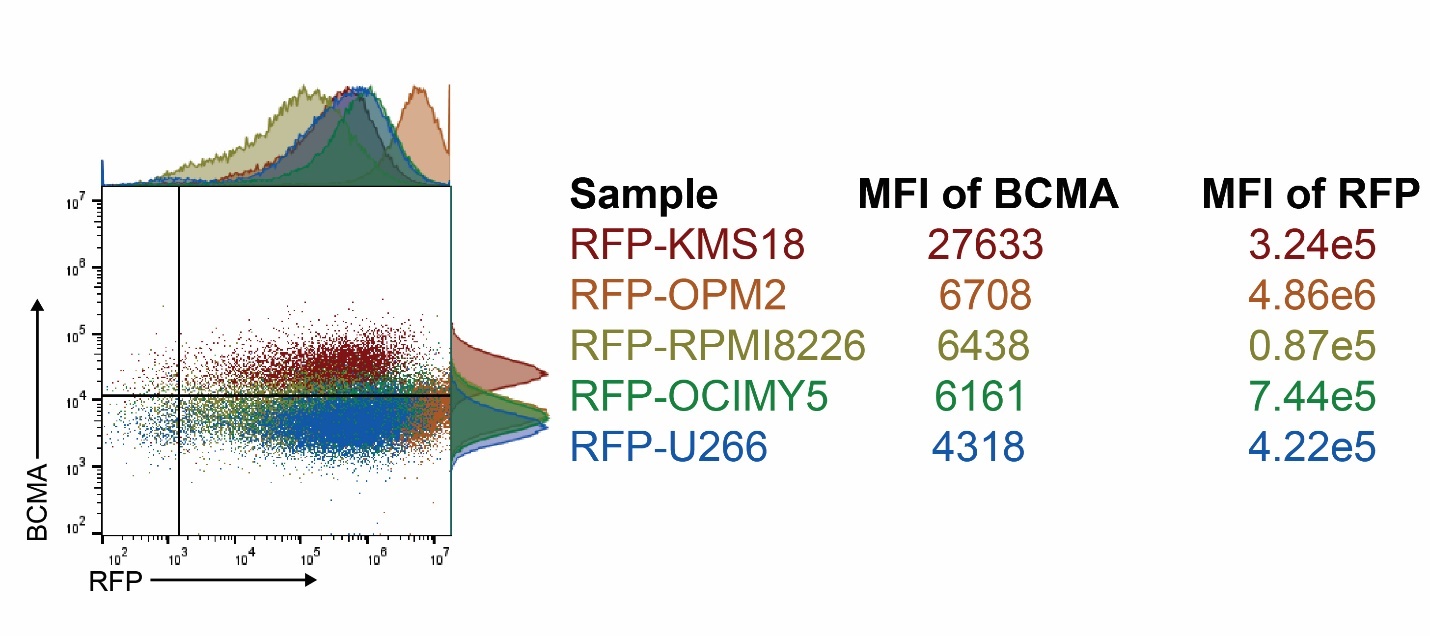
Fig. S2.

Characterization of reporter MM cell lines. (A) Flow cytometric analysis of RFP expression plotted versus BCMA expression showed varying, albeit detectable, degrees of expression.

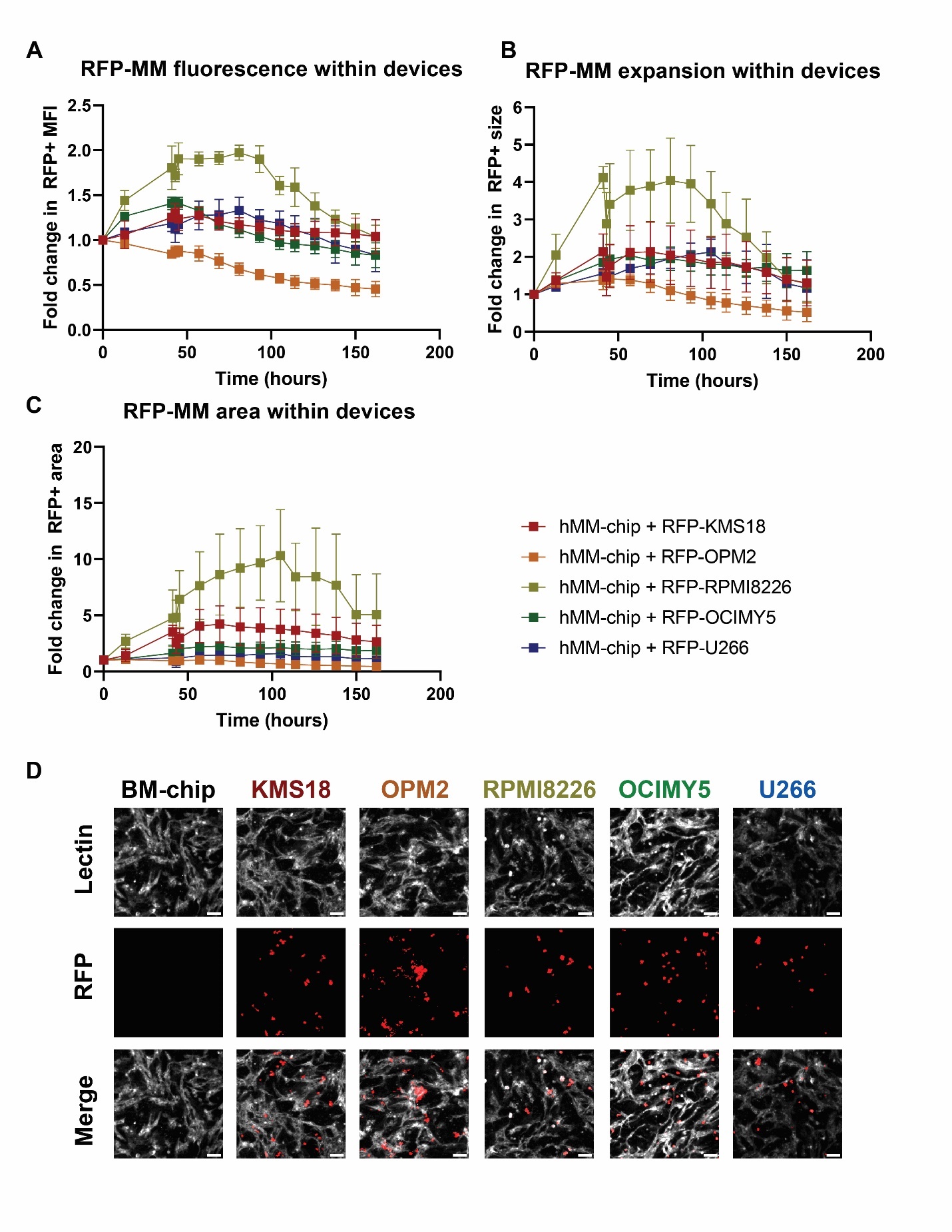
Fig. S3.

Further quantification of MM reporter lines within hMM-on-chip. All figure axes were chosen to match those in the main figure. (A) The relative MFI of MM objects for 4 of the cell lines did not decrease; OPM2's loss of MFI could indicate the cells were not surviving well within the system, as supported by the proliferation data. (B) Average object size and (C) Number of objects for the 5 considered cell lines. Because object size increased in several cases while number of objects often remained stable, this suggested potential clonal expansion of the cell lines. This is why area was chosen to quantify MM cell growth within the system. (D) Clonal expansion and perivascular localization for each cell line. Scalebars = 100 µm.

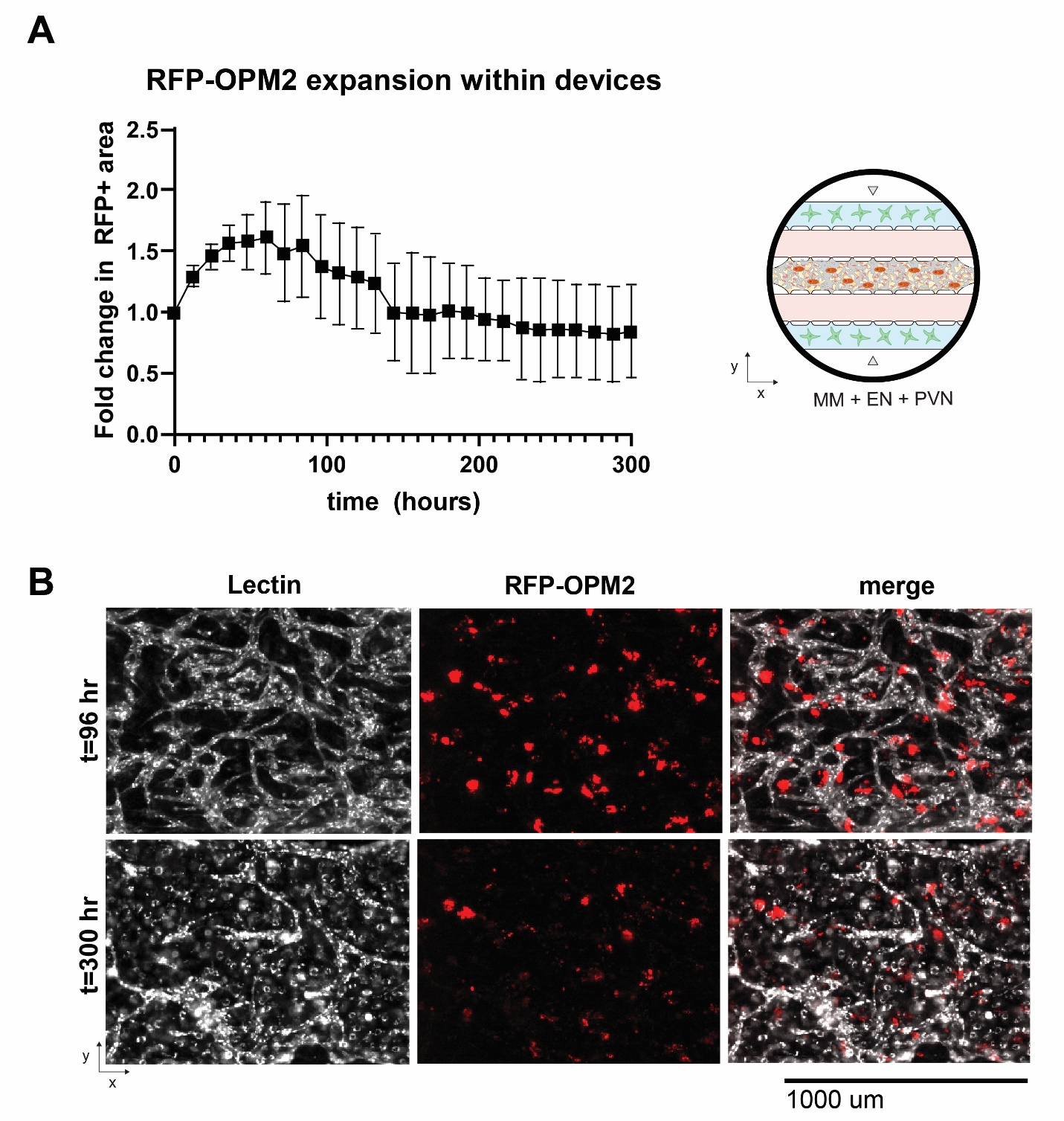
Fig. S4.

Longevity of reporter MM cell culture within hMM-on-a-chip. (A) When RFP-OPM2 was cultured within the hMM-on-a-chip, we were able to track its proliferation for about 12.5 days (schematic on right). (B) Culture could have continued theoretically indefinitely because the MM cells seemed to reach a steady state of survival, but the devices were endpointed because the lectin-labeled vessels seemed to have overgrown into non-patent monolayers from their original network.

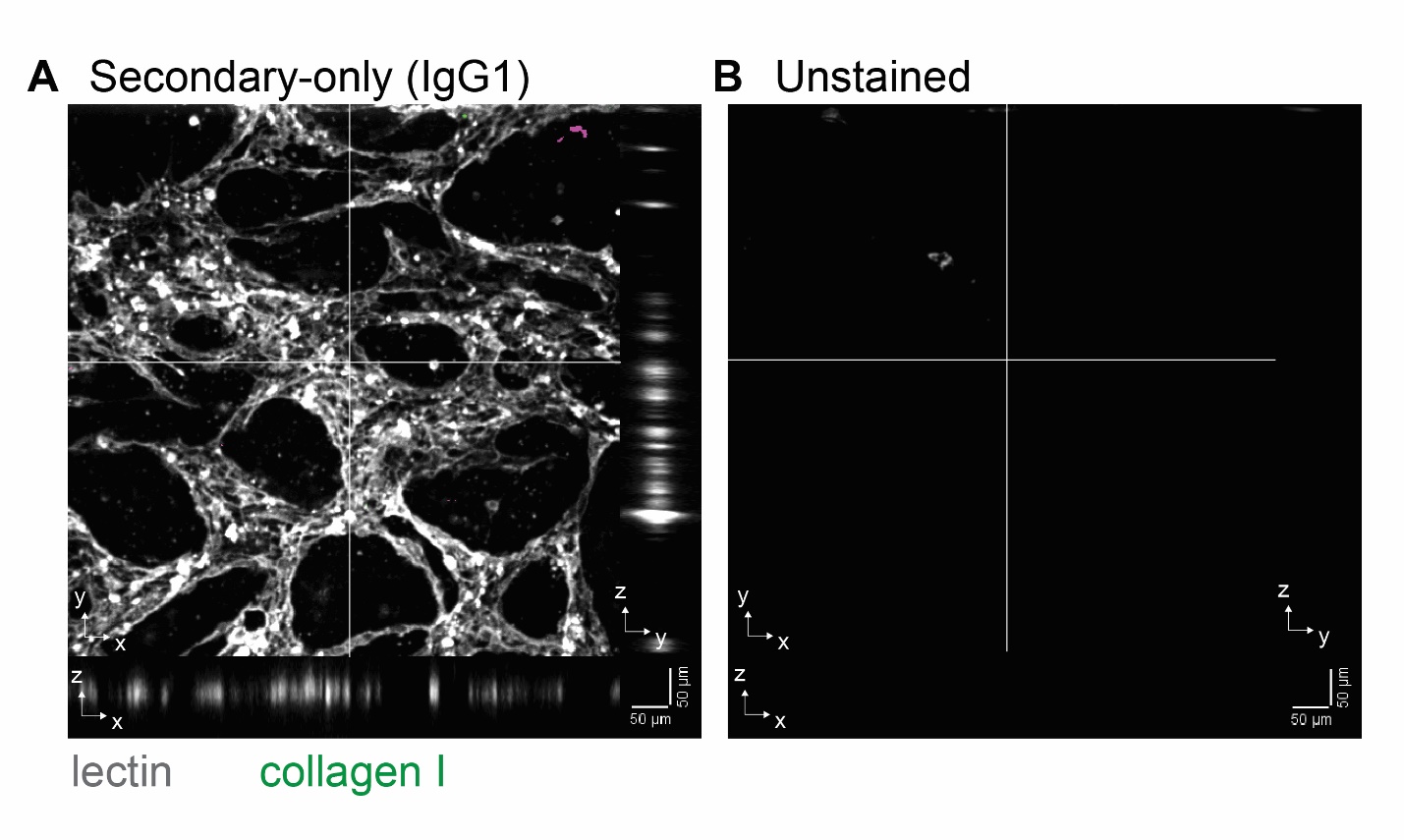
Fig. S5.

(A) Secondary-only staining control and (B) unstained control to demonstrate fidelity of collagen I staining in main figure. Scalebars = 50 µm.

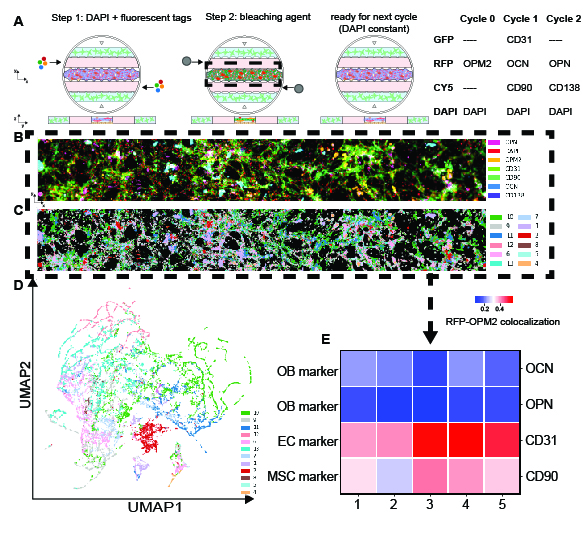
Fig. S6.

Cyclic immunofluorescence (CycIF) to confirm perivascular colocalization of MM cell lines. (A) Schematic of CycIF workflow, as well as markers tested. (B) Superimposed image of 7 different bone marrow and multiple myeloma markers stained within a representative device. When UMAP analysis was performed on the images, the clustering data was used to color-code the pixels of the image (C). (D) The overall UMAP plot of clustered pixels. (E) The heatmap of representative devices showing relative co-expression of RFP-OPM2 signal with 4 different bone marrow niche markers (rows) within 5 representative devices.(columns) showed significant colocalization of OPM2 MM cells within the perivascular niche of the hMM-on-a-chip.

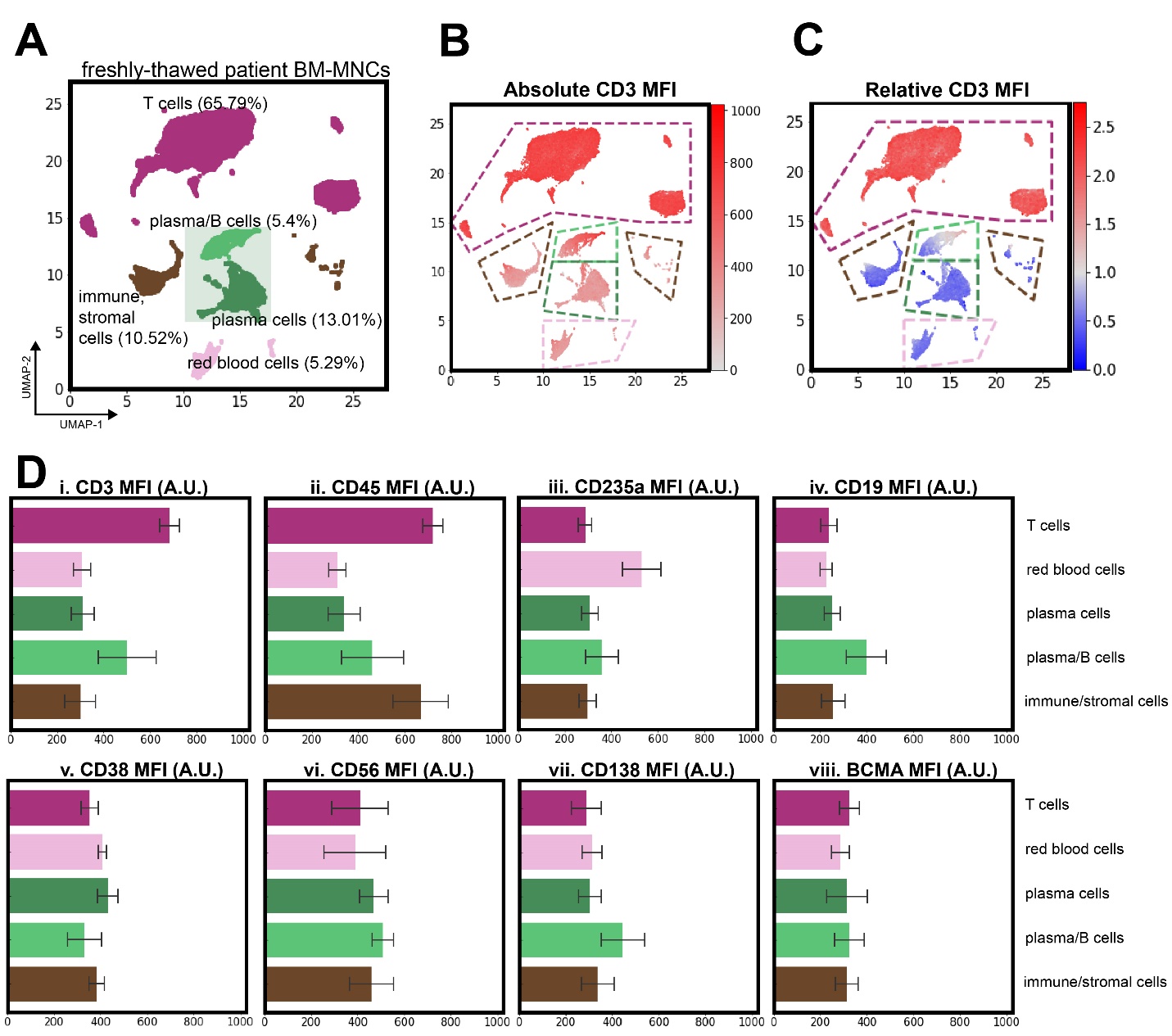
Fig. S7.

Quantification of patient BM-MNCs prior to introduction to hMM-on-a-chip. (A) the original patient bone marrow mononuclear cells (BM-MNCs) that were cocultured within the hMM-on-a-chip. In order to identify the cell populations within each cluster, the (B) absolute MFI of each flow cytometric marker could be studied, with CD3 shown as an example. We could also (C) normalize the MFI within each cluster in order to measure the differential expression within populations. These observations could be tabulated for multiple antigens as in (D) to justify the cell population labels assigned to each cluster (MFI shown as mean +/- SD within each cluster, with colors corresponding to Panel A). For example, the T cell cluster (purple) was CD3+ CD45+ and the red blood cell cluster (pale pink) was CD235a+. Other cell types were less straightforward to classify; for example, a combination of CD38+ CD56+ CD138+ BCMA+ CD19lo CD45- was needed to classify plasma cells.

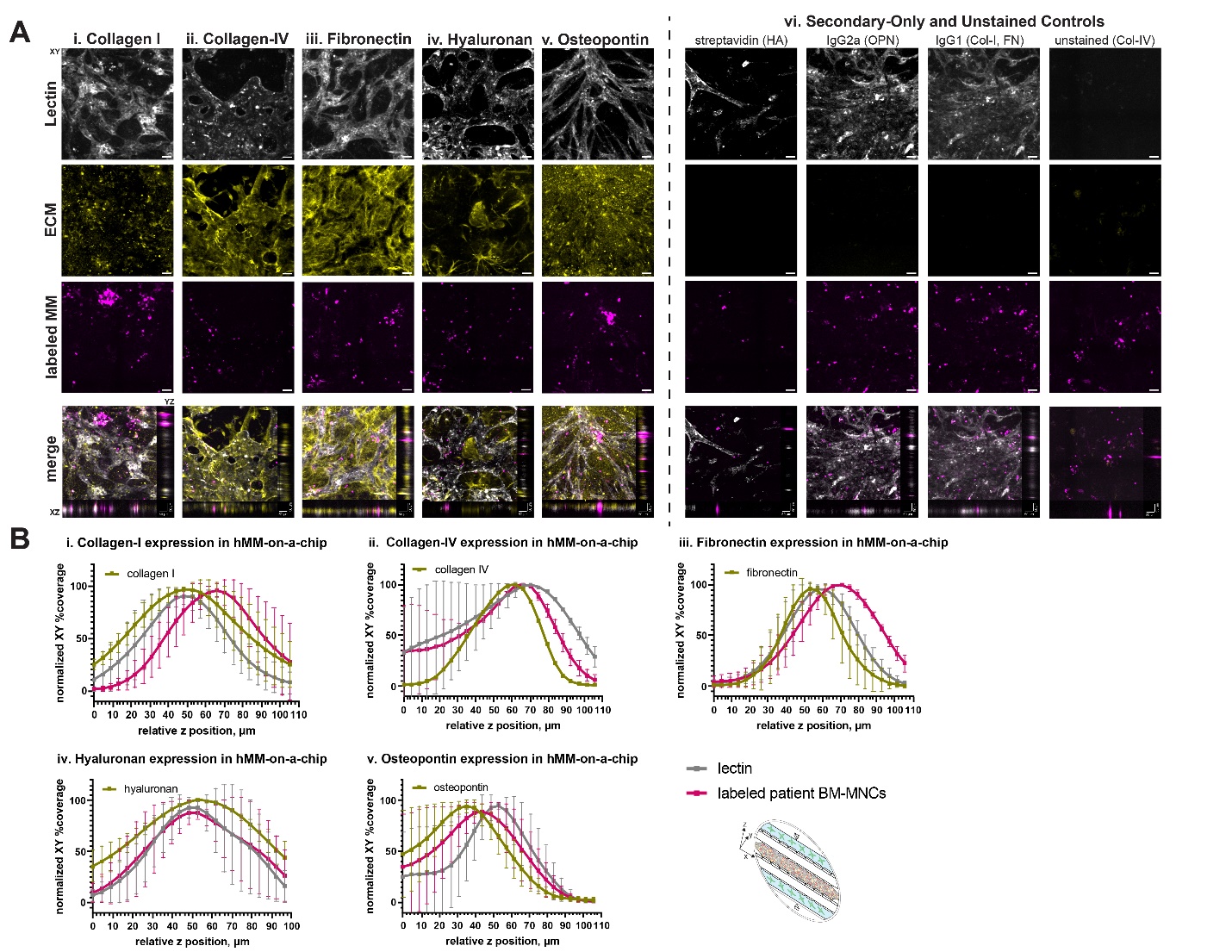
Fig. S8.

Further quantification of ECM localization within hMM-on-a-chip. (A) Extended-focus Z-stacks of the MM-chip could be used to study the expression of different ECM niche markers (i-v) and their localization with the perivascular niche, as represented by the lectin signal tagging the endothelial cell networks. Each panel is the XY plane of the device in a single channel and the merged image has the XZ and YZ planes focused on a particular ROI. (vi) unstained and secondary-only images that were brightness- and contrast-adjusted the same way as the fully-stained images. Scalebars = 50 µm. (B) the relative percent coverage at each Z slice of n=3 devices could then be plotted for each channel of the signal (lectin, labeled patient BM-MNCs, and the antigen of interest) to further quantify whether the signal was endosteal or perivascular in its location. Endosteal markers, as expected, were (i) collagen I, (iii) fibronectin, and (v) osteopontin. (iv) Hyaluronan could be found throughout, as MSCs were loaded in both niches of the hMM-on-a-chip, and (ii) collagen IV was localized to the perivascular niche. Schematic of device as viewed in 3D provided to orient the reader.

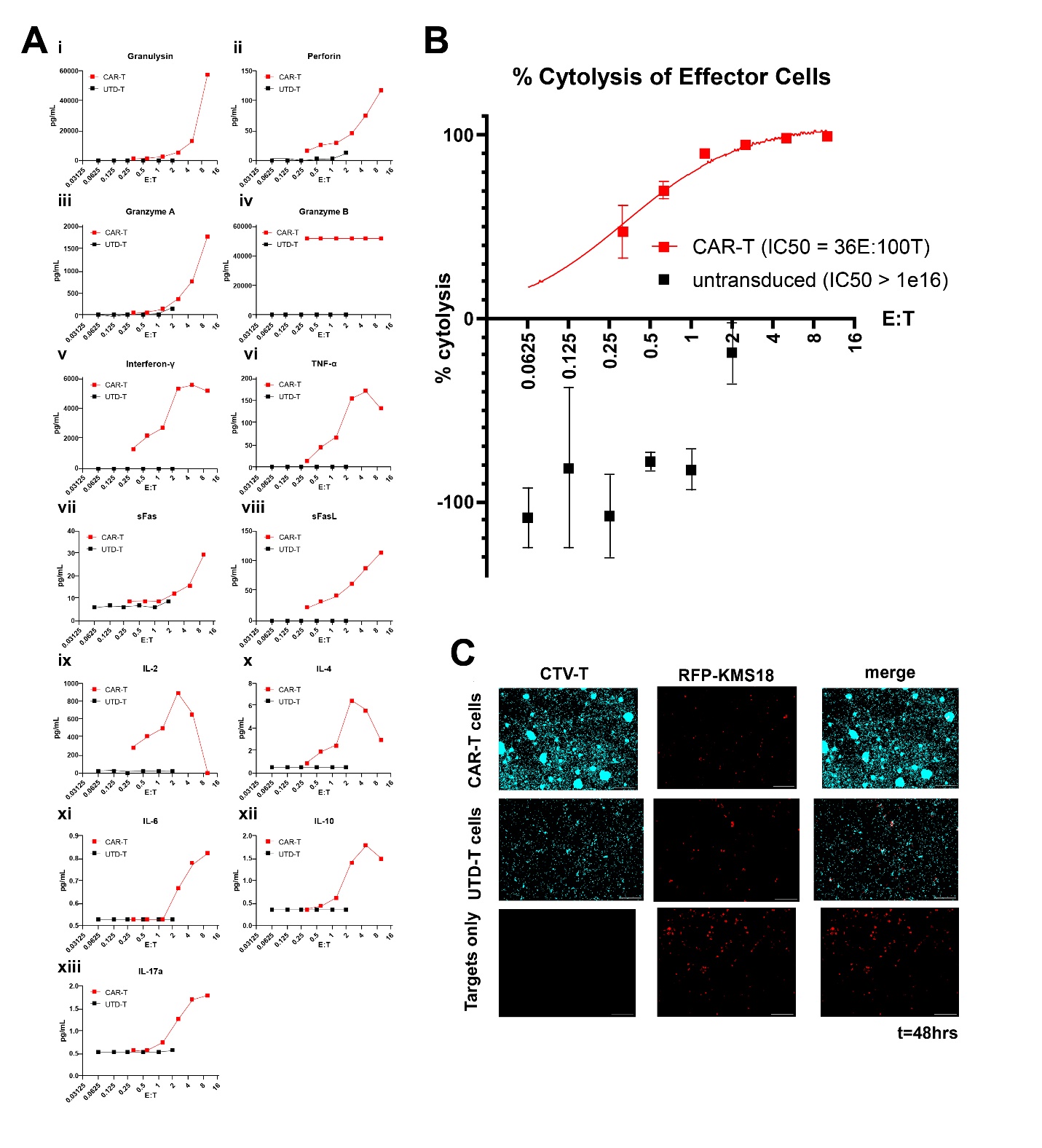
Fig. S9.

2D cytotoxicity results for CAR-T cells from a healthy donor. (Ai-xiii) 13-plex LEGENDplex detection of T cell activation cytokines from the coculture of CAR-T cells (red traces) or UTD-T cells (black traces) after 48 hours of coculture with RFP-KMS18 target cells at a variety of E:T ratios, with the estimated IC50 listed for each effector type. (B) Flow cytometric analysis of the percentage of these target cells remaining after this 48-hour coculture. (C) representative images of effector and target cells after 48 hours; scale bar =100 μm.

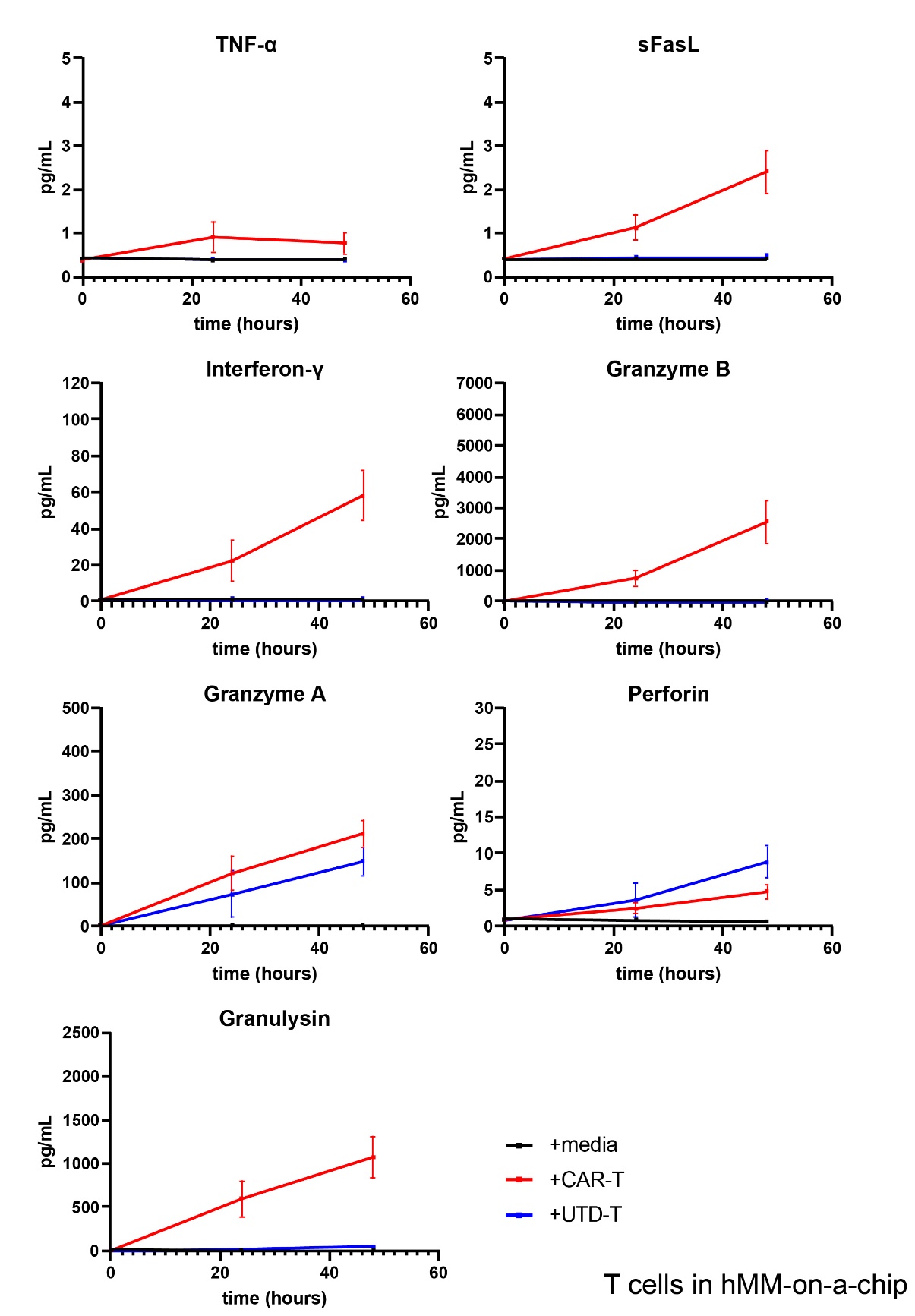

Fig. S10.

Cytokine release data within hMM-on-a-chip. Timecourses of cytokine release within treated hMM-on-a-chip devices (n=3-4) after 0, 24, and 48 hours of treatment with effectors.

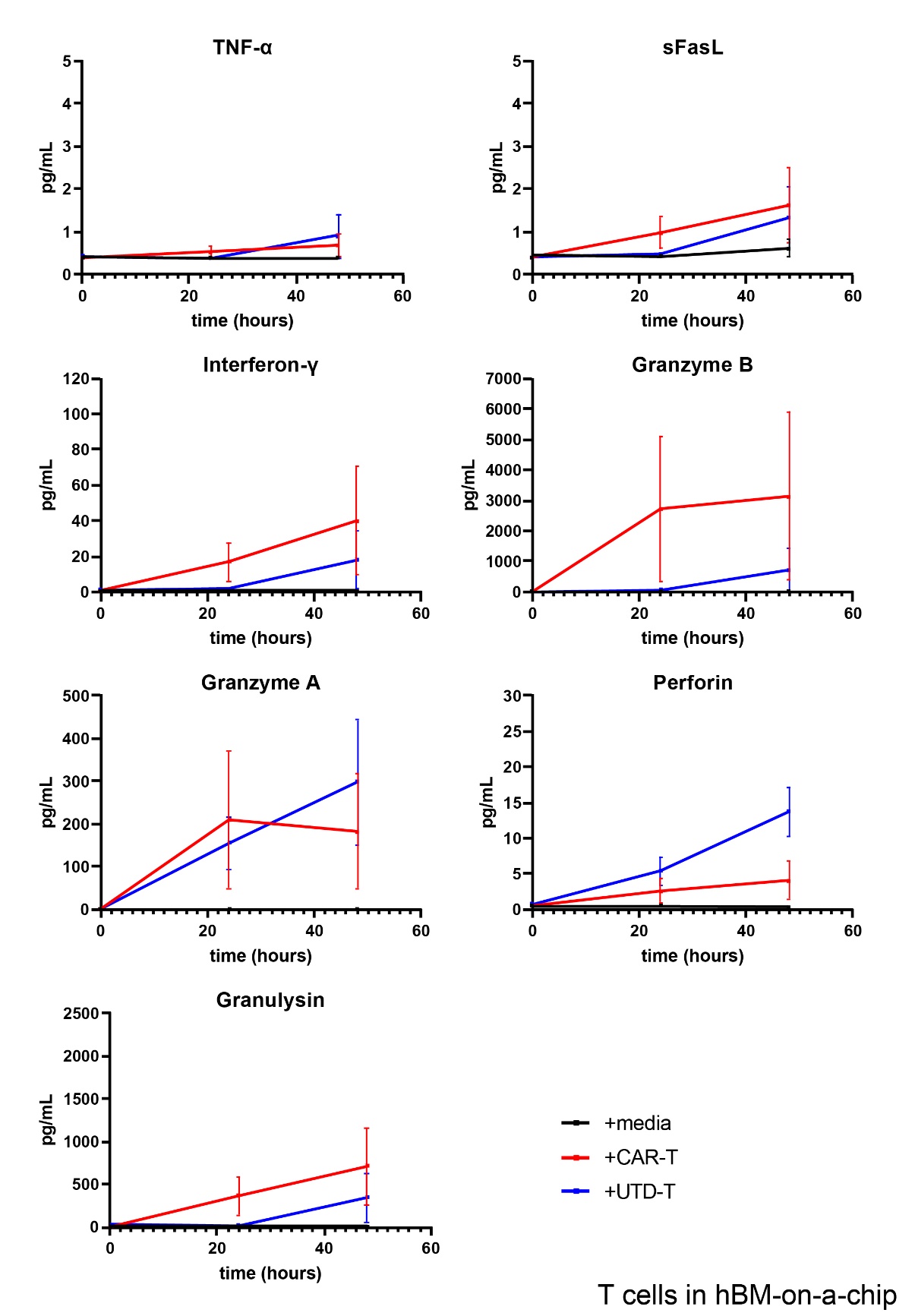

Fig. S11.

Cytokine release data within hBM-on-a-chip. Timecourses of cytokine release within treated hBM-on-a-chip devices that do not contain any target cells (n=3-4) after 0, 24, and 48 hours of treatment with effectors.

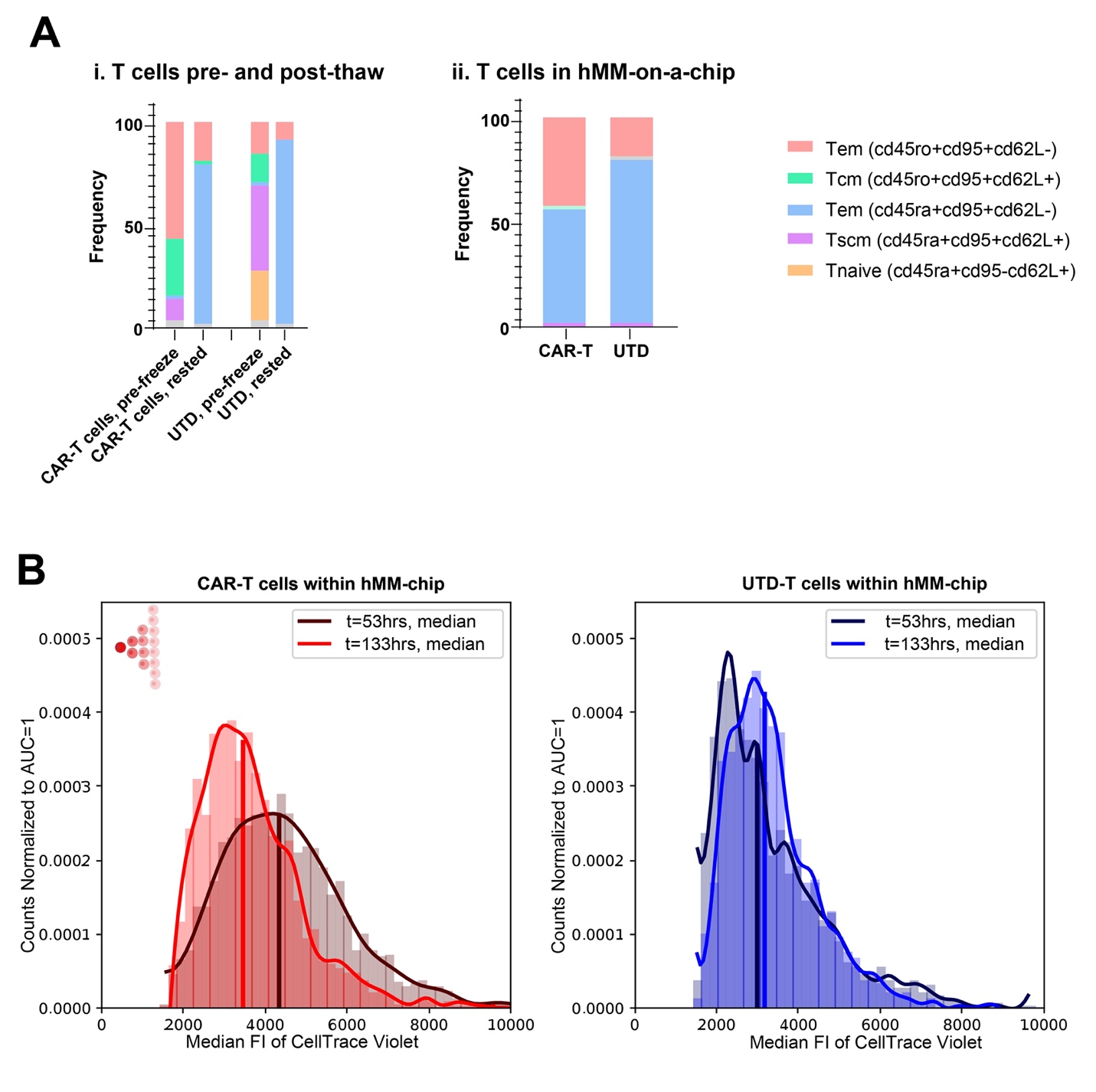
Fig. S12.

T cell characterization within hMM-on-a-chip treated with healthy-donor T cells. (A) Subsets of T cells (i) before and after cryopreservation, as well as (ii) relative abundances after endpoint within devices. (B) Comparison of MFI for individual labeled T cells at the initiation and termination of culture, to investigate whether the cells divided and lost their fluorescent signal (left inset schematic). n=3-4 devices per condition.

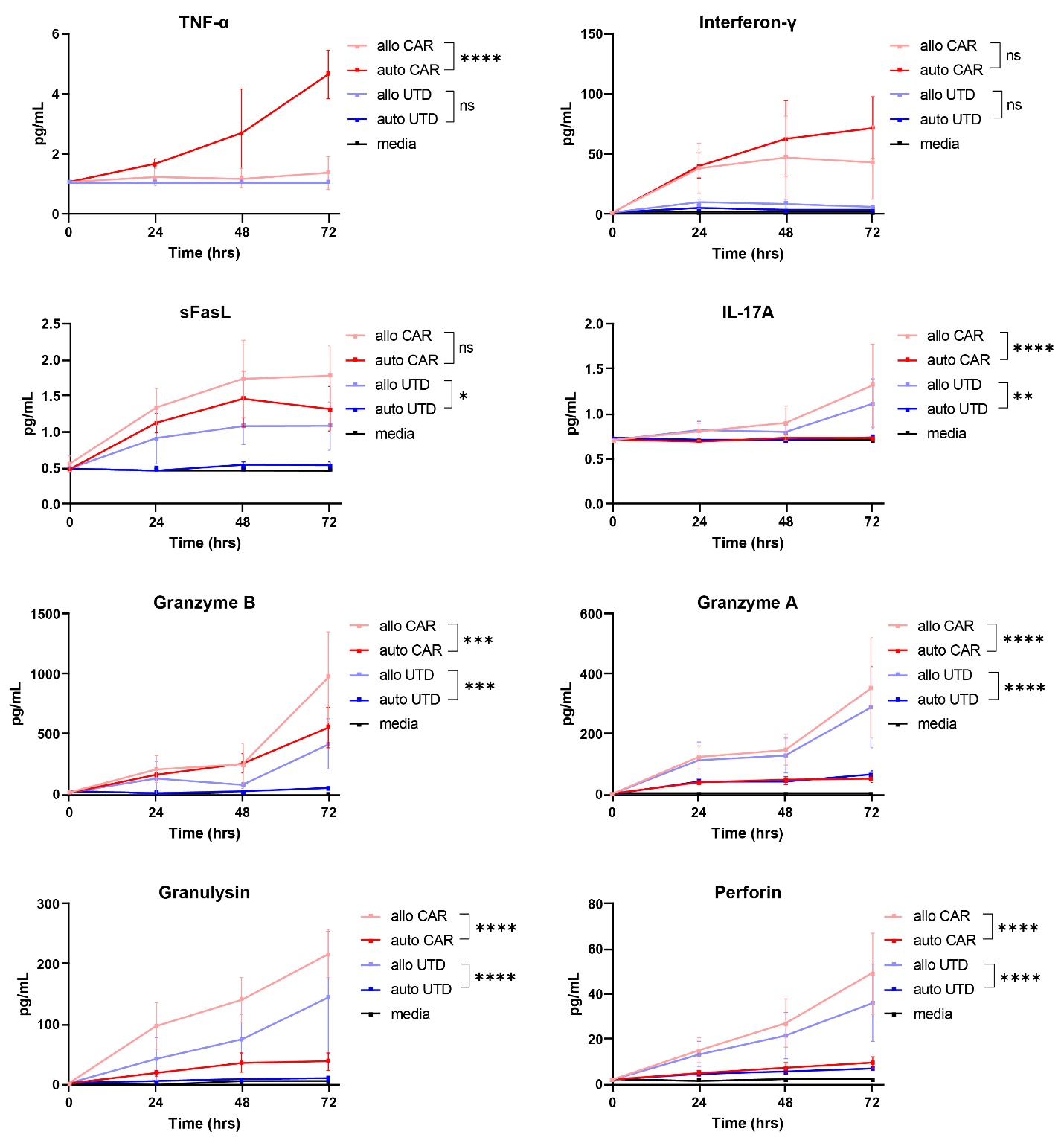
Fig. S13.

Graft-versus-host cytokine release within hMM-on-a-chip. Raw concentrations of cytokines tracked over time between allogeneic and autologously-introduced CAR- and UTD-treated devices. 2-way repeated measures ANOVA with Tukey post-hoc test, with stars indicating significance of final timepoint between on- and off-target cells. * = p ≤ 0.05, ** = p ≤ 0.01, *** = p ≤ 0.001, **** = p ≤ 0.0001.

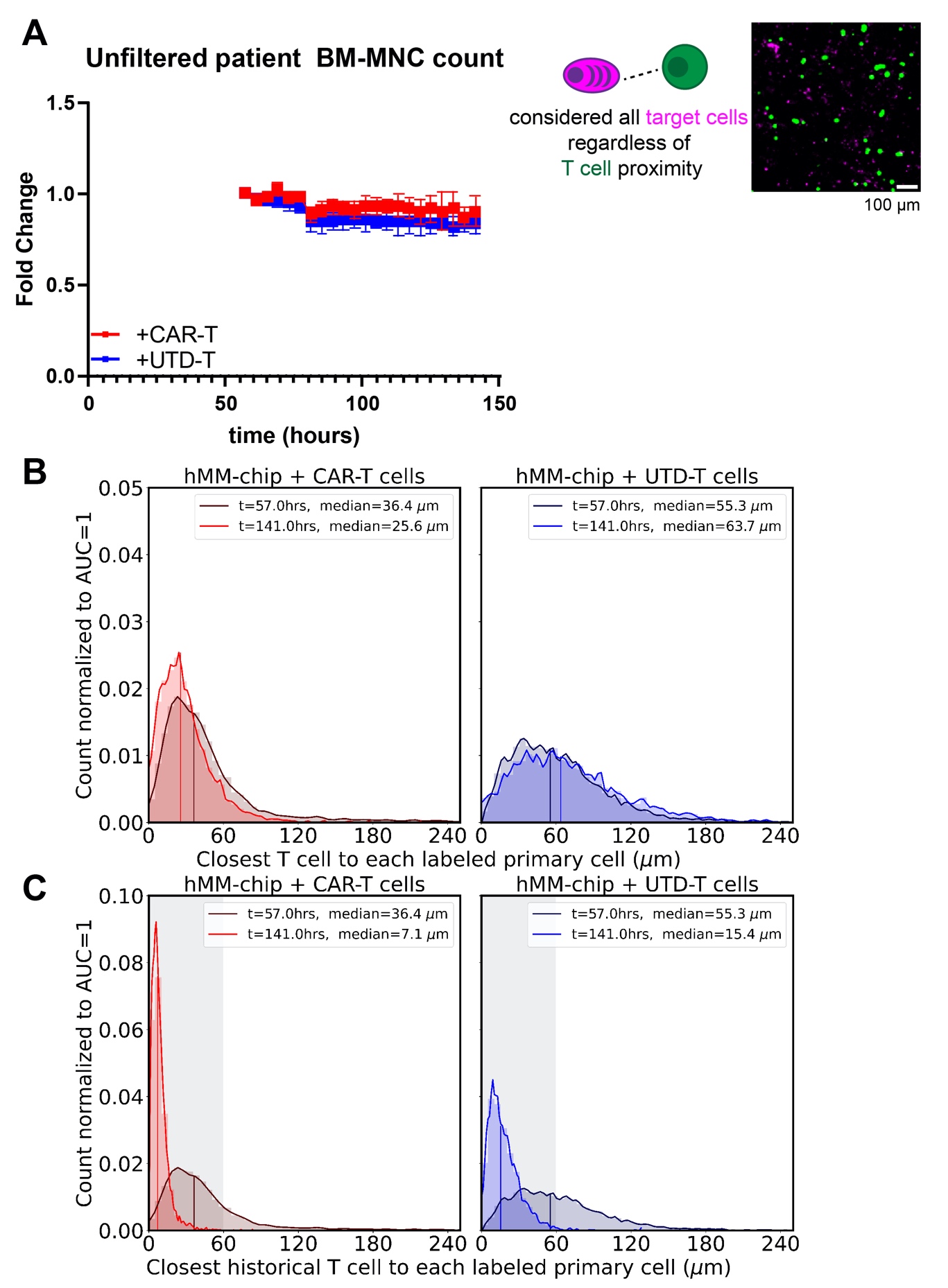

Fig. S14.

Justification for distance filtering. (A) After CellTrace Violet- labeled CAR-T cells (red trace) or UTD -T cells (blue trace) were introduced into the hMM-on-a-chip, the number of red-fluorescent cell objects did not significantly differ between effector conditions without distance filtration of the data. 2-way repeated-measures ANOVA, n=3-4 devices per condition. (B) Normalized histogram of the nearest T cell distances for each labeled primary cell for each timepoint. (C) Normalized histogram of the nearest historical T cell distances for each labeled primary cell for all cumulative timepoints. Shaded box indicates the 60 µm threshold applied to data depicted in the main figure.

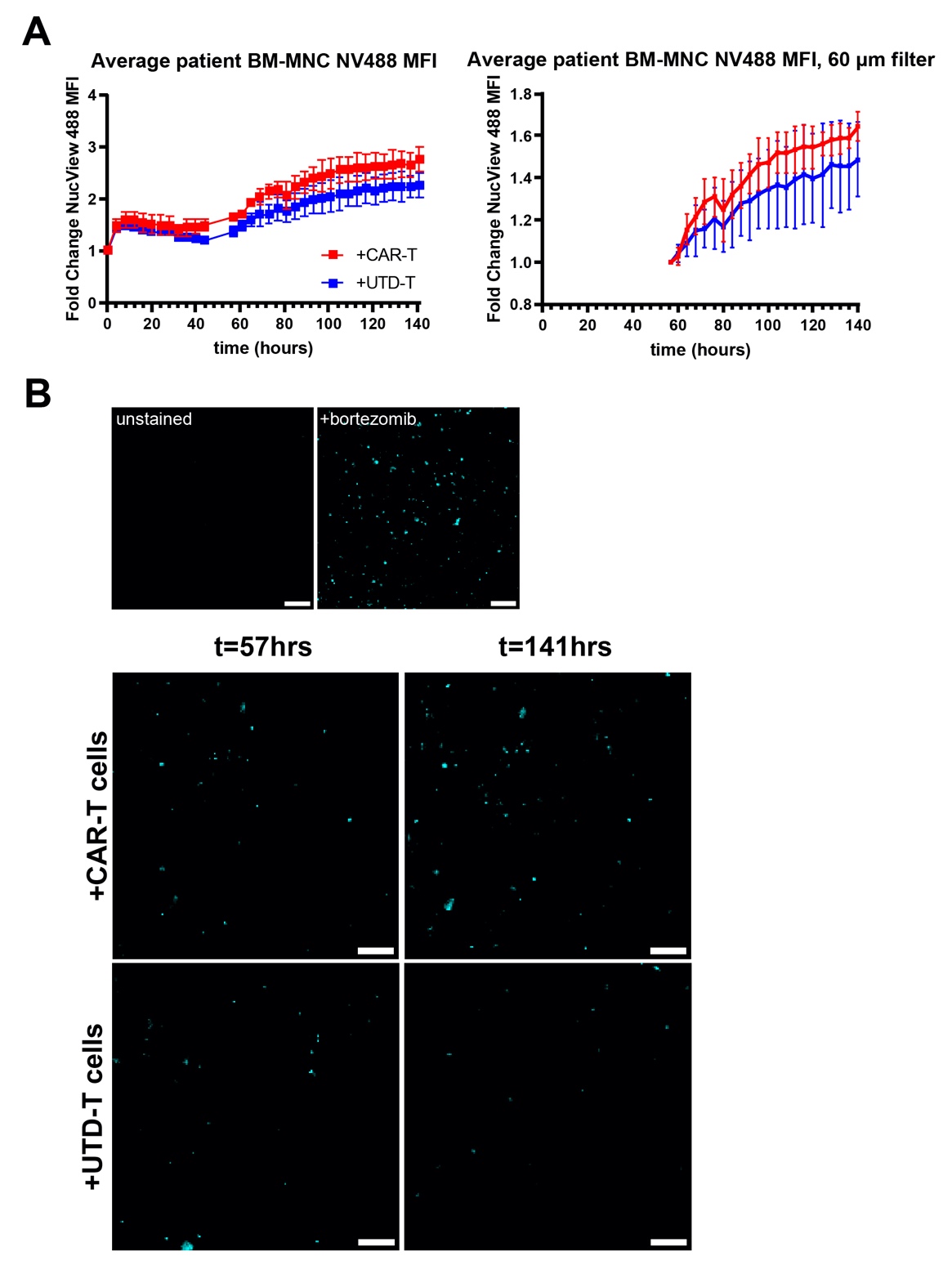

Fig. S15.

Results from using real-time caspase-sensitive dye to track target cell death. (A) After CellTrace Violet-labeled CAR-T cells (red trace) or UTD-T cells (blue trace) were introduced into the hMM-on-a-chip about 50 hours after initiation of culture within the BioSpa, the MFI of NucView 488 within labeled target cell objects did not significantly differ between effector conditions for either (i) raw data or (ii) distance-filtered data. n=3-4 devices per condition. (B) Representative images of NucView 488 data that was quantified in part (A). Top: signal from unstained device and device treated with bortezomib, an apoptosis inducer. Bottom: NucView 488 signal at beginning of T cell culture (t=57 hours) and at culture endpoint (t=141hrs)

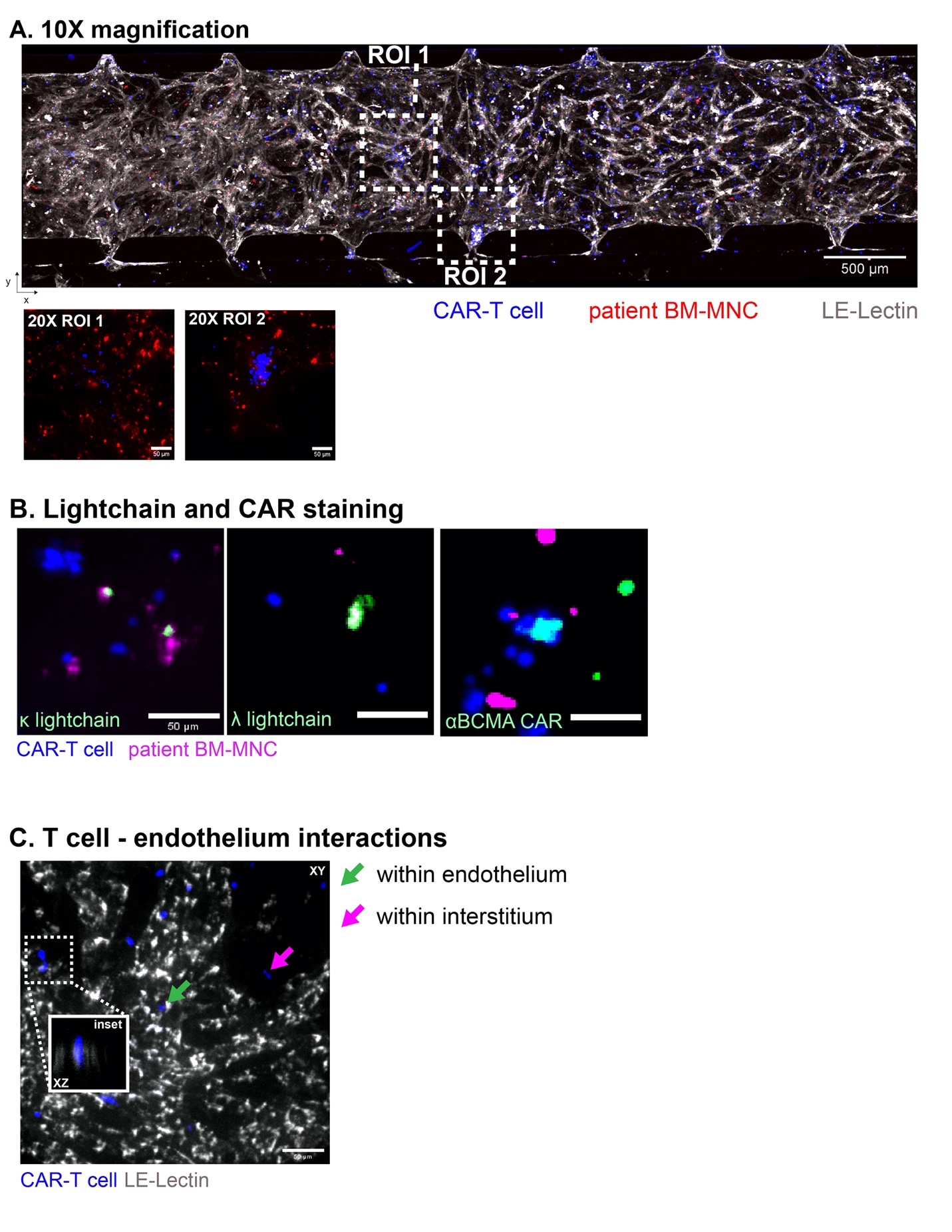
Fig. S16.

Microscopy of T cell and MM cell markers within hMM-on-a-chip. (A) CAR-T cells could be seen in 10X entering through each pillar opening and mostly remaining within the boundaries of lectin-labeled endothelium. When 20X images were captured (insets, lectin signal omitted for ease of viewing). (B) Labeled CAR-T cells (blue) interacting with labeled BM-MNCs (magenta) stained for κ and λ light-chain+ plasma cells (left, middle) or the anti-BCMA CAR molecule on the T cell (right). (C) T cells flow into vasculature and home to various regions within the vessels or in the interstitial space. They can travel from within the vessels to the interstitium by way of extravasation through the endothelium wall (20X inset).

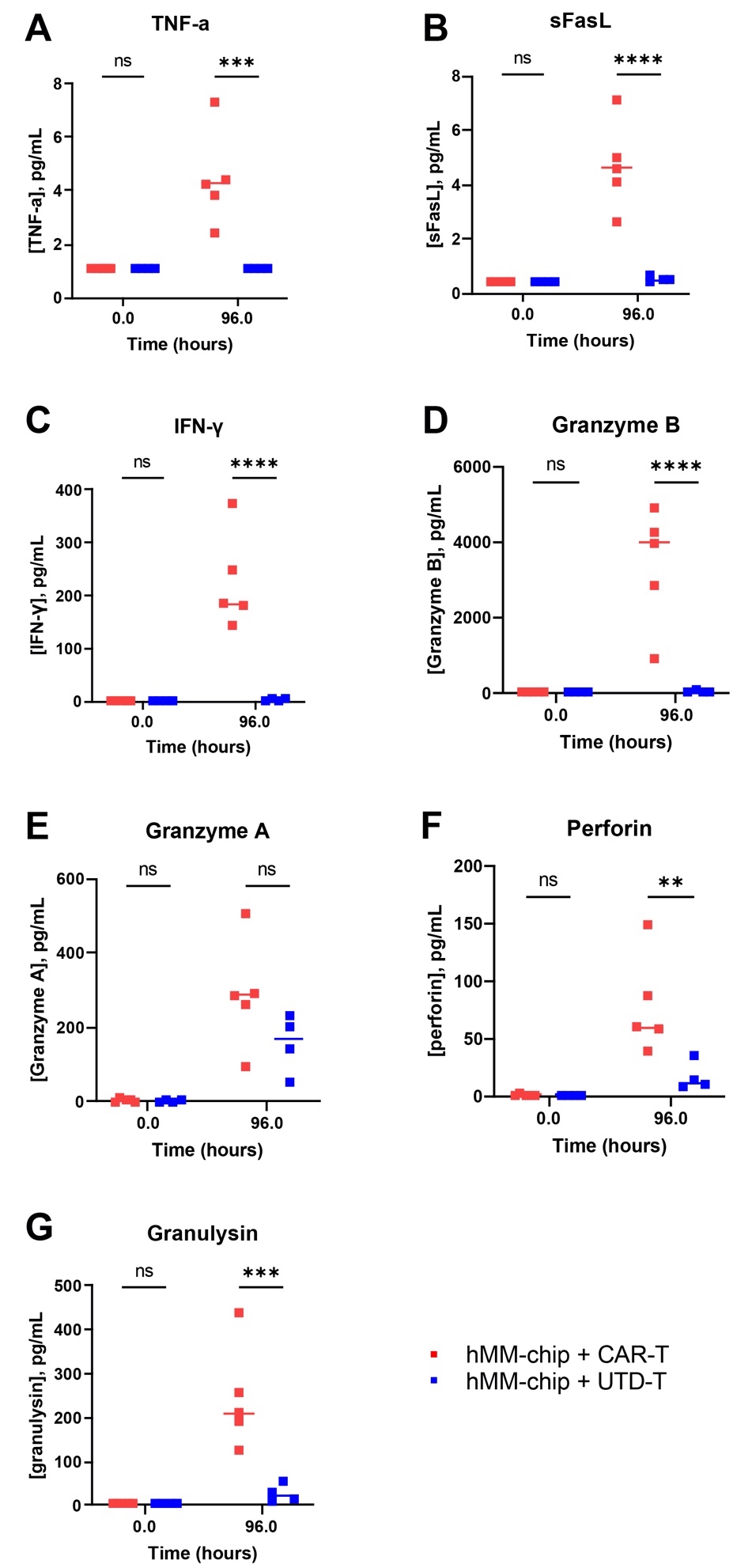

Fig. S17.

Raw cytokine release values for hMM-on-a-chip with primary BM-MNC targets. Welch's non-parametric t-test with Dunnett T3 post-hoc tests, ** = p ≤ 0.01, *** = p ≤ 0.001, **** = p ≤ 0.0001. The CAR-T-specific cytokines, as before, were (A) TNF-α, (B) sFasL, (C) IFN-γ, and (D) Granzyme B. Non-specific cytokines released by both cell types were (E) Granzyme A, (F) Perforin, and (G) Granzyme A.

Table S1.

Device fabrication materials list.

| **Material** | **Vendor** | **Catalog** | **Notes** |
| --- | --- | --- | --- |
| Trichloro (1H,1H,2H,2H-perfluorooctyl) silane | Sigma-Aldrich | 448931 |  |
| 96-Well No-Bottom Plates | Greiner Bio-One | 655000-06 |  |
| 96-Well Plate Lids | Greiner Bio-One | 656170 |  |
| Biopsy Punches | Integra Miltex | 12-460-401 |  |
| Sylgard 184 Silicone Elastomer Kit | Dow Corning | 2065622 |  |
| (3-mercaptopropyl) trimethoxy silane | Sigma-Aldrich | 175617 |  |
| HT-6240 | Bisco | HT-6240 |  |
| Dopamine hydrochloride | Sigma-Aldrich | H8502 | dispose of as toxic waste |
| Rat Tail Collagen I | Corning | 354249 |  |

Table S2.

Cell culture reagent list.

| **Reagent** | **Vendor** | **Catalog** | **Notes** |
| --- | --- | --- | --- |
| HUVEC, pooled donors | Lonza | C2519A | used until passage 8 |
| EGM-2MV | Lonza | CC-3162 |  |
| gelatin from porcine skin | Sigma-Aldrich | G2500 |  |
| VEGF | PeproTech | 100-20 | used at 50 ng/mL |
| Ang-1 | PeproTech | 130-06 | used at 100 ng/mL |
| IL-2 | PeproTech | 200-02 | used at 100U/mL |
| IL-6 | PeproTech | 200-06 | used at 10 ng/mL |
| NHLF, donor 615568 | Lonza | CC-2512 | used until passage 8 |
| FGM | Lonza | CC-3132 |  |
| MSC, donor 183 | RoosterBio | MSC-001 | used until passage 6 |
| α-MEM | Sigma-Aldrich | M0644 |  |
| sodium bicarbonate | Gibco | 25080094 |  |
| β-glycerophosphate | Sigma-Aldrich | G9422 | used at 10 mM |
| L-ascorbic acid | Sigma-Aldrich | 255564 | used at 50 µM |
| dexamethasone | Sigma-Aldrich | D2915 | used at 100 nM |
| CAG-RFP lentiviral vector | Cellomics Technology LLC | PLV10071 | used at MOI *≈*5 |
| polybrene | Sigma-Aldrich | TR-1003-G | used at 8 µg/mL |
| puromycin dihydrochloride | Gibco | A1113803 | used at 0.5µg/mL |
| RPMI | Gibco | 11875093 |  |
| penicillin-streptomycin (PS) | HyClone | SV30010 | used at 1% |
| gentamicin | Sigma-Aldrich | G1397 | used at 50 µg/mL |
| antibiotic/antimycotic | Sigma-Aldrich | A5955 | used at 1% |
| fetal bovine serum (FBS) | various | various | used at 10% |
| dimethylsulfoxide (DMSO) | Sigma-Aldrich | D2438 | used at 10% |
| CryoStor 10 | BioLife Solutions | 210102 |  |
| Histopaque-1077 | Sigma-Aldrich | 10771 |  |
| RBC lysis buffer | Tonbo Biosciences | TNB-4300-L100 |  |
| RO4929097 | SelleckChem | S1575 | use at 100nM |

Table S3.

IHC antibody list.

| **Antigen or Reagent** | **Clone** | **Vendor** | **Catalog** | **Isotype** | **Concentration Used** |
| --- | --- | --- | --- | --- | --- |
| Alizarin Red S | NA | Sigma-Aldrich | A5533 | NA | 2% |
| Collagen I | 5D8-G9 | Invitrogen | MA1-141 | Mouse IgG1 | 1.2 µg/device |
| Collagen IV | 1042 | eBioscience | 53-9871-82 | Mouse IgG2b, κ | 0.15 µg/device |
| Collagen IV | 1042 | eBioscience | 51-9871-82 | Mouse IgG2b, κ | 0.15 µg/device |
| Fibronectin | FN-3 | eBioscience | 14-9869-82 | Mouse IgG1 | 0.06 µg/device |
| CD90 | F15-42-1 | Invitrogen | MA5-16671 | Mouse IgG1 | 0.6 µg/device |
| Osteocalcin | OC4-30 | Invitrogen | 33-5400 | Mouse IgG2a | 1.2 µg/device |
| Osteopontin | 7C51112 | Invitrogen | MA5-17180 | Mouse IgG1 | 1.2 µg/device |
| SCF | Polyclonal | Bioss | bs-0545R-A555 | Rabbit | 1.2 µg/device |
| Hyaluronan | NA | AMSbio | AMK-HKD-BC41 | NA | 0.6 µg/device |
| Phalloidin | NA | Biotium | 00040 | NA | 0.5U/device |
| Phalloidin | NA | Biotium | 00042 | NA | 0.5U/device |
| DAPI | NA | Cayman Chemical | 14285 | NA | 0.1 µg/device |
| αIgG1 | Polyclonal | Invitrogen | A21121 | Goat | 0.6 µg/device |
| αIgG2a | Polyclonal | Invitrogen | A21131 | Goat | 0.6 µg/device |
| αIgG2b | Polyclonal | Invitrogen | A21141 | Goat | 0.6 µg/device |
| Streptavidin | NA | Invitrogen | S32354 | NA | 0.6 µg/device |
| LEL | NA | Vector Labs | DL-1174 | NA | 0.25 µg/device |
| LEL | NA | Vector Labs | DL-1177 | NA | 0.25 µg/device |
| LEL | NA | Invitrogen | L32472 | NA | 0.25 µg/device |

Table S4.

Flow cytometry antibody list.

| **Antigen or Reagent** | **Clone** | **Vendor** | **Catalog** | **Isotype** | **Concentration per Test** |
| --- | --- | --- | --- | --- | --- |
| BCMA | 19F2 | BioLegend | 357520 | Mouse IgG2a, κ | 0.075 µg/test |
| CD3 | SK7 | BioLegend | 344811 | Mouse IgG1, κ | 0.05 µg/test |
| CD11b | ICRF44 | BioLegend | 301365 | Mouse IgG1, κ | 0.1 µg/test |
| CD14 | 63D3 | BioLegend | 367147 | Mouse IgG1, κ | 0.1 µg/test |
| CD19 | HIB19 | BioLegend | 302217 | Mouse IgG1, κ | 0.05 µg/test |
| CD31 | WM59 | BioLegend | 303112 | Mouse IgG1, κ | 0.05 µg/test |
| CD34 | 4H11 | eBioscience | 12-0349-41 | Mouse IgG1, κ | 0.05 µg/test |
| CD34 | 561 | BioLegend | 343615 | Mouse IgG2a, κ | 0.05 µg/test |
| CD38 | HIT2 | BioLegend | 303519 | Mouse IgG1, κ | 0.1 µg/test |
| CD45 | HI30 | BioLegend | 304043 | Mouse IgG1, κ | 0.05 µg/test |
| CD45 | HI30 | BioLegend | 304019 | Mouse IgG1, κ | 0.04 µg/test |
| CD56 | 5.1H11 | BioLegend | 362509 | Mouse IgG1, κ | 0.125 µg/test |
| CD90 | 5E.10 | BioLegend | 328107 | Mouse IgG1, κ | 0.1 µg/test |
| CD138 | DL-101 | BioLegend | 352324 | Mouse IgG1, κ | 0.125 µg/test |
| CD138 | MI15 | BioLegend | 356523 | Mouse IgG1, κ | 0.05 µg/test |
| CD235a | 10F7MN | eBioscience | 367-9886-42 | Mouse IgG1, κ | 0.05 µg/test |
| Isotype Control | MOPC-173 | BioLegend | 400259 | Mouse IgG2a, κ | equal to test antibody |
| CD45RO | UCHL1 | BD Horizon | 564292 | Mouse IgG1, κ | 0.1 µg/test |
| CD62L | DREG-56 | BioLegend | 304825 | Mouse IgG1, κ | 0.125 µg/test |
| hBCMA | NA | AcroBio | BCA-HF254 | NA | 0.333 µg/test |
| CD8 | SK1 | BioLegend | 344703 | Mouse IgG1, κ | 0.05 µg/test |
| CD95 | DX2 | BioLegend | 305621 | Mouse IgG1, κ | 0.04 µg/test |
| CD45RA | HI100 | BioLegend | 304119 | Mouse IgG1, κ | 0.125 µg/test |
| CD3 | SK7 | BioLegend | 344839 | Mouse IgG1, κ | 0.04 µg/test |
| Human TruStain FcX | NA | BioLegend | 422302 | NA | 2 µL/test |
